# Proteome Remodeling of the Eye Lens at 50 Years Identified with Data-Independent Acquisition

**DOI:** 10.1101/2022.05.06.490936

**Authors:** Lee S. Cantrell, Kevin L. Schey

## Abstract

The eye lens is responsible for focusing and transmitting light to the retina. The lens does this in the absence of organelles yet maintains transparency for at least five decades before onset of age-related nuclear cataract (ARNC). It is hypothesized that oxidative stress contributes significantly to ARNC formation. It is additionally hypothesized that transparency is maintained by a microcirculation system (MCS) that delivers antioxidants to the lens nucleus and exports small molecule waste. Common data-ependent acquisition (DDA) methods are hindered by dynamic range of lens protein expression and provide limited context to age-related changes in the lens. In this study we utilized data-independent acquisition (DIA) mass spectrometry to analyze the urea insoluble, membrane protein fractions of 16 human lenses subdivided into three spatially distinct lens regions to characterize age-related changes, particularly concerning the lens MCS and oxidative stress response. In this pilot cohort, we measured 4,788 distinct protein groups, 46,681 peptides, and 7,592 deamidated sequences, more than in any previous human lens DDA approach. Our results reveal age-related changes previously known in lens biology and expand on these findings, taking advantage of the rich dataset afforded by DIA. Principally, we demonstrate that a significant proteome remodeling event occurs at approximately 50 years of age, resulting in metabolic preference for anaerobic glycolysis established with organelle degradation, decreased abundance of protein networks involved in calcium-dependent cell-cell contacts while retaining networks related to oxidative stress response. Further, we identified multiple antioxidant transporter proteins not previously detected in the human lens and describe their spatiotemporal and age-related abundance changes. Finally, we demonstrate that aquaporin-5, among other proteins, is modified with age by PTMs including deamidation and truncation. We suggest that the continued accumulation of each of these age-related outcomes in proteome remodeling contribute to decreased fiber cell permeability and result in ARNC formation.

## Introduction

The ocular lens is a transparent tissue lacking vasculature and is responsible for light transmission to the retina for visual perception (1). The lens originates from primary fiber cells that differentiate from epithelial cells *in utero*. Throughout life, concentric growth rings of secondary fiber cells are added to the lens, differentiating at the lens equator from an anterior monolayer of epithelial cells to elongated fiber cells that extend toward the anterior and posterior poles of the lens (Figure 1A). Unlike most cell types, lens fiber cells are not degraded throughout life but experience organelle degradation and enter a senescent-like state shortly after elongation as part of cellular maturation (2). Thus, proteins in the center of the lens are effectively as old as the subject and proteins are spatially organized, as are fiber cells, in concentric growth rings. Unlike other cell systems, cellular stress response cannot be managed by transcriptional control, and must be accommodated by long-lived lens proteins and their small molecule interacting partners.

**Figure 1.**
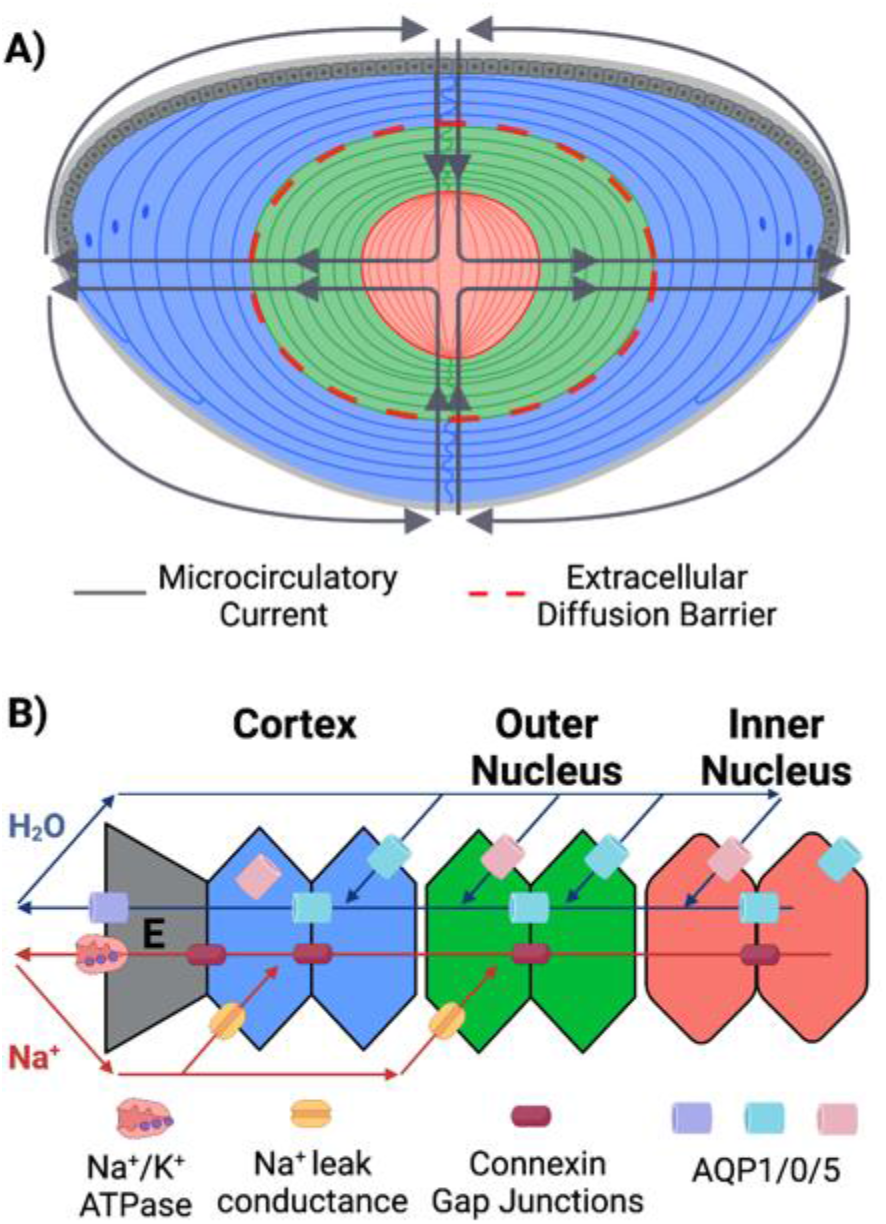
Cartoons of the lens and microcirculatory system: A) Cartoon of the lens with fiber cells divided into cortex (blue), outer nucleus (green) and inner nucleus (red) with the net convection of the microcirculatory system displayed. The approximate positioning of the extracellular diffusion barrier is noted within the inner nucleus. B) Cartoon of the electromotive potential establishment and net current in the cross section of cells in the microcirculatory system. Na/K ATPases at the epithelium (E) transport sodium from the lens, with re-uptake enabled by sodium leak conductance channels in fiber cells. Sodium transport is enabled by aquaporin-0 and −5 in fiber cells and aquaporin-1 in epithelial cells. The current established by microcirculation allows small molecule metabolites to transport intercellularly through connexin gap junctions at the cross section of fiber cells. Figure adapted from Schey et al. 2017.

Lens protein oxidation, particularly in the lens inner nucleus is hypothesized to contribute to formation of age-related nuclear cataract (ARNC)(3, 4). ARNC is the most common form of blindness and pharmacological options for delaying onset, prevention, or reversal are lacking. In a young lens, oxidative stress response is primarily mediated by glutathione-requisite proteins (5–10). In the absence of vasculature, small molecules including water and glutathione (GSH) must be delivered to the inner nucleus of the lens through the extracellular space of the lens sutures, established at the anterior and posterior poles of the lens between the tips of fiber cells (8).

The delivery of antioxidants and efflux of waste products is hypothesized to be established by the lens microcirculatory system (MCS)(11). In the MCS, small molecules are delivered through the lens sutures to the inner nucleus, taken up by fiber cells, and are exported through the equator by a concerted network of intercellular junctions (connexins) and water channels (12). Exported molecules are then convected around the lens exterior towards the anterior and posterior poles before re-entering the lens. Molecular convection is established by an electromotive potential of Na/K ATPases transporting sodium out of the lens at the equator while concurrent export of potassium is mediated by potassium channels on the lens epithelium (13, 14). The resulting potential, coupled to aquaporin water transport and gap junction intercellular contacts leads to a net current of metabolite convection into the lens at the anterior and posterior poles (15, 16)(Figure 1B). Although unshown in Figure 1, it is also hypothesized that transient receptor potential cation channel subfamily V members 1 and 4 (TRPV1/4) mediate fiber cell osmolarity by regulation of Na/K ATPase activity and Aquaporin-5 trafficking to the plasma membrane (12, 17–19).

In the young lens, and especially in young cortical fiber cells, MCS functionality is sufficient to transport water and metabolites through the lens (20, 21). However, convection of small molecules to the lens nucleus and through progressively aged lenses is inhibited by a barrier to diffusion (6, 20). This barrier is formed and becomes kinetically significant by 40-60 years of age (20). This diffusion barrier is thought to arise due to age-related changes in the concerted MCS protein network and reduction of extracellular space between aged fiber cells (11, 13, 22). After barrier formation, small molecules may enter the lens through the sutures but do not proceed through the restricted extracellular space at the same rate as in the young lens. Thus, as the lens ages and GSH is not delivered to the lens nucleus, proteins incur oxidative damage at an accelerated rate relative to the young lens (3, 4, 6, 21). Protein byproducts of oxidative damage may ultimately result in protein misfolding, cross-linking, aggregation, and light scattering ARNC (4, 23).

Proteostatic response to the accumulation of oxidative damage is solely mediated by the proteins present in fiber cells at the time of oxidative stress. For example, alpha-crystallin, an abundant lens protein and member of the small heat shock protein family, binds partially misfolded substrates to prevent further misfolding or aggregation (24). In addition to critical prevention of light scattering, crystallin-driven prevention of protein degradation establishes and sustains a gradient index of refraction in the lens wherein protein concentration is not homogenous throughout the lens, to reduce spherical aberration (25). Although age-related modifications to lens crystallins, such as abundance change, deamidation, and truncation have been identified, a substantial gap in our knowledge exists in understanding how the MCS changes at the molecular level as a function of age.

Measurement of the lens proteome has previously been achieved by measurement of whole lens lysates, imaging mass spectrometry, and spatially targeted approaches employing manual separation or laser capture microdissection prior to sample analysis (26–32). A limitation to all lens proteomics experiments is the high abundance of crystallin proteins, reported to constitute up to 90% of the lens proteome (33). Crystallins are functionally critical in the lens, as described above (24). While important, high crystallin abundance limits the depth of analysis enabled without crystallin depletion. In this study, we use Data-Independent Acquisition (DIA) mass spectrometry coupled to a membrane protein enrichment strategy to measure the membrane proteome of human lenses at different ages. To quantitatively measure spatiotemporal changes, we have divided the lens into three regions with distinct biological functions: the inner nucleus, which contains primary fiber cells formed *in utero* alongside the oldest secondary fiber cells of the lens, the outer nucleus, where fiber cells are fully matured and reside on the interior side of the diffusion barrier (Figure 1A); and the cortex, which is composed of the youngest mature secondary fiber cells, differentiating secondary fiber cells with organelles still intact and a monolayer of epithelial cells. Proteome measurement in each of these regions is critical for understanding how age and barrier establishment perturb protein machinery responsible for MCS establishment and delivery of antioxidants to the inner nucleus.

## Experimental Design and Statistical Rationale

Sixteen human lenses from age 15-74 years were analyzed with sample selection guided by known age-related lens physiology. Exclusion criteria included several cataract comorbidities such as diabetes mellitus, nicotine use, and non-age-related cataract. Additionally, no cataract lenses were used in this study. These studies were conducted in accordance with the ethical standards of the institutional research committee and with the 1964 Helsinki declaration and its later amendments. Each of the 16 lenses was divided into 3 regions (cortex, outer nucleus, and inner nucleus) to yield a total of 48 samples in the dataset. For sample quality control, 0.75 ug MassPREP protein standard was added to each approximately 75 ug lysate to monitor digest efficiency between replicates.

Samples were analyzed with Data-Independent Acquisition without pre-fractionation, with MS1 spectra interspersed every 30-31 scans. Each of the 48 samples analyzed in the study were used to generate a spectral library in DIA-NN (v1.8.0) (34) with an initial *in silico* library predicted by DIA-NN against a UniProt SwissProt canonical human fasta database with 9 MassPREP spike in proteins added (UP000005640, downloaded 10/12/2021, 20,402 entries). From this initial library, a subset of experimentally measured spectra were used to create an experiment specific library. After generation of the experimental spectral library, samples were re-analyzed, and proteins were quantified. No retention time standards were used, instead, endogenous peptide retention times were used for alignment in DIA-NN. Each sample was prepared and analyzed in an order determined by a random number generator and a blank gradient was run between samples to minimize carry over and quantitative bias. Trimmed mean of M-Values (TMM) normalization was employed for sample normalization. Sample groups were defined by exploratory data analysis that revealed clustering of lenses before and after 50 years of age. Within the young lens cohort, 9 lenses were analyzed (aged 15, 18, 22, 34, 34, 41, 44, 46, 49 years) and 7 lenses were measured in the old lens cohort (aged 53, 57, 63, 64, 65, 68, 74 years). Sample size of this pilot cohort was set at sixteen to demonstrate changes that occur in human lenses on a protein network level and protein abundance changes that occur over a gradient of human age. Statistical significance of change was assessed with 2-sample t-tests and PSEA-Quant operated in “labeled” analysis mode(35). Statistical power of 2-sample t-tests was calculated with RnaSeqSampleSize package in R (36).

## Urea Insoluble Protein Isolation

The severity of cataract present in each eye was visually evaluated prior to tissue processing to confirm that advanced cataractous lenses were not analyzed. Mild yellowing of the lens was allowed, but each measured lens retained transparency. In addition to sample evaluation, de-identified ophthalmic medical history was reviewed to confirm that no cataract had been diagnosed. Samples were prepared as previously described (37). Briefly, lenses were mounted with the equatorial axis parallel to a cryostat chuck before removal of the anterior and posterior poles of the lens yielding a 1.0 mm thick equatorial lens section. Concentric biopsy centerpunches were taken at 4.5- and 7-mm diameter to yield inner nucleus (0-4.5 mm), outer nucleus (4.5 – 7.0 mm) and cortex (7.0 – 9.2 mm) samples. Tissue was hand homogenized in buffer containing 25 mM Tris (pH 8), 5 mM EDTA, 1 mM DTT, 150 mM NaCl, 1 mM PMSF. After homogenization, samples were centrifuged at 100,000g for 30 minutes and the supernatant discarded. Pellets were washed twice with the above homogenization buffer followed by washes with 3.5 M and 7 M urea added to the homogenization buffer. Centrifugation at 100,000g for 30 minutes was performed to separate the supernatant and pellets for each urea wash. The remaining urea insoluble pellet was taken up in 50 mM TEAB with 5% SDS and protein concentration was measured with a BCA assay.

Membrane pellets of urea insoluble sample (75 μg total protein) were suspended in 50 mM TEAB with 5% SDS. Spike-in of 0.75 ug MassPREP protein standard was added before DTT was added to 10 mM before incubation at 56°C for 1 hour to reduce disulfide bonds. Reduced cysteines were alkylated by adding IAA to 20 mM and incubating in the dark at room temperature for 30 minutes. Phosphoric acid was added to 2.5% to acidify proteins. Proteins were precipitated by addition of 6 working volume equivalents of cold 100 mM TEAB in 90% methanol before transferring protein to the S-Trap micro (Protifi). Samples were washed on the S-Trap with 100 mM TEAB in 90% methanol four times with centrifugation between steps to remove salts and detergents. Samples were then digested in the S-Trap with a 1:15 trypsin:protein ratio in 20 uL 50mM TEAB, pH 7.5 for 2 hours at 46°C. Digested peptides were eluted in 4 steps of 50 mM TEAB, 0.2% formic acid, 50 mM TEAB and 50% ACN. Eluted peptides were dried under vacuum centrifugation and taken up in 0.2% formic acid prior to data acquisition.

## Instrumentation and Data Analysis

Peptides were analyzed using a Dionex Ultimate 3000 UHPLC coupled to an Exploris 480 tandem mass spectrometer (Thermo Scientific, San Jose, CA). An in-house pulled capillary column was created from 75 μm inner diameter fused silica capillary packed with 1.9 μm ReproSil-Pur C18 beads (Dr. Maisch, Ammerbuch, Germany) to a length of 250 mm. Solvent A was 0.1% aqueous formic acid and solvent B was 0.1% formic acid in acetonitrile. Approximately 200 ng peptide was loaded and separated at a flow rate of 200 nL/min on a 95-minute gradient from 2 to 29% B, followed by a 14-minute washing gradient and 35-minute blank injection between runs. The exact gradient was determined by linearized separation of the top 50% most intense cortical peptide signals by the Gradient Optimization Analysis Tool (GOAT v1.0.1)(38).

For DIA, the Exploris 480 instrument was configured to acquire 61×20m/z (390-1010 m/z) precursor isolation window DIA spectra (30,000 resolution, AGC target 1e6, max IIT 55 msec, 27 NCE) using a staggered window pattern with window placements optimized by ThermoFisher XCalibur instrument controls. Precursor spectra (385-1015 m/z, 60,000 resolution, AGC target 3e6, max IIT 100 msec) were interspersed after each sequence of 30-31 MS/MS spectra of the mass range. Default charge state was set to +3, S-Lens RF level set at 40%, NCE set at 27 and data collected in profile mode. Each scan cycle of 63 spectra took approximately 4.4 seconds.

For analysis of DIA data, RAW files were converted to mzML files in MSConvert (39), with staggered window deconvolution performed to improve precursor specificity to a pseudo 10 m/z window width. Processed DIA files were searched in DIA-NN with an Intel Core i7-7700 CPU at 3.60 GHz utilizing 8 threads. For all searches, up to one missed trypsin cleavage was allowed on peptides 7-30 residues in length with N-terminal M excision and cysteine carbamidomethylation enabled. All fragments between m/z 200 and 1800 and in charge states +1-4 were considered. An initial spectral library was prepared by DIA-NN with deep learning-based spectra and retention time prediction against a UniProt SwissProt canonical human fasta database (UP000005640, downloaded 10/12/2021,20,402 entries) with a predicted trypsin/P protease used. In each search, the neural network classifier was run in double pass mode with likely interferences removed, quantitation was performed in Robust LC (high accuracy) mode and cross-run normalization was turned off. Two separate searches were performed, one without variable modification and one with up to one variable deamidation on asparagine or glutamine (+0.984016 Da). An initial search of all files produced a spectral library that was used to search the data a second time (termed match between runs). After a search of files against the experimental spectral library (7,504 proteins and 55,875 peptides in the library built without modifications and 5,659 proteins and 27,639 peptides in deamidation enabled library) precursors and protein groups were filtered at 1% FDR within DIA-NN.

## Statistical Analysis

Statistical analysis was initiated through custom R scripts on peptides having <1% q-value and <1% global protein q-value. Prior to protein assembly, all peptides within the MaxQuant contaminant list were removed (40). Proteins were only assembled on peptides considered proteotypic and abundances for peptides and proteins was calculated by the diann R package function diann_maxlfq (https://github.com/vdemichev/diann-rpackage). Because lens samples were not of high (>90%) quantitative similarity by pearson correlation of DIA-NN normalized protein abundances, normalization performed in DIA-NN was rejected for quantitation of peptides and proteins. To minimize the assumption employed by most normalization algorithms that all compared samples are congruent, a subset list of peptides and proteins detected in all samples was selected and TMM normalization was applied (41). TMM applies a linear multiplier for normalization, which was extracted and uniformly applied to all rows of the peptide and protein matrices, including rows where missing values were present. Subsequent comparisons between lenses of different age are indicative of the representative contribution of a protein to the urea insoluble lens proteome. The distribution of peptides and proteins was qualitatively assessed to ensure that TMM treatment produced similar abundance distributions between samples (Supplemental Figure S1). DIA analysis assumes that all measurable proteins are detected, especially when the match between runs mode of DIA-NN is employed; thus, no missing value imputation was used.

Statistical significance between young and old lenses was calculated with a simple parametric 2-sample t-test to minimize overfitting of data that had been log2 treated. Significance cutoff was set to a p-value of 0.01 for hierarchical clustering and volcano plot visualizations. Volcano plot significance was further established at 1.5 log2 fold change. Ontology significance was not calculated on proteins determined as statistically differentiated by hierarchical clustering or volcano plot analysis. Instead, PSEA-Quant was used to evaluate significance on an ontology level without arbitrary significance cutoffs (35). PSEA-Quant was implemented as recommended by the authors with enrichment between sample groups considered instead of enrichment within a single sample. Default parameters for PSEA-Quant were used, with iterations increased to 1,000,000 to enable empirical p-value assignment as low as 1/1,000,000.

## Data Visualization and Presentation

All data visualizations were produced in R with either the ggplot2, heatmap.2 or EnhancedVolcano packages. Principal component analysis (PCA) from the prcomp package was used to separate proteins based on relative protein abundance for all proteins without missing values in the compared dataset (number of proteins used in caption of each PCA figure). PCA processing included scaling of non-log2 transformed data. For hierarchical clustering, a Ward based approach was used (base R stats) to produce the dendrogram of proteins below the 0.01 p-value threshold (number of proteins in caption). Subsequent visualization was done with the heatmap.2 package. Equivalent statistics were used to plot the volcano plots with the EnhancedVolcano package. Proteins that were measured as significantly differentiated in volcano testing are listed in Supplemental Table S1.

For PSEA-Quant visualizations, the list of significant protein network ontologies exceeds visually interpretable space, so terms returned from PSEA-Quant search were manually filtered based on FDR and p-value to 0.1 and 0.01, respectively. The remaining terms were manually reduced to eliminate unspecific or redundant terms from the visualization. A full list of significant ontologies is available as Supplemental Tables S2-S7. Finally, selected protein abundance was visualized respective to age and lens region with corresponding linear trendlines calculated. For comparison of deamidation accumulation, the results from the DIA search with 1 variable deamidation were prepared as described and the ratio of deamidated peptide accumulation in each sample was calculated as deamidated peptide abundance / (deamidated + undeamidated abundances) to demonstrate the proportional accumulation of deamidation with age. Biological cartoons were drawn with BioRender.

## Quality Control

Samples were prepared and were ran in random order. A BSA standard was run every 8 samples to verify integrity of retention time, instrument calibration, and chromatographic similarity. Several samples were re-injected to match TIC intensity and subsequently control for variability in quantitation of peaks with different peak shapes. The initial injections of these samples were not used in spectral library development or statistical analysis. The MassPREP protein standard was spiked into protein lysates prior to digestion to monitor digestion efficiency by sequence coverage and Pearson correlation profile of peptide abundances. For t-testing, log2 treatment was performed to reduce non-biological variance.

## Results

### Data-independent acquisition of the lens proteome

To measure proteins associated with the membrane and intercellular interactions as key components of the lens MCS, we employed DIA on membrane and insoluble protein fractionated lens lysates from human lenses of different age. Fiber cells of increasing age were measured by separating the lens into three age-indexed regions: the cortex (youngest cells), outer nucleus, and inner nucleus (oldest cells). In total, 16 human lenses were measured (15-74 years old) resulting in 48 total samples. Each region described here approximately corresponds to growth regions previously defined by electron microscopy and MRI analysis of lens fiber cell morphology and physiology (11, 42). To evaluate the prevalence of deamidation, an age-related modification, we performed two distinct DIA data searches. The first was based on a spectral library with no variable modifications and the second was based on a spectral library with up to one variable deamidation. We used deamidation as a proxy for age-related accumulation of post-translational modifications that are less measurable with unenriched DIA approaches utilizing library-free search. In total, 4,788 distinct protein groups and 46,681 distinct peptides, not including modified sequences, were identified at 1% FDR. Additionally, 7,592 deamidated sequences were identified. In this study, DIA facilitated measurement of 2.5-fold more lens proteins than any previous human lens DDA method (37). Protein groups and peptides were normalized according to a TMM normalization strategy. The resulting data are evaluated as the representative contribution of each protein within the lens membrane or insoluble proteome. Global protein normalization was not performed to preserve the distribution of protein representation where we expect proteins to be progressively modified over time. Although lens proteins are not synthesized after fiber cell maturation, the accumulation or positive differential representation of proteins suggests a reduced rate of degradation of a protein relative to its constituent proteome or the accumulation of aggregated or other insolubilized cytosolic proteins.

### Determination of sample grouping by age

A primary goal of this study was to identify key subsets of protein networks that are preserved in the aging lens since they may relate to long-term maintenance of transparency. First, we performed PCA on proteins identified in all samples without imputation (n=884) (Figure 2A). Relative to conventional cell line model systems (43), few proteins were identified in all samples, which is reflective of long-lived protein degradation, modification, and of human biological variability. Throughout this study, imputation was not used to prevent invalid representation of protein abundance changes, especially in DIA analysis where it is believed that all peptides within the mass range and above the signal to noise threshold are measured (44). In total, 43% of variance was explained on principal component axis 1 (PC1), which showed a progressive transition of fiber cells from young cortical fiber cells to old inner nucleus fiber cells with some overlap of middle-aged fiber cell populations (e.g., old cortical, young inner nucleus and all outer nucleus fiber cells). Cortical lens region separation on PC1 was significantly explained by cytoskeletal elements periaxin, vimentin, and neurofilament medium polypeptide; cadherin junction proteins cadherin-2 and catenin alpha-2; and brain acid soluble protein 1 (BASP1). Consistent with lens proteome investigations, older nuclear fiber cells were most associated with crystallins γ-A/B/C/D and β-B1; crystallin species that are known to associate with the membrane with age (26, 32, 37, 45). Further supporting the biological integrity of normalization approach employed, intermediate filament proteins were enriched in the young, still maturing cortical fiber cells as previously reported (29).

**Figure 2.**
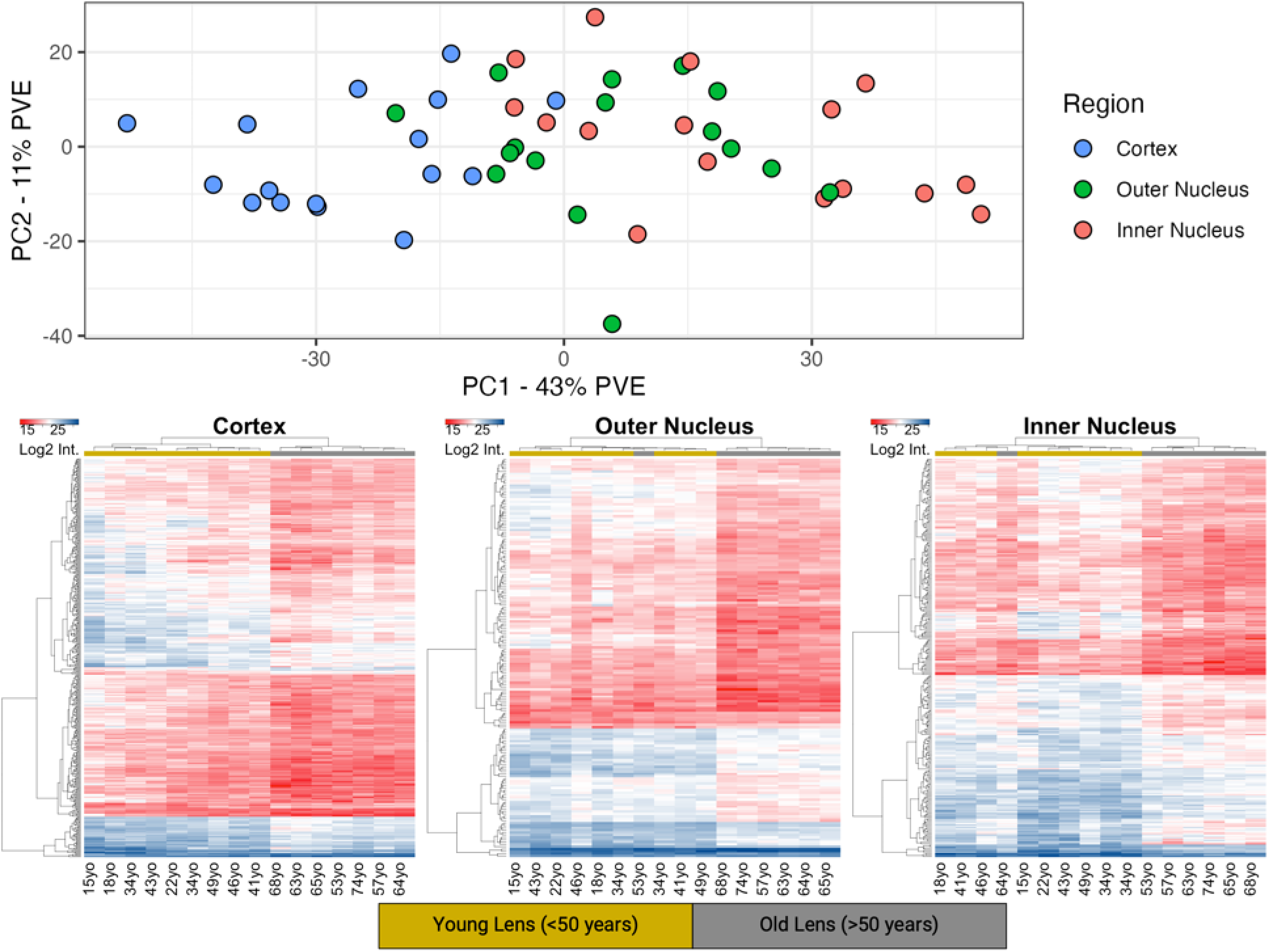
Determination of age groups in the human lens based on fiber cell positioning and age. A) Principal Component Analysis (PCA) on all 48 samples based on only protein groups measured in all samples (n = 884). Each point was colored according to lens region as in Figure 1. B) Cortex hierarchical clustering demonstrates clustering of proteins differentially present in samples above and below 50 years. Significance was measured by a t-test with a 0.01 significance cutoff (n=432). Columns colored by t-test sample grouping. Protein group names presented in Supplemental Table S1 if protein log2 fold change between groups exceeds 1.5. C) Outer Nucleus hierarchical clustering presented as in B (n=170). D) Inner Nucleus hierarchical clustering presented as in B (n=278).

From PCA analysis of all samples, no clear sample separation of subject age emerges. Age grouping was instead defined with hierarchical clustering on proteins with t-test significance at a 0.01 p-value between lenses below 50 years (young) and above 50 years (old). This preliminary cutoff was established congruent with the hypothesis of barrier formation at 45-50 years old (46). To evaluate consistency of biological changes directly associated with human age, each region of the lens was separately evaluated by hierarchical clustering (Figure 2B-D). As demonstrated in the dendrograms of each lens region, there is an appreciable change in the clustering of lenses before and after 50 years of age with single sample deviations for the 64-year-old inner nucleus and 53-year-old outer nucleus. To our knowledge, no prior dataset of the lens has demonstrated the clear separation of lenses observed at 50 years as shown here. Congruent clustering occurs without protein group filtering based on t-testing. These results suggest that a biological event occurs after approximately 50 years of aging that triggers a proteome remodeling event throughout the lens. As a result of these findings, we considered sample groups in the lens as young or old based on a 50-year cutoff.

### Age-related changes in the lens cortex

To demonstrate changes at the single-protein and protein-network level in the aging lens cortex, a three-step approach was taken. First, we performed PCA on all protein groups identified in each cortex sample. While the cortical data approximately clustered in the PCA plot shown in Figure 2A, clear separation of lens fibers from young and old lenses is demonstrated on the PC1 axis of this subset of samples (Figure 3A). Proteins that significantly contribute to the negative (younger) loading on PC1 include epoxide hydrolase, fibrillin-1, phosphate carrier protein, cytochrome c oxidase subunit 2, and tubulin beta-4A chain. Loadings that most significantly contribute to the positive (older) loading on PC1 include γB-crystallin, WD repeat-containing protein 25, 4-hydroxyphenylpyruvate dioxygenase, and glutamate synthesis enzyme kynurenine oxoglutarate transaminase. A volcano plot analysis was done to evaluate all proteins that change between young and older lenses (Figure 3B, filtered results available as Supplemental Table S1), with few proteins measured as increased in the old cortex relative to the young cortex. The statistical power of this test is estimated to be approximately 0.65 with a sample size of seven, the size of the old lens sample group, and a 5% FDR.

**Figure 3.**
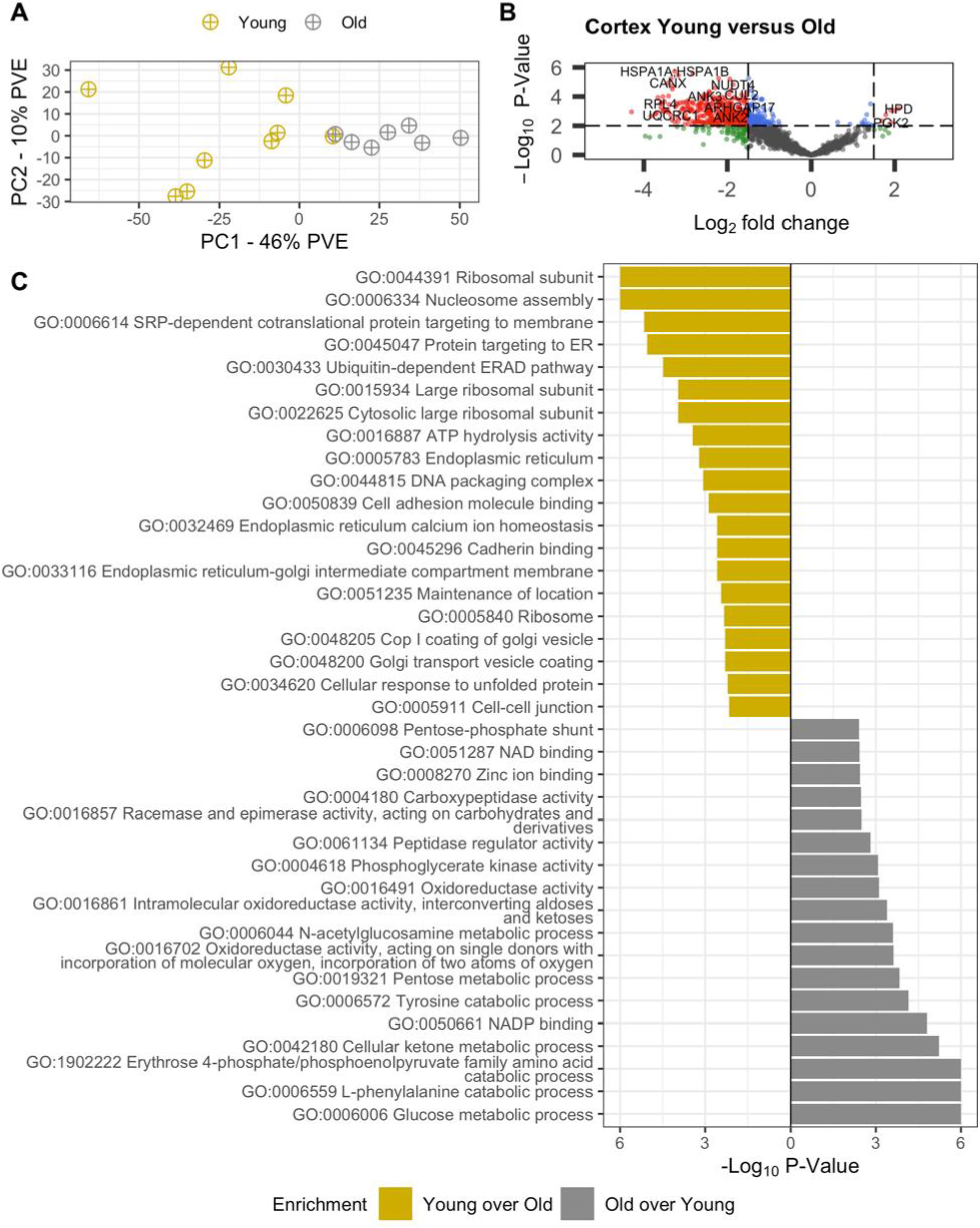
Graphical representation of age-related changes in the lens cortex region. A) PCA plot colored by age group demonstrates separation of young and old fiber cell populations on PC based on protein groups identified in all 16 samples (n=1,429). B) Volcano plot of preferentially retained or degraded proteins with significance cutoffs of 0.01 unmoderated p-value and 1.5 log2 fold change. UniProt identifiers converted to gene names. C) PSEA-Quant pairwise enrichment calculated between young and old lens fiber cell regions. Separate calculations performed to determine each enrichment. Significant Gene Ontology terms were filtered at 0.01 p-value and 0.1 FDR. Ontologies in graphic are a subset of all measured, demonstrating non-redundant daughter terms indicative of the complete enrichment set. Full list of enriched terms included in Supplemental Tables S2-S3.

To limit the effect of a relatively low statistical power in t-testing, PSEA-Quant was employed to identify protein networks that are most changed with age. Unlike t-tests, PSEA-Quant eliminates arbitrary p-value cutoffs by establishing enrichment scores as done in the Gene Set Enrichment Algorithm (47). The results file of significant GO terms was filtered at p <0.01 and <10% FDR (Figure 3C, Supplemental Tables S2,S3) consistent with PSEA-Quant developer suggested use (35). As expected, young lenses with intact organelle machinery still maintain ribosome, endoplasmic reticulum, Golgi, and other classical organelle structures (GO:0044391, GO:0048200, GO:0005783). While these organelle structures are present in both young and old lenses, the proportion of young and still maturing fiber cells is greater in the young lens than in the old lens. These young fiber cells also demonstrate canonical proteostasis enrichment relative to old fiber cells (GO:0030433, GO:0032469). Proteins involved in proteostasis include hsp70 BIP, hsp90 endoplasmin, wolframin, and DNAJB2. In the older cortical fiber cells, the results suggest that metabolic remodeling occurs as mitochondria are degraded, transitioning from oxidative phosphorylation to anaerobic glycolysis via the hexose monophosphate shunt pathway (GO:0051287, GO:0005911). It was also shown that cell-cell junctions are enriched in the young fiber cells relative to old and that oxidative stress response machinery is enriched in old fiber cells relative to young. Based on this result, we hypothesize that NADH production machinery is established in cortical fiber cells to metabolically support enzymes responsible for GSH-mediated oxidative stress response in aging fiber cells. Note that in-text graphics of PSEA-Quant results are a subset of all significant GO enrichment terms and were manually filtered by eliminating redundant or unspecific terms. The full list of enriched terms is available in Supplemental Tables S2 and S3.

### Age-related changes in the lens outer nucleus

For the outer nucleus region, the same approach was taken as was done for the cortex. PCA analysis (Figure 4A) demonstrates consistent separation of young lenses from old lenses as previously suggested. However, a single sample from the old cohort (53-years-old) did not separate according to the prescribed trend, as also demonstrated in hierarchical clustering (Figure 2C). Cell adhesion molecule 3, guanine nucleotide binding protein, insulin-like growth factor-binding protein 7, and BASP1 are protein groups associated with negative, young PC1 protein loadings. Protein phosphatase-1 regulatory subunit 15a, hspB3, βB1-crystallin and WD repeat-containing protein 25 are associated with positive, old PC1 protein loadings. No clear trends in biology emerge from these results aside from consistent young-lens loading annotation of BASP1 and old-lens βB1-crystallin and WD repeat-containing protein 25. There is no clear function for BASP1 or WD repeat-containing protein 25 in the lens. Outside of the lens, BASP1 interferes with MYC binding to calmodulin and its oncogenic capacity in cancer cell lines (48). WD repeat-containing protein 25 has diverse functionality outside of the lens, including RNA procession, signal transduction, vesicular trafficking, cytoskeletal assembly, and cell cycle control (49). It is unclear if these functions are operative in the lens, but maturing lenses must undergo extensive cytoskeletal remodeling as part of fiber cell elongation process (45, 50, 51).

**Figure 4.**
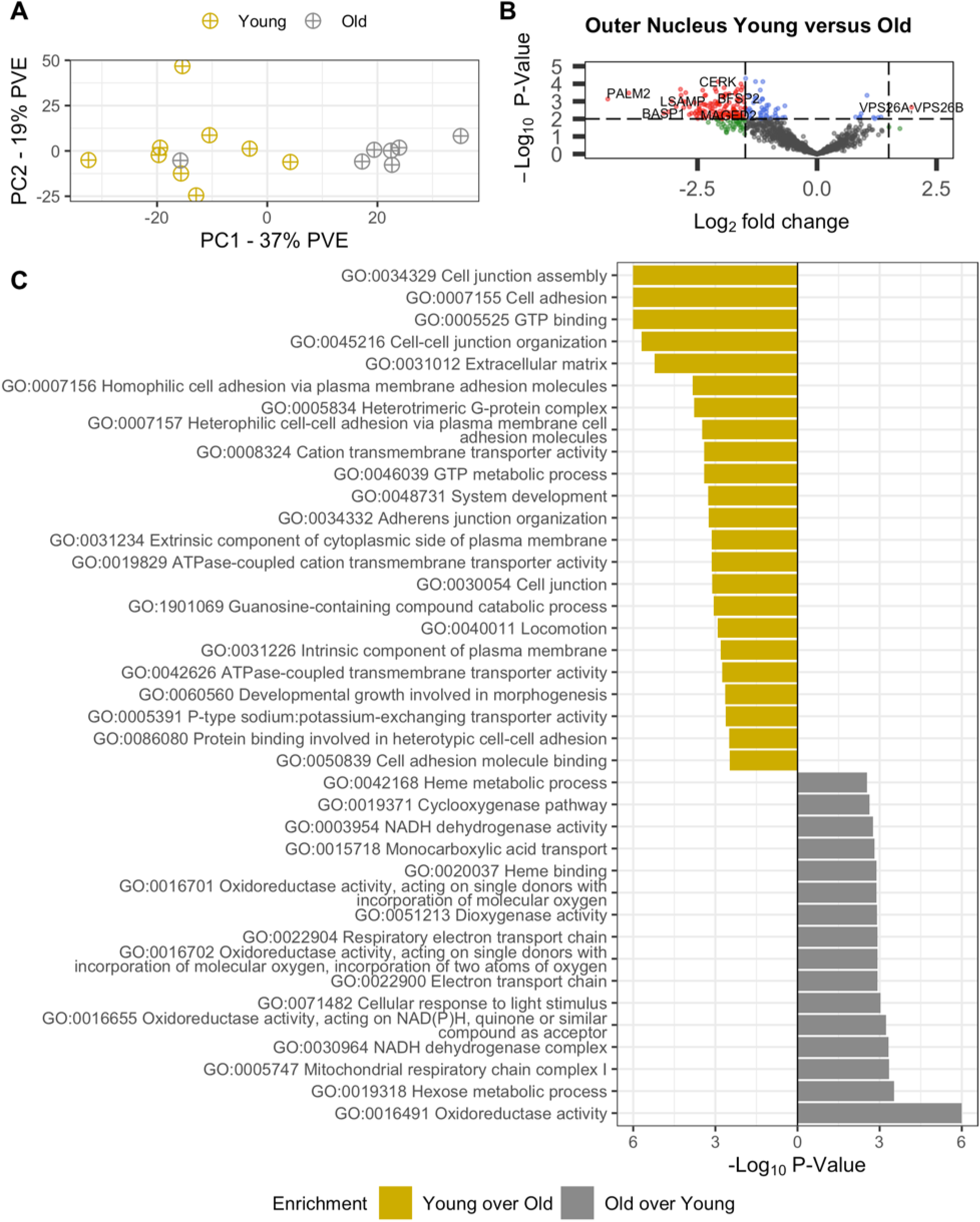
Graphical representation of age-related changes in the lens Outer Nucleus region. A) PCA plot colored by age group demonstrates separation of young and old fiber cell populations on PC based on protein groups identified in all 16 samples (n=1,114). B) Volcano plot of preferentially retained or degraded proteins with significance cutoffs of 0.01 unmoderated p-value and 1.5 log2 fold change. UniProt identifiers converted to gene names. C) PSEA-Quant pairwise enrichment calculated between young and old lens fiber cell regions. Separate calculations performed to determine each enrichment. Significant Gene Ontology terms were filtered at 0.01 p-value and 0.1 FDR. Ontologies in graphic are a subset of all measured, demonstrating non-redundant daughter terms indicative of the complete enrichment set. Full list of enriched terms included in Supplemental Tables S4-S5.

Two-sample t-testing of young and old lens outer nuclei did not yield as many significantly changed proteins as in the cortex; power estimated 0.6 at 5% FDR (Figure 4B, Supplemental Table S1). The decrease in power is attributed to fewer statistically different proteins (n = 178) and fewer proteins measured in all samples (n = 1,114). As done previously, this limitation was mitigated by implementation of PSEA-Quant in labeled mode (Figure 4C, Supplemental Table S4,5). In the outer nucleus, the prevailing theory is that anaerobic metabolism established in the cortex is less active and that remodeling of the proteome is exclusively attributed to age-related changes caused by intercellular signaling and oxidative homeostasis, not to transcriptional control. This is demonstrated by the enrichment of GO terms related to gap junction (GO:0034329) and cell-to-cell contacts (GO:0007156) in young lenses and enriched representation of oxidoreductase (GO:0016491, GO:0016655, GO:0016701) protein networks in old lenses relative to young lenses. Each of these trends are expected to be a continuation of the aging process observed in the cortex where transcription is eliminated. Specific analysis of proteins related to respiratory electron transport reveals most of these proteins (UniProt identifiers O96168, O43574, Q86Y39, O95298, Q9P0J0, O43181) are NADH dehydrogenases. It is expected that if these dehydrogenases are functionally active, they are not contributing to mitochondrial respiratory electron chain function because organelles are degraded in the outer nucleus.

### Age-related changes in the lens inner nucleus

The lens inner nucleus was evaluated as done for the cortex and outer nucleus regions. In PCA (Figure 5A), young and old lenses separated precisely at the 50-year age cutoff, in contrast to clustering results from hierarchical clustering (Figure 2D). PC1 protein loadings with the largest negative value include cell adhesion molecule 3, insulin-like growth factor binding protein 7, A-kinase anchor protein 2-related, and BASP1. Positive PC1 protein loadings included AP-1 complex subunit sigma 1a, protein phosphatase 1 regulatory subunit 15a, hspB3, βB1-crystallin, and vacuolar protein sorting-associated protein 26A. Trends here closely match those of the outer nucleus. Unlike previous volcano plot results, no proteins were significantly represented in the old lenses relative to young lenses (Figure 5B, Supplemental Table 1); 0.6 power at 5% FDR. The low number of statistically enriched old inner nucleus proteins relative to young inner nucleus abundances is likely due to to accumulation of age-related modifications including deamidation (27, 52–54). Deamidation is the most abundant non-isobaric irreversible PTM in the lens and randomly occurs on disordered region of client proteins, which is the region most likely to be measured in a membrane proteome measurement (28). Therefore, we examined the extent of deamidation as an explanation for decreased protein group abundance with age. We suggest that if the abundance of the unmodified peptide decreases and the deamidated peptide does not increase to a similar extent, that an alternative PTM is occurring, for example, truncation.

**Figure 5.**
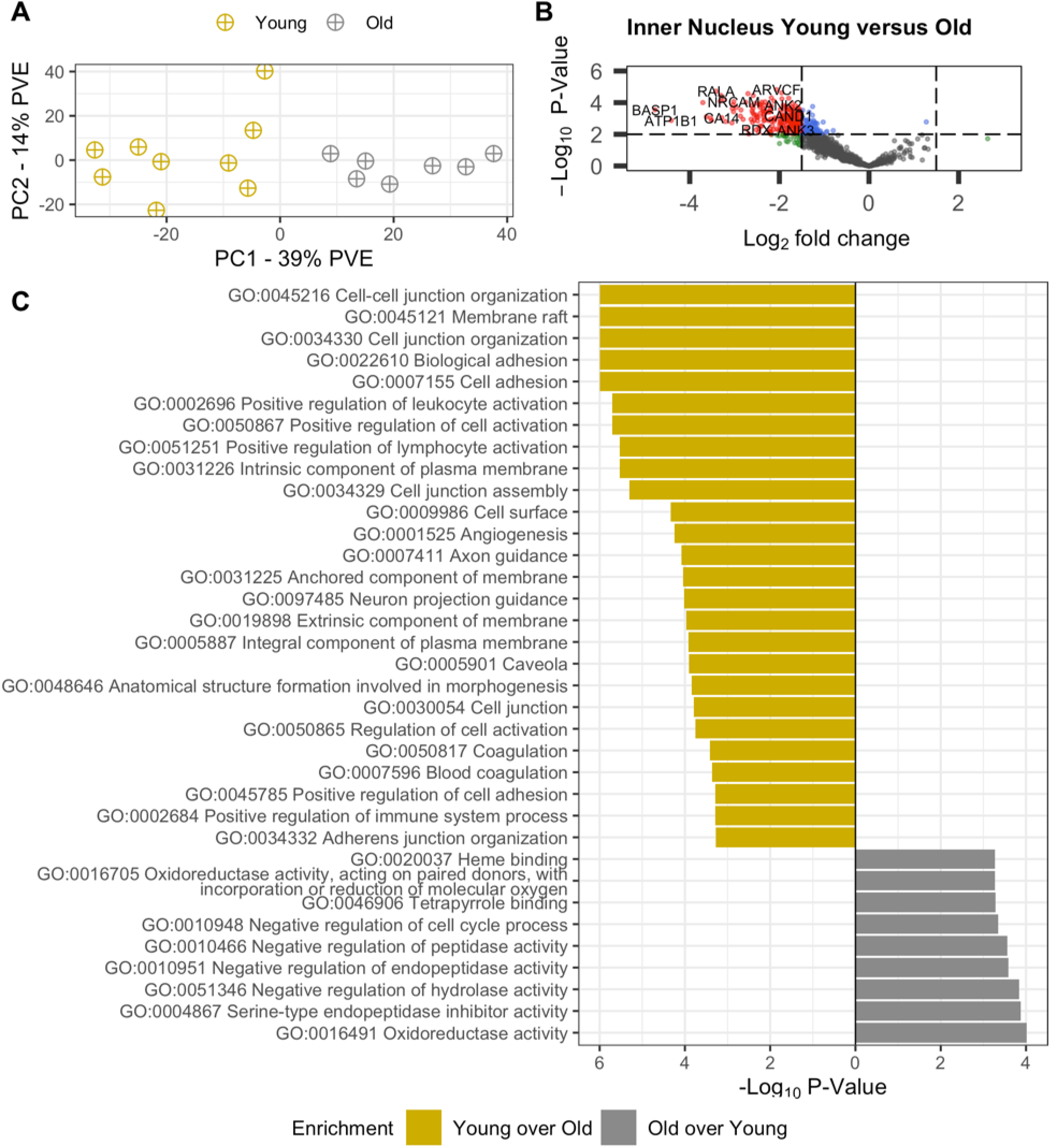
Graphical representation of age-related changes in the lens Inner Nucleus region. A) PCA plot colored by age group demonstrates separation of young and old fiber cell populations on PC based on protein groups identified in all 16 samples (n=969). B) Volcano plot of preferentially retained or degraded proteins with significance cutoffs of 0.01 unmoderated p-value and 1.5 log2 fold change. UniProt identifier converted to gene names. C) PSEA-Quant pairwise enrichment calculated between young and old lens fiber cell regions. Separate calculations performed to determine each enrichment. Significant Gene Ontology terms were filtered at 0.01 p-value and 0.1 FDR. Ontologies in graphic are a subset of all measured, demonstrating non-redundant daughter terms indicative of the complete enrichment set. Full list of enriched terms included in Supplemental Tables S6-S7.

To demonstrate the impact of deamidation with age, we plotted the age-related abundance of connexin 46 (GJA3) when no variable modifications were included in the spectral library alongside the abundance of deamidated peptide L10-K23 measured in a separate search where a single deamidation modification was enabled (Figure 6). Figure 6A demonstrates that while there is a modest decrease in the abundance of GJA3 in the inner nucleus with age (p <0.05). This may be partially explained by the accumulation of deamidation on the L10-K23 peptide (Figure 6C, proportional deamidation being the proportion of measured deamidated peptide divided by the sum of deamidated and unmodified peptide). When protein abundance change is calculated with deamidated and unmodified peptides combined, the statistical significance of abundance change is not observed (Supplemental Figure S2). The single variable deamidation dataset was not used for quantitative evaluation of all proteins because search engine stringency for deamidation decreases the number of proteins identified and deamidated or alternatively modified proteins may have decreased or otherwise unconventional protein function (53, 54). For all deamidated proteins discussed where more than one modified peptide was measured, the presented peptide proportional deamidation is congruent to unshown peptides.

**Figure 6.**
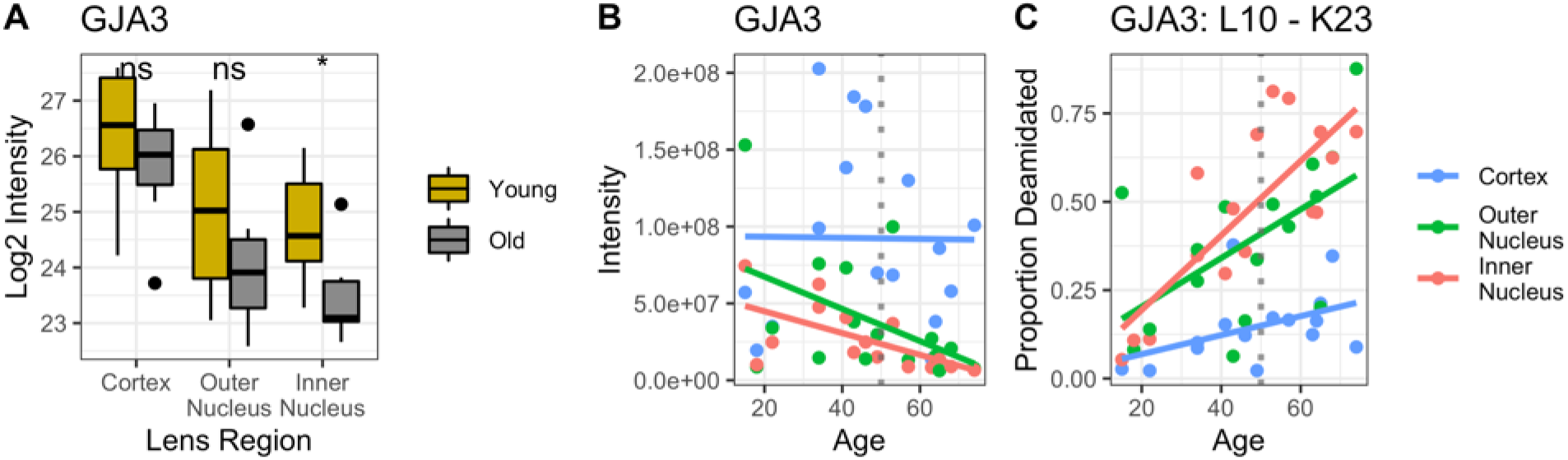
Representative abundance of Connexin 46 (GJA3) in each sample. A) Protein abundance of GJA3 in a search with no variable modifications and t-test significance of distributions (* = <0.05, ** = <0.01, *** = <0.001) between young and old lenses. B) Individual representative intensity of GJA3 signal in unmodified search, linear trendline plotted to demonstrate trend. C) Representative deamidation accumulation of GJA3 L10-K23 calculated on single variable deamidation dataset with linear trendline to show accumulation of modification.

In inner nucleus PSEA-Quant results (Figure 5C, Supplemental Table S6,7), oxidoreductase related terms (GO:0016491, GO:0016705) and negative regulation of several protein or peptide truncation ontologies (GO:0004867, GO:0010948) were among the few ontologies enriched in the old inner nucleus relative to the young inner nucleus. Peptide truncation ontologies were represented by protease inhibitors (UniProt identifiers P30740, Q92530, P04080, P35237, P50453) including leukocyte elastase inhibitor. The term “Heme binding” (GO:0020037) here corresponds to the enhanced representation of succinate dehydrogenase, peroxiredoxin-1, and 7-alpha-hydroxychloest-4-en-3-one 12-alpha-hydrolase. The consistent identification of enriched oxidoreductase activity in each old lens region relative to the young lens supports the hypothesis that a primary function in the aging lens is reduction of oxidative stress, and that this functionality is preserved by sustained protein abundances. Likewise, protein truncation ontology enrichment suggests that the lens attempts to preserve intact protein structure and function. More enriched GO terms were measured in the young inner nucleus relative to old, demonstrating age-related decrease of terms especially related to cellular adhesion and intercellular transport by gap junctions. A final observation from the PSEA-Quant results is the continued enrichment of cellular adhesion-related terms in young lenses relative to their old counterparts (GO:0045216, GO:0034330, GO:0007155, GO:0034329). The inferred depletion of these terms in old lenses demonstrates that age may be responsible for depletion of cellular adhesion and intercellular metabolite transport. Taken together, we propose that while putatively active, oxidative stress protein networks are inhibited by the decrease of cellular contacts and metabolite transport in the lens that accumulate with age.

## Discussion

In this study we have measured more proteins than previously detected in the human lens with DDA, exceeding all prior approaches in total protein groups identified by 2.5-fold (37). It should be emphasized that a protein not detected in this study is not indicative of protein absence from the lens. For example, while putatively involved in osmotic regulation of the lens and regulation of MCS current (18, 55, 56), neither TRPV1/4 were measured in any cortex sample. Immunohistochemistry in murine and porcine lenses supports the cortical presence of these proteins (55, 57, 58); however, no mass spectrometry reports support the presence of TRPV1/4 in the human lens cortex. This may be due to low abundance of TRPV1/4 and the expectation that the majority of each channel is expressed in the monolayer epithelium (58).

As a label-free analysis in samples with putatively high biological variability, it was important to confirm previously described lens biology in the present analysis. Previous studies comparing progressively aged fiber cells have been performed and are consistent with the results presented here and with the MCS hypothesis (26, 29). In addition to considering aging fiber cells, Truscott and colleagues previously evaluated human lens aging with isobaric tags (32). While those results are less rich than those afforded by DIA, several similarities can readily be made. In Truscott and the presented work, the measured abundance of β- and γ-crystallin variants was increased in progressively aged lenses (see supplemental data for protein group intensities). Each study showed no significant age-related change in the abundances of α-crystallin A and B subunits, AQP0. Similar distributions of connexin proteins were also observed in each study. Finally, each study demonstrates age-related decline in the abundance of BASP1, paralemmin 1, and vimentin. The lone disagreement between these studies is the inner nucleus measurement of GJA3: both studies suggest that the abundance is decreased significantly in older lenses, but we find that this change is insignificant when considering deamidation as a variable modification (Supplemental Figure S2). Taken together, these results suggest that the label-free DIA analysis employed here is sensitive to measurement of known changes in the lens proteome with age.

The prevailing hypothesis for ARNC formation is that the accumulation of oxidized species results in the misfolding or aggregation of proteins that disrupts transparency and the gradient index of refraction in the lens. As presented in the introduction, oxidative stress response must occur in the inner nucleus, where fiber cells are metabolically less active and are not connected to a vascular transport system. Stress response is then thought to be mediated by the lens MCS delivery of reduced GSH and other metabolites to the inner nucleus through extracellular space along fiber cell sutures, and the export of metabolites through intercellular contacts (gap junctions) toward the equator of the lens. As such, this discussion of proteome measurements is primarily focused on the oxidative stress response and the physiology of the MCS before and after 50 years of age.

### Antioxidant protein networks

Previous lens proteome studies have identified putative transporters of GSH and its precursors, but not in the human lens (8, 10, 59). GSH uptake from extracellular suture space is putatively mediated by organic anion transporter 3 (OAT3) and sodium-dependent dicarboxylate transporter 3 (NaDC3) and efflux is mediated by multidrug resistance proteins 4 and 5 (MRP4/5) and connexins 46 and 50 (GJA3/8), which can transport molecules up to approximately 1 kDa through gap junction channels (15). The transport of GSH-precursor amino acids (glutamate, cysteine, and glycine) is also important for *in vivo* synthesis of GSH. These transport proteins include glycine transporters sodium- and chloride-dependent glycine transporters 1 and 2 (GLYT1/2), glutamine/glutamate transporter neutral amino acid transporter B(0) (ASCT2), glutamate and sodium cotransporters excitatory amino acid transporters 1 and 2 (EAAT1/2) and cysteine/glutamate antiporter complex (System X_c_^-^) composed of 4F2 cell-surface antigen heavy chain (4F2 or SLC3A2) and light-chain cysteine/glutamate transporter (XCT or SLC7A11). The functionality of GSH synthesis in the lens is beyond the scope of this study, but we note that the rate limiting enzyme of GSH synthesis – glutamate cysteine ligase catalytic subunit (GCLC) is measured in all 48 samples, but its abundance is not changed with age. As a cytoplasmic protein, it is expected that GCLC would not be measured in the membrane fraction (60), however it is possible that it is unfolded and aggregated as part of cellular maturation or tightly binds to the plasma membrane in its native folded state as observed for crystallins (32). Finally, while GSH is believed to be the antioxidant most responsible for lens homeostasis, ascorbic acid (vitamin C) is also proposed to be transported through the lens as an antioxidant in its oxidized form via glucose transporter 1 (GLUT1) and its reduced form by sodium-coupled vitamin C transporter 2 (SVCT2) as an alternative route for oxidative stress response.

For each of the proteins described above, the frequency of identification and description of sample distribution is given in Table 1 (graphical abundances in Supplemental Figure S3). From these data, several key observations were made. First, the most ubiquitous antioxidant transporter measured in the membrane is OAT3, facilitating uptake of intact GSH. Age does not appear to have an effect on the abundance of OAT3 in the membrane. Next, that NaDC3, MRP4, EAAT1, and SVCT2 are only measured in the youngest cortical samples, indicates little importance in long-lived proteome homeostasis. System X_c_^-^, a two-protein complex thought to contribute to cysteine accumulation in the lens nucleus for thiol protection is not measured as intact in these samples as XCT is not measured in any sample. Concurrently, 4F2 is less abundant in the membrane with increasing fiber cell age and subject age. Finally, GLUT1 is measured in all samples and its abundance does not change with age. Our results suggest that the influx and efflux of GSH and vitamin C through the lens is possible via substrate specific transport proteins, but that GSH precursor transport is challenged by subject and fiber cell age. Future experiments should be directed towards specific measurement of missing or otherwise low abundance antioxidant transporters to further support the spatial regulation and absence of these transporter proteins.

**Table 1.**
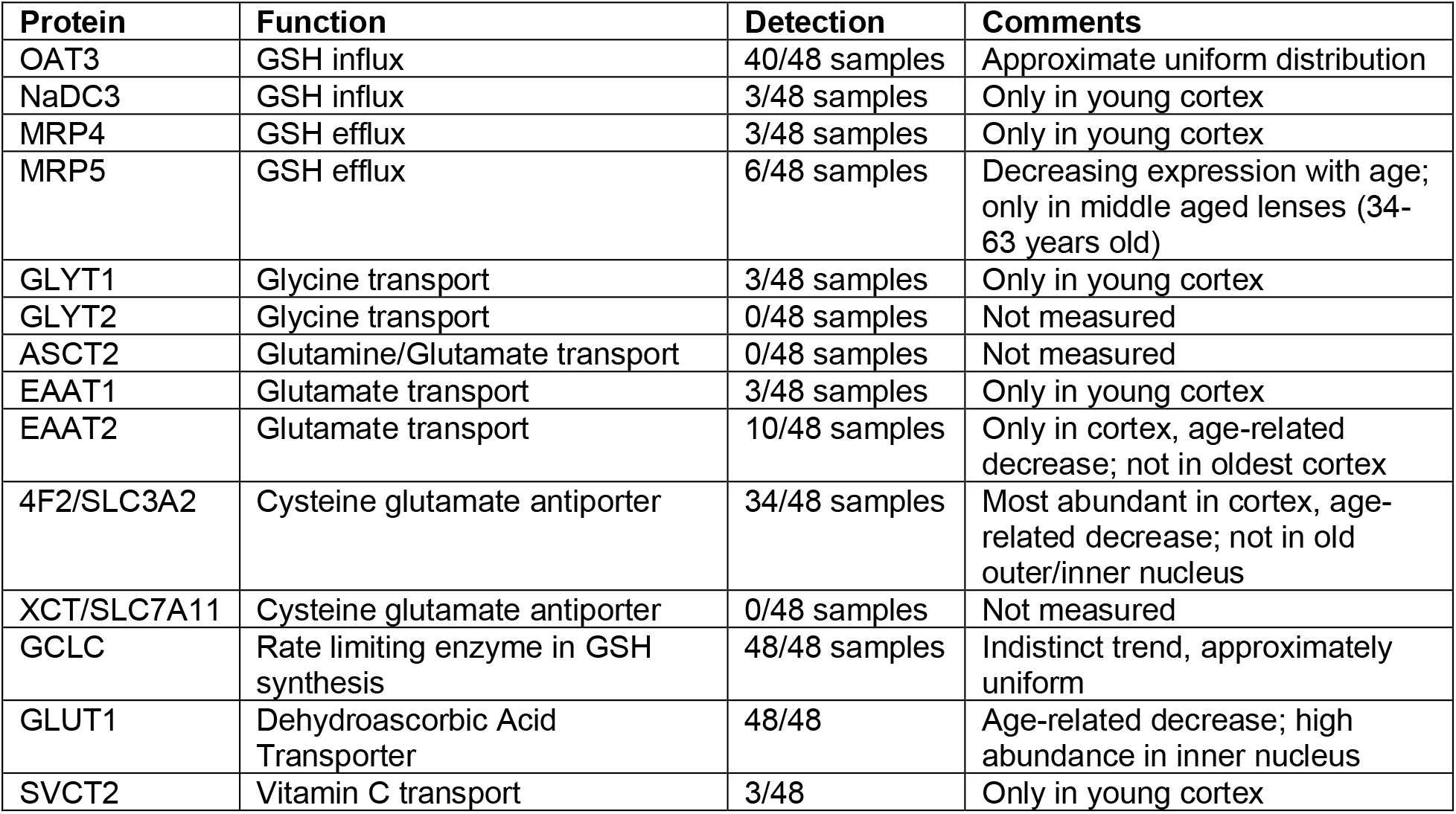
Proteins involved in antioxidant transport in the lens, their putative function, detection in the lens cohort and approximate age-related distributions. Visualized abundances represented in Supplemental Figure S6

Our DIA analysis allowed additional novel observations to be made regarding oxidative stress response in the lens. First, PCA separation of cortical fiber cells showed kynurenine oxoglutarate transaminase, responsible for formation of kynurenic acid from kynurenine and oxoglutarate, is enriched in older subject donors. Interestingly, kynurenic acid is shown to upregulate the activity of the GCLC transcription factor Nrf2, suggesting a role for the transaminase in oxidative stress response in transcriptionally regulated cortical fiber cells (61). We also identified several key cytosolic proteins involved in GSH-mediated oxidative stress response. These include glutathione-s-transferase (GST) and glutathione reductase (GR) which are ubiquitously measured in each region of the lens (Supplemental Figure S4). Each of these proteins show a slight increase in measured intensity at approximately 50 years, but no age-dependent statistical significance is detected. Conjugation of thiols to reduced cysteine residues, while preventing protein aggregation, may result in protein misfolding and loss of function. GST can restore a thiolated cysteine to its reduced form to allow restoration of perturbed structure and function (62). Extensive work by Lou and colleagues showed a thioltransferase system established by glutaredoxin proteins is 50% more efficient than equivalent GST function. However, the glutaredoxin family of proteins were not measured in the DIA dataset suggesting that they do not interact with the plasma membrane and that plasma membrane cysteine dethiolation is primarily achieved through non-glutaredoxin routes which may include GST. Functionality of GR is known to decrease with age, but a more significant decline in its activity is observed in progressively more severe age-related cataract (63). The consistent measure of GST and GR in all lenses studied suggests that oxidoreductase pathways may be at least partially functional in older lenses.

### Physiology of the lens microcirculatory system

#### Ion gradient establishment

Sodium-potassium ATPases along the epithelium are responsible for the establishment of a sodium cation gradient in the lens. It is not immediately apparent that these ATPases are modified in abundance in young or old lenses, suggesting that there is little change in expression with respect to age. This measurement may be artifactual as fiber cells were not separated from the epithelium, where these ATPases are functionally assigned in the MCS. Coupled to Na/K ATPases, a non-selective sodium leak channel is hypothesized to be responsible for the continuous influx of sodium to mature fiber cells. A suggested protein responsible for this, non-selective sodium leak channel, NALCN, was not measured in any sample. The alternative hypothesis to NALCN as a leak channel is GJA3, which is ubiquitously measured in the lens. Previous biophysical studies have identified GJA3 as critical in maintenance of sodium conductivity in the lens (64) but not as a leak channel explicitly. As described in results (Figure 6), GJA3 does not decrease in abundance as a result of the proteome remodeling event.

#### Cell-cell junctions

A consistently measured change in PSEA-quant analysis for each region of the lens was age-related depletion of cell-cell adhesion and junctions (e.g., GO:0005911, GO:0034329, GO:0045216). Cell-cell junctions of the lens are largely established by connexin gap junctions, actin-network tight junctions, and AQP0 (16, 29, 65). Connexin proteins play a key part in the transport of metabolites in the lens via formation of GJA3 and GJA8 gap junctions. For GJA3, abundance change is not observed (Figure 6), but there is gradual decrease in GJA8 abundance with fiber cell age and proteome remodeling (Supplemental Figure S5). A statistically significant (p < 0.001) change in the inner nucleus occurs with proteome remodeling at 50 years. These decreases cannot be explained solely by deamidation, as represented by age-related accumulation of GJA8 G265-K273 proportional deamidation (Supplemental Figure S5) and it instead suggested that a real decrease in protein abundance by truncation occurs. From prior DDA analyses, it is known that connexins are truncated at the N-terminus, C-terminus, or undergo cleavage in a central cytoplasmic loop (66). Gap junction channel formation is hypothesized to be disrupted by phosphorylation and pH; however, C-terminal truncation of each connexin leaves a modified channel that is functionally resistant to phosphorylation or pH variation present in the lens (67). Thus C-terminal truncation is believed to be an essential PTM in lens development. Neither gap junction is functionally restricted by truncation of the N- or C-terminus, however truncation by cleavage at a mid-sequence, cytoplasmic loop restricts hemi-channel formation (66). Evaluating the cytoplasmic loop peptide E110-K139, we show that this peptide decreases similarly to the whole GJA8 protein group in the cortex, but in inner nucleus samples older than 43 years, is not detected (Supplemental Figure S5D). In the outer nucleus, we measured this peptide in lenses up to 53-years-old. This demonstrates that GJA8 undergoes progressive decreases in abundance, but in mature fiber cells of older lenses, is not functional, decreasing net connexin gap junction permeability in older lenses.

Non-connexin junctional proteins decreased with the age-related remodeling in the cell-junction or cytoskeleton associated structural protein family of terms include vinculin, cell adhesion molecule 1/2/3, lens fiber membrane intrinsic protein 2, and cadherin-2. Many of the proteins associated with cell-cell and cell-junction ontologies are calcium-dependent for maintenance of adhesion. The functional relevance of the decreased abundance of these proteins in older lenses is unclear, but similar cell junction ontology terms to those measured here have been detected in previous analysis of progressively older fiber cell populations (37, 68). Several of the cell-junction proteins decreased with proteome remodeling have known interactions with calcium, leading us to investigate other calcium-interacting proteins. Calcium acts as a second messenger, and in addition to calmodulin-dependent inhibition of water permeability through aquaporin-0, regulates actin cytoskeleton integrity and the activity of the ubiquitin proteasome system (UPS), and calpain proteases (69–71). Prior measurements of the lens demonstrate that there is a parabolic decrease of calcium concentration in the lens with age, reaching its lowest point around 50 years, and increasing in lenses thereafter, with significantly elevated levels of calcium in cataract lenses (72). The accumulation of gene ontology terms presented here led us to evaluate the effect of aging on selective calcium transporters.

#### Calcium transport

To evaluate age-related changes in calcium transporters in the lens, we first isolated the subset of broadly defined solute-carrier membrane (SLC) proteins. This was done to evaluate the significant changes in all SLC transporters prior to assigning significance to an arbitrary protein group. Of 121 SLC protein groups, only 21 candidates were measured in at least 36/48 samples. This allowed filtering for SLC proteins that may be completely absent in old inner nucleus, but required that they be measured at young age, when calcium export is putatively functional. From this subset list of 21 SLC proteins, we evaluated if age-related changes in abundance, consistent with the proteome remodeling event at 50 years, occurred. Many SLC proteins undergo statistically significant reduction in representation with age in the cortex, but this is anticipated to be a result of decreased organelle concentration and loss of protein synthesis in old fiber cells relative to young fiber cells. Thus, we elected to only identify candidate SLC proteins where statistical reduction occurred in at least two lens regions, reducing the 21 SLC candidate proteins to 2 proteins of interest. The first is SLC30A1, a relatively low abundance zinc transporter not measured in the old inner nucleus and significantly decreased in the outer nucleus. Interestingly, the only other protein which showed consistent, age-related decrease in abundance was SLC24A2, a light-activated sodium, potassium, calcium exchange protein. The abundance profile of SLC24A2 with age (Supplemental Figure S6) demonstrates little change in the cortex, but especially interior to the established diffusion barrier, a remodeling-dependent decrease in abundance occurs. No other members of the sodium, potassium, calcium exchange protein family were detected.

It is unclear whether SLC proteins contribute to intracellular transport after the extracellular space between fiber cells is decreased with fiber cell maturation. However, considering biophysical evidence, we suggest that the age-related proteome remodeling event at 50 years of age is responsible for functional restriction of calcium export from lens fiber cells. Thus, if SLC24A2 contributes to calcium export, it is expected that its degradation is a precursor to calcium accumulation and subsequent signaling events. The functional alternative to SLC24A2 for calcium transport is through non-selective connexin gap junction channels. Recent studies show that GJA8 is permeable to calcium (73). In contrast, calcium binding to GJA3 decreases channel permeability and age-related accumulation of calcium may lead to inhibition of GJA3 function. In the context of GJA3 and GJA8 age-related abundances (Figure 6, Supplemental Figure S5), we suggest that the age-related decrease in GJA8 abundance and function may inhibit net calcium export from the lens resulting in calcium accumulation, which in turn inhibits the permeability of GJA3, for which abundance is not significantly decreased with age. Taken together, it is anticipated that proteome remodeling at 50 years directly contributes to calcium accumulation in the lens and that accumulation may lead to inhibition to MCS as a whole. The physiological significance of this being that a current of metabolic waste is less efficient than in young lenses, identifying a potential role of calcium in cataract formation (Figure 7).

**Figure 7.**
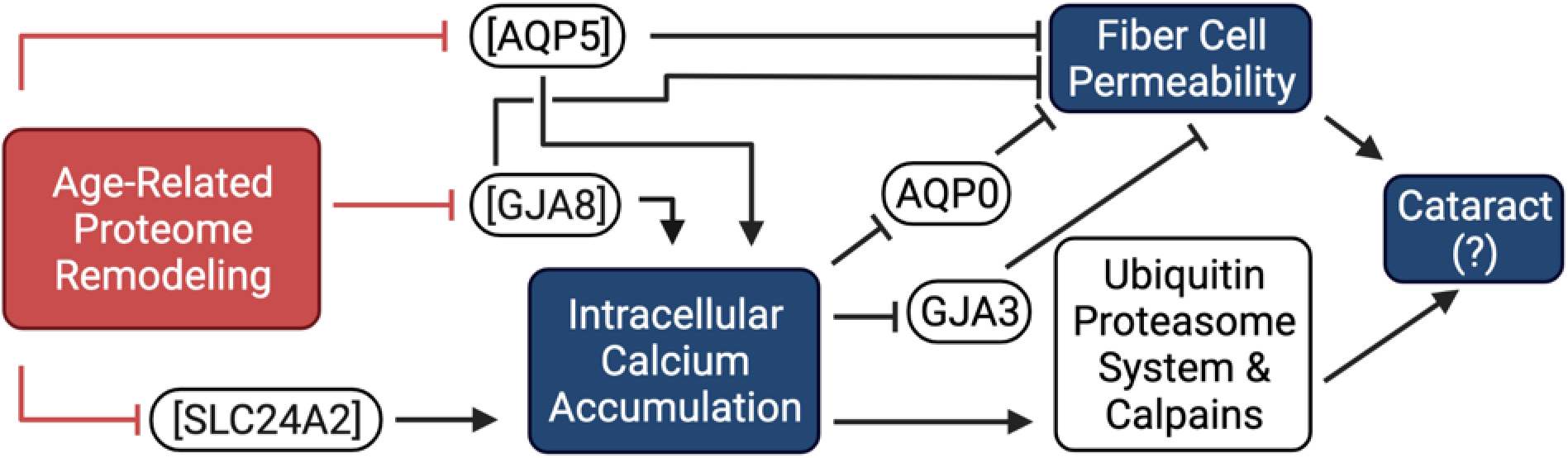
Schematic of suggested effect of age-related proteome remodeling. Remodeling results in decreased abundance of AQP5, decreasing fiber cell permeability and decreases GJA3 and SLC24A2 abundances, resulting in calcium accumulation. Calcium accumulation inhibits GJA8 and AQP0 functionality, further inhibiting fiber cell permeability. Fiber cell permeability inhibition may then result in cataract. Calcium accumulation also results in ubiquitin proteasome and calpain protease activation which may lead to cataract. Activation, →, and inhibition, ⊥, is indicated in reference to proteome remodeling treatment and not necessarily young lens function.

Aside from MCS functionality, prior studies establish the UPS as functional in mature fiber cells, which lack significant metabolic activity (74). Further studies are necessary to establish calcium-concentration perturbation of UPS in mature lens fiber cells in the inner nucleus, however protein components of the proteasome are not age-degraded (Supplemental Figure S7). Our data suggest that calcium accumulation then results in UPS activation as in other cell types, initiating elevated rates of protein degradation and E3 ligase conferred substrate specificity (69). As a result, UPS activity may contribute to degradation of oxidative stress response proteins leading to further accumulation of reactive oxygen species and oxidized substrate proteins responsible for ARNC. Finally, it is anticipated that calcium accumulation results in activation of calpain proteases, including calpain-2 catalytic subunit (P17655) which is measured in all 48 samples with no age-related change in abundance (71). In K6W ubiquitin model systems, UPS is inactive and in the presence of accumulated calcium, calpain-associated fragmentation occurs on critical proteins including filensin, vimentin and β-crystallin. These lenses also formed cataract, further supporting a role of calcium accumulation in ARNC formation.

Future functional studies are necessary to validate 1) the function of SLC24A2 in calcium transport in mature fiber cells, 2) the direction of calcium current – here assumed to be moving from the inner nucleus towards the epithelium, 3) the role of calcium accumulation on ubiquitin proteasome function in the lens, and 4) the role of connexin gap junctions in calcium transport. If SCL24A2 is not responsible for calcium export, additional studies are required to determine why calcium accumulates in the lens in the presence of putatively function gap junction proteins.

#### Aquaporins

Previous publications have thoroughly reviewed the importance of aquaporins in the lens MCS (12, 27, 75–78). Briefly, aquaporins serve as water channels that facilitate water permeability in fiber cells. Aquaporin-1 is exclusively expressed in epithelial cells and is detected at only trace levels in fiber cells still differentiating from epithelial cells. In addition to AQP1, two additional aquaporin proteins, AQP0 and AQP5 play roles in lens MCS as well as in cell-cell adhesion (65). AQP0 is the most abundant membrane protein in lens fiber cells, and it undergoes truncation of the C-terminus as an age-related modification throughout the lens (77). Truncation of AQP0 has been visually mapped with imaging mass spectrometry (27). These reported findings are supported by Supplemental Figure S8, demonstrating age-related reduction of the C-terminal tryptic peptide AQP0 G239-L263 in the cortex and absence from aged outer nucleus and inner nucleus samples. The abundance profile of the AQP0 N-terminal A12-R33 peptide and intact protein group are consistent with each other, but not with the abundance of this C-terminal G239-L263 peptide. This truncation site is then in agreement with top-down and DDA proteomics studies (27, 77). Since AQP0 is degraded continuously and starts early in maturation, it is not believed that proteome remodeling is responsible or caused by the truncation of AQP0.

Aquaporin-5 is a second, less abundant aquaporin family member measured in lens fiber cells. Though structurally similar to AQP0, AQP5 has approximately 20-fold higher water permeability than AQP0 (79). While AQP0 is consistently localized in the plasma membrane, AQP5 originates in lysosomal structures and is translocated to the plasma membrane as a part of maturation and in response to of osmotic stress (12, 19). Unlike AQP0, which does not undergo any significant change in abundance with the proteome remodeling event, AQP5 undergoes a significant decrease in abundance with age in the inner nucleus (Supplemental Figure S9). To evaluate if a significant change to AQP5 abundance is derived from deamidation, we evaluated the accumulation of deamidation on AQP5 S189-R198. The change in proportional deamidation on this peptide and the other deamidated peptide (G241-R262, not shown) is incongruent with the magnitude of protein abundance decline, which suggests that deamidation does not account for the decrease in measured AQP5 abundance. Thus, we speculate that AQP5 is further modified in an age-dependent manner, which may include truncation. The functional implication of AQP5 modification may be reduced export of water from the inner nucleus to the epithelium, resulting in decreased cellular permeability. Because water permeability must first be established in the inner nucleus to transport small molecules towards the epithelium, the decrease in unmodified AQP5 abundance has putative functional consequence on the net current of water and small molecules throughout the lens. The net effect of AQP5-relative to AQP0-mediated water permeability in the inner nucleus has not been established, but AQP5-degradation may result in accumulation of small molecules including calcium and oxidized GSH. Ultimately, this may result in further proteostatic and homeostatic stress, leading to eventual ARNC formation.

## Summary

In summary, we demonstrate that DIA identifies more proteins in the lens than previously possible and delineates age-related changes in defined spatiotemporally distinct regions. Novel to this study, we show that a proteome remodeling event occurs at 50 years of aging and that oxidative stress response networks are retained with age, while cell-cell contacts are degraded with age. Expanding on this, we demonstrate the first proteomic identification of multiple GSH and ascorbic acid transport proteins in the human lens. In addition to supporting prior measurements of the aging lens, we identified that calcium transporter SLC24A2 is less abundant after proteome remodeling, leading to its accumulation and potential inhibition of the MCS, which is essential for maintaining tissue transparency. Finally, we show that AQP5 is depleted in the inner nucleus of the lens, coincident with proteome remodeling; a finding with important physiological implications for MCS activity changes with age (Figure 7). Further functional studies are needed to evaluate these hypotheses, especially related to calcium accumulation in the aging lens. Additionally, it is desirable to further expand this approach to cataract lenses to clearly delineate changes that occur in ARNC lenses relative to old, healthy lenses.

## Data Availability

Raw MS data, deconvoluted mzML files, DIA-NN created .dia files, DIA-NN reports, and R script saved data are available on ProteomeXchange. These data can be accessed using the project accession number PXD033722.

## Supplemental Data

Supplemental Figure S1: Distribution of protein abundances after protein group normalization.

Supplemental Figure S2: Abundance of GJA3 when deamidation is included in protein measurement

Supplemental Figure S3: Abundance of each measured protein in antioxidant transport

Supplemental Figure S4: Abundance of glutathione-s-transferase and glutathione reductase

Supplemental Figure S5: Abundance of GJA8, G265-K273 deamidation peptide, and E110-K139 cytoplasmic loop peptide

Supplemental Figure S6: Abundance of SLC24A2

Supplemental Figure S7: Abundance of several components of the proteasome

Supplemental Figure S8: Abundance of AQP0 and C-terminal peptide G239-L263

Supplemental Figure S9: Abundance of AQP5 and S189-R198 peptide

Supplemental Table S1: List of proteins significantly changed in volcano plot analysis

Supplemental Table S2: List of GO terms enriched in the young cortex over old cortex

Supplemental Table S3: List of GO terms enriched in the old cortex over young cortex

Supplemental Table S4: List of GO terms enriched in the young outer nucleus over old outer nucleus

Supplemental Table S5: List of GO terms enriched in the old outer nucleus over young outer nucleus

Supplemental Table S6: List of GO terms enriched in the young inner nucleus over old inner nucleus

Supplemental Table S7: List of GO terms enriched in the old inner nucleus over young inner nucleus

## Acknowledgements

The authors wish to acknowledge the Vanderbilt Mass Spectrometry Research Center, particularly Kristie L. Rose and Purvi Patel for assistance in acquiring DIA data for this experiment. We also thank Zhen Wang, Romell B. Gletten, Sarah R. Zelle and all other members of the Schey lab for thoughtful conversations in the development of this work. Financial support is acknowledged from NIH grants EY013462 (KLS), EY008126, T32 GM065086 (LSC), and NSF GRFP (LSC).

