## Supplemental Figures S1-S9 for "Proteome Remodeling of the Eye Lens at 50 Years Identified with Data-Independent Acquisition"

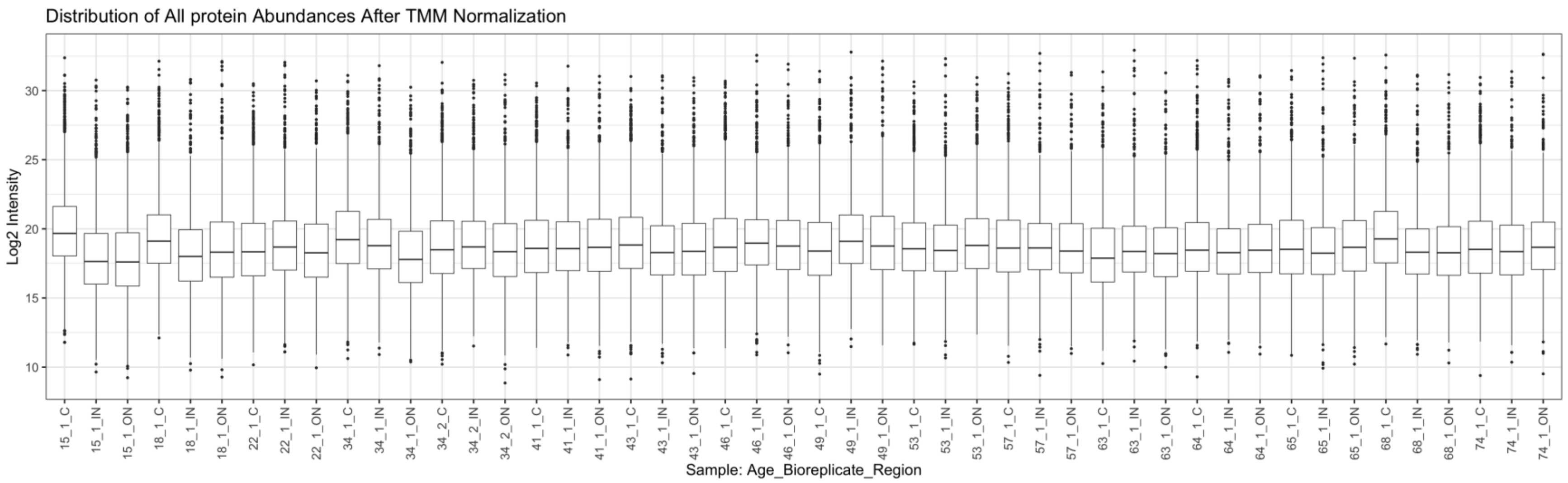


*Supplemental Figure S1 - Distribution of protein groups after TMM normalization was performed on all protein groups measured in every sample. Deviation from population distributions is indicative of representative abundance change for protein groups not measured in all samples*

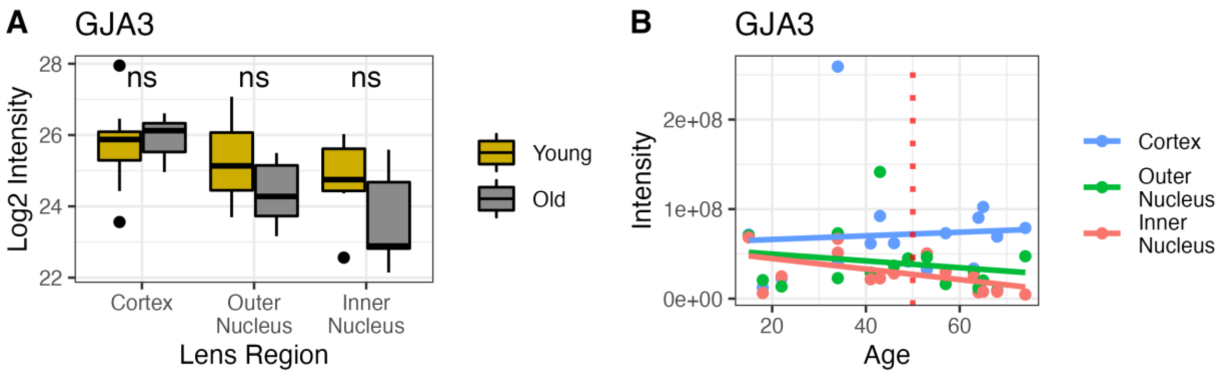


*Supplemental Figure S2 - Abundance of Connexin 46 (GJA3) when deamidated residues are considered towards the total abundance of the protein group. When GJA3 is unmodified (Figure 6), a significant change is measured by two-sample t-test. A) When deamidation is included, no significance is established; B) There is a qualitative decline in GJA3 abundance in the outer and inner nucleus with age, but it is not consistent with the proteome remodeling event. T-test significance cutoffs were set at * = <0.05, ** = <0.01, *** = <0.001, **** =<0.0001*

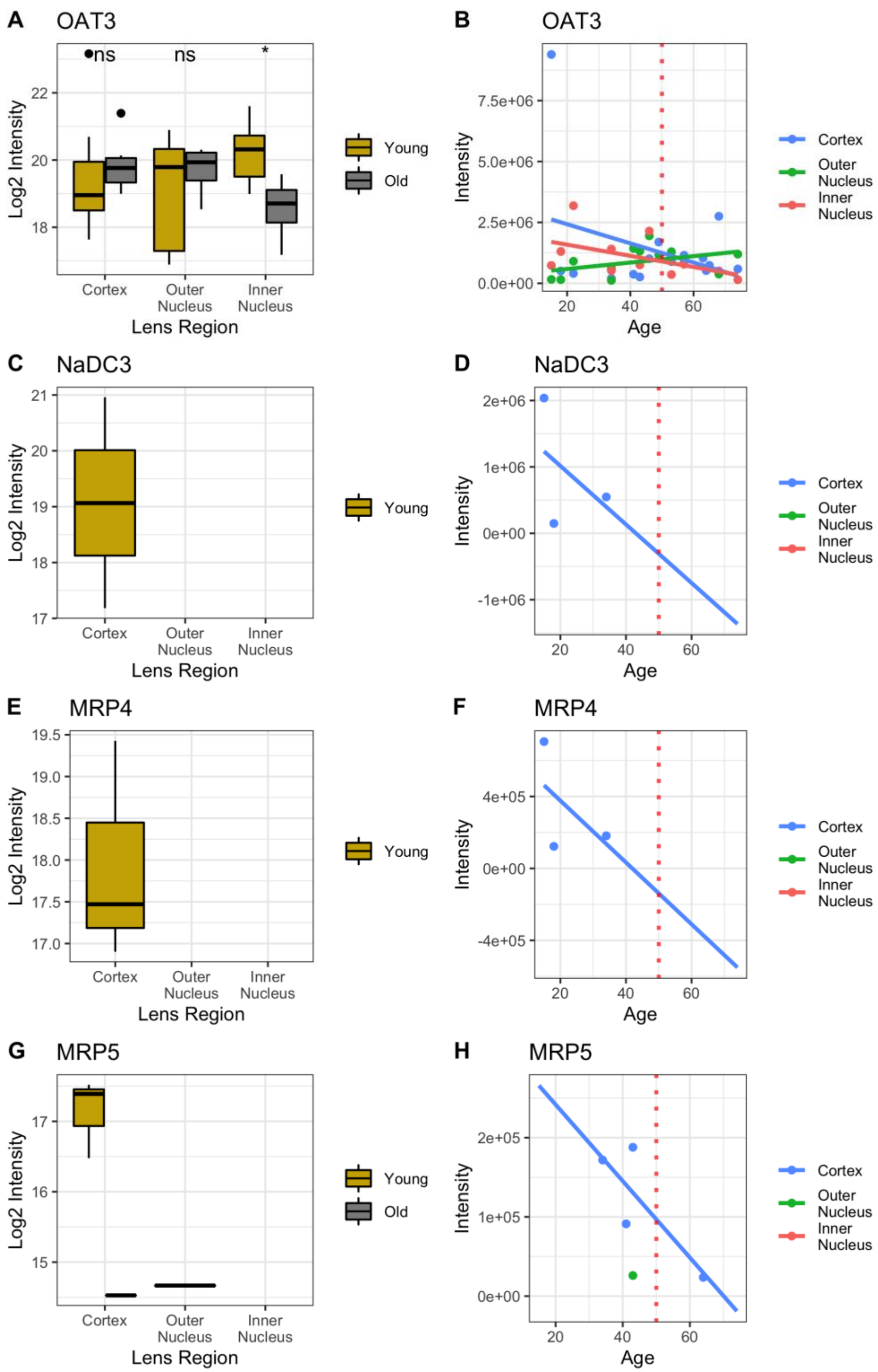
A
OAT3
ns
ns
*
Log2 Intensity
Lens Region
Cortex
Outer Nucleus
Inner Nucleus
Young
Old
B
OAT3
Intensity
Age
Cortex
Outer Nucleus
Inner Nucleus
C
NaDC3
D
NaDC3
E
MRP4
F
MRP4
G
MRP5
H
MRP5

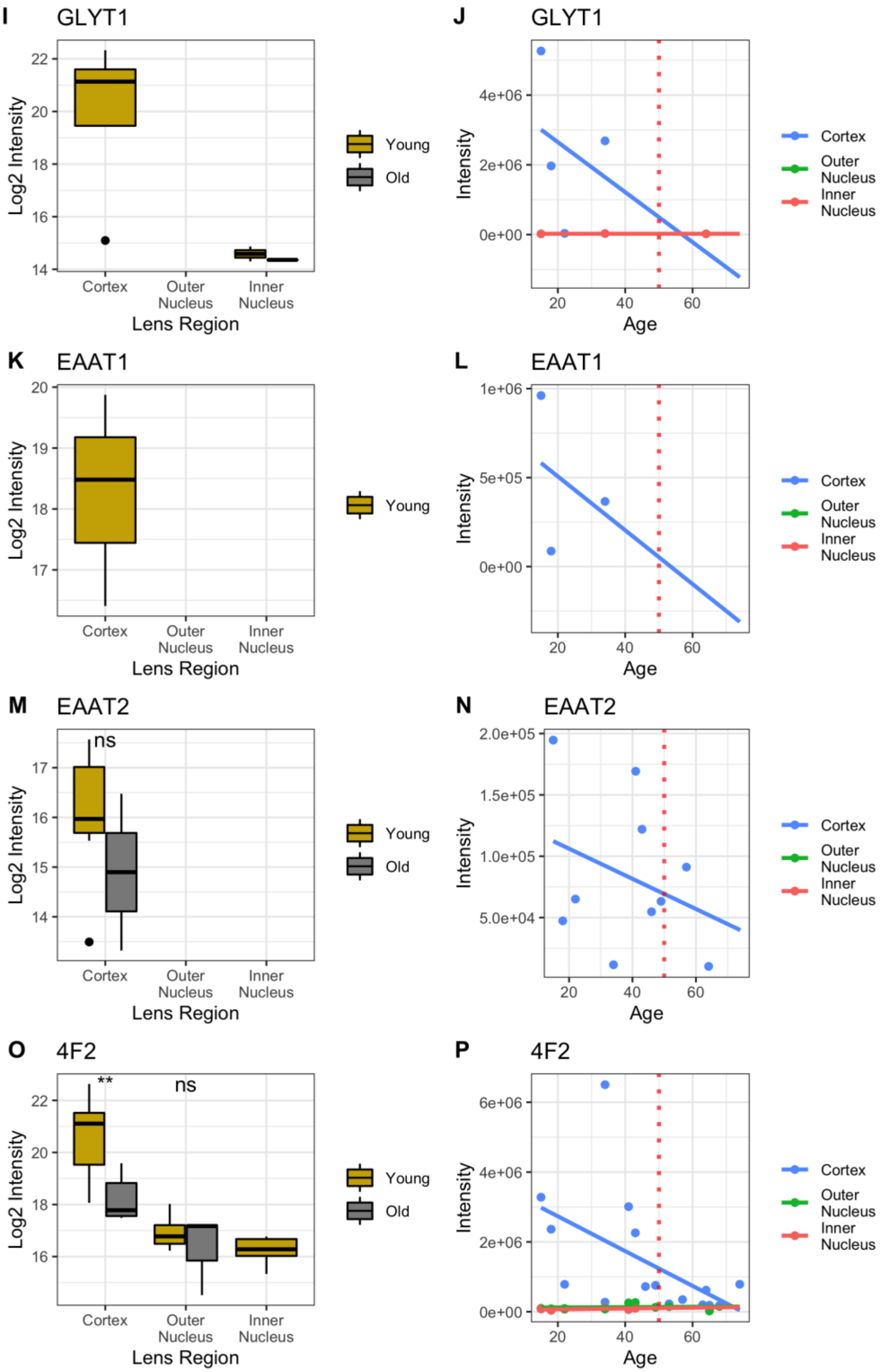
I
GLYT1
Log2 Intensity
22
20
18
16
14
Cortex
Outer Nucleus
Inner Nucleus
Lens Region
Young
Old
J
GLYT1
Intensity
4e+06
2e+06
0e+00
20
40
60
Age
Cortex
Outer Nucleus
Inner Nucleus
K
EAAT1
Log2 Intensity
20
19
18
17
Cortex
Outer Nucleus
Inner Nucleus
Lens Region
Young
L
EAAT1
Intensity
1e+06
5e+05
0e+00
20
40
60
Age
Cortex
Outer Nucleus
Inner Nucleus
M
EAAT2
ns
Log2 Intensity
17
16
15
14
Cortex
Outer Nucleus
Inner Nucleus
Lens Region
Young
Old
N
EAAT2
Intensity
2.0e+05
1.5e+05
1.0e+05
5.0e+04
20
40
60
Age
Cortex
Outer Nucleus
Inner Nucleus
O
4F2
**
ns
Log2 Intensity
22
20
18
16
Cortex
Outer Nucleus
Inner Nucleus
Lens Region
Young
Old
P
4F2
Intensity
6e+06
4e+06
2e+06
0e+00
20
40
60
Age
Cortex
Outer Nucleus
Inner Nucleus

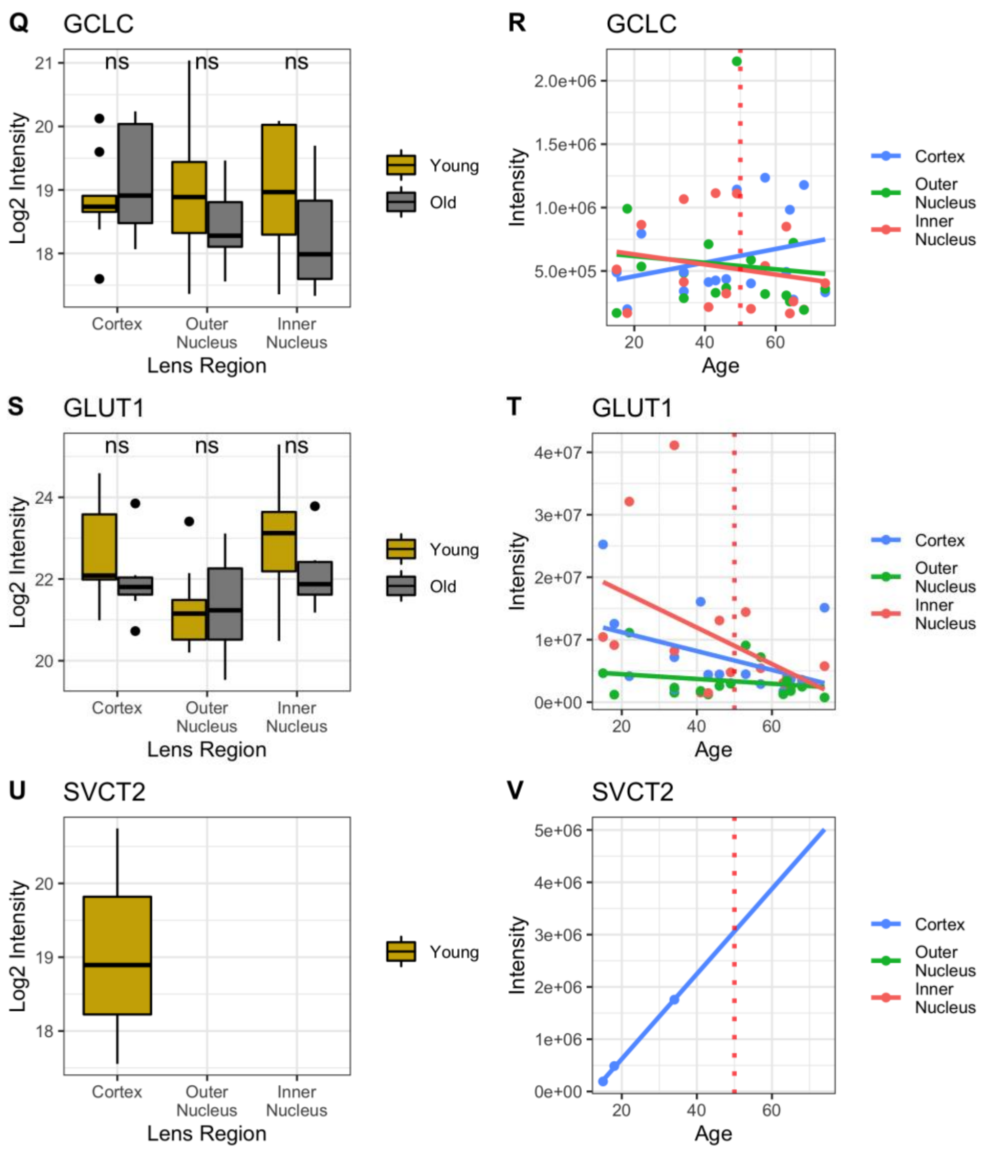


*Supplemental Figure S3 - Age-related expression of each protein mentioned in the discussion as related to glutathione transport, glutathione synthesis or vitamin C transport. Proteins not measured are not plotted. T-test significance cutoffs were set at * = <0.05, ** = <0.01, *** = <0.001, **** =<0.0001*

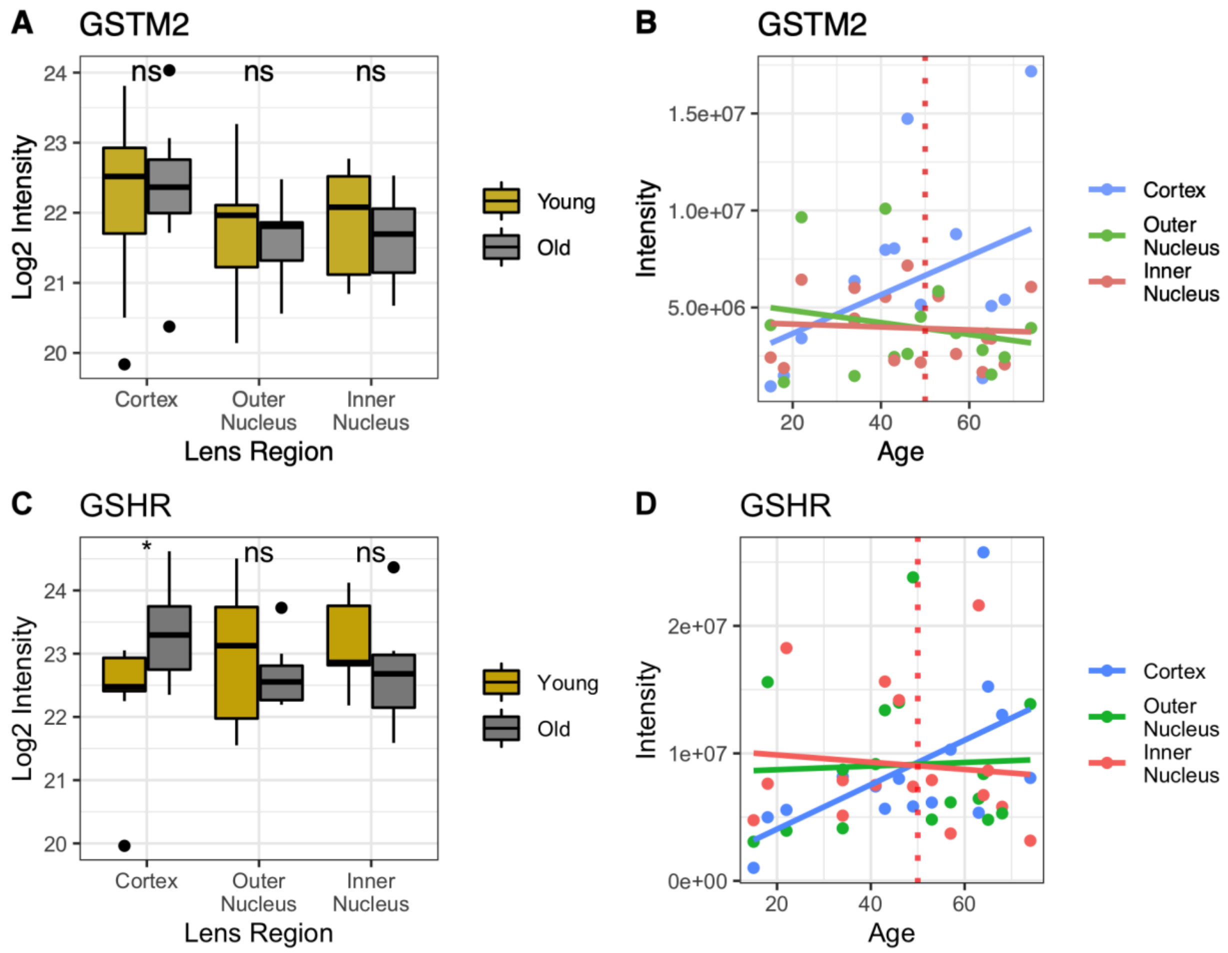


*Supplemental Figure S4 – A,C) Age-grouped t-test (* = <0.05, ** = <0.01, *** = <0.001, **** =<0.0001) and B,D) age-related expression of A,B) glutathione-s-transferase mu 2 (GSTM2) and C,D) GSH reductase (GSHR). There is not a clear statistical change in the representation of either protein within the dataset and each protein is consistently measured throughout the lens. It is hypothesized that the accumulation of each cytosolic protein represents insolubilization by misfolding.*

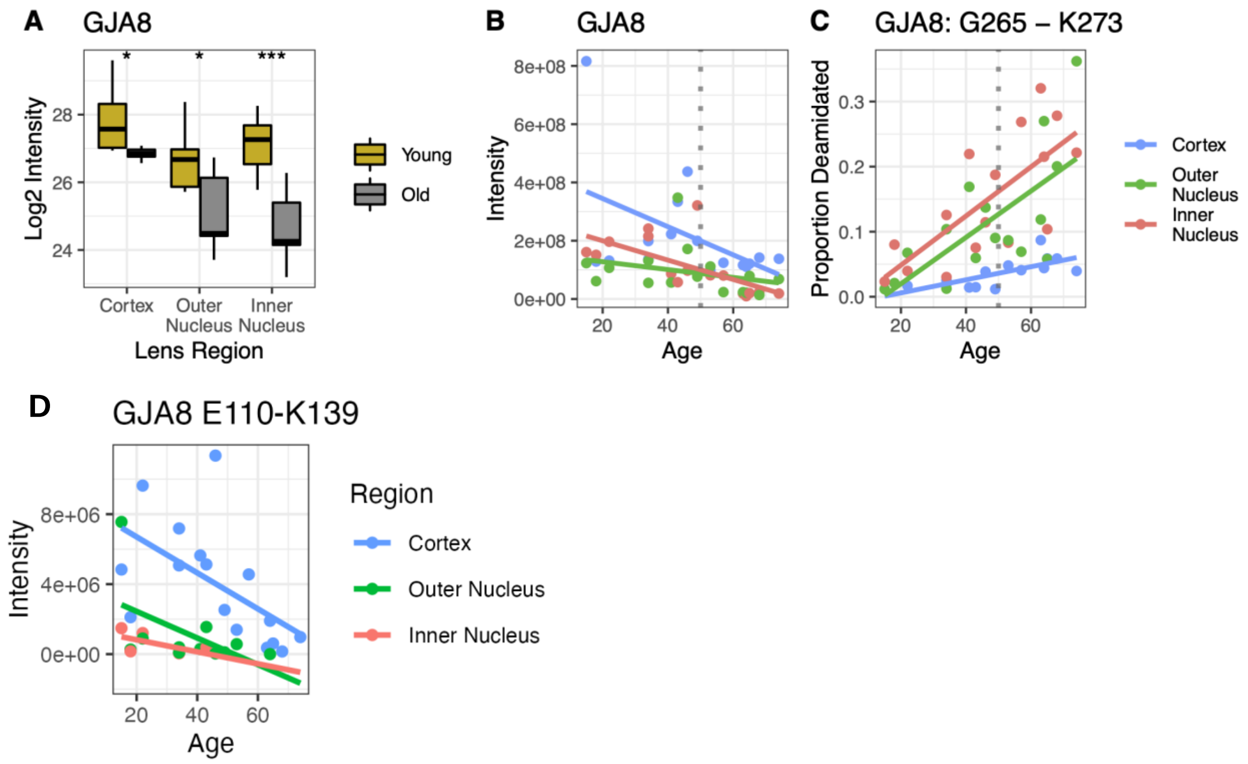


*Supplemental Figure S5 - Demonstration of age-related abundance change of connexin 50 (GJA8) and A) An age-related decrease was appreciable for GJA8, especially in the inner nucleus, and B) decrease in protein expression cannot be solely attributed to PTMs, as shown by C) accumulation of deamidation on the G265-K273 peptide. D) Abundance of a cytoplasmic loop region peptide (E110-K139) for GJA8 demonstrates that cytoplasmic loop cleavage and functional deletion occurs in older nuclear lens regions after 50 years. T-test significances shown as * = <0.05, ** = <0.01, *** = <0.001, **** =<0.0001*

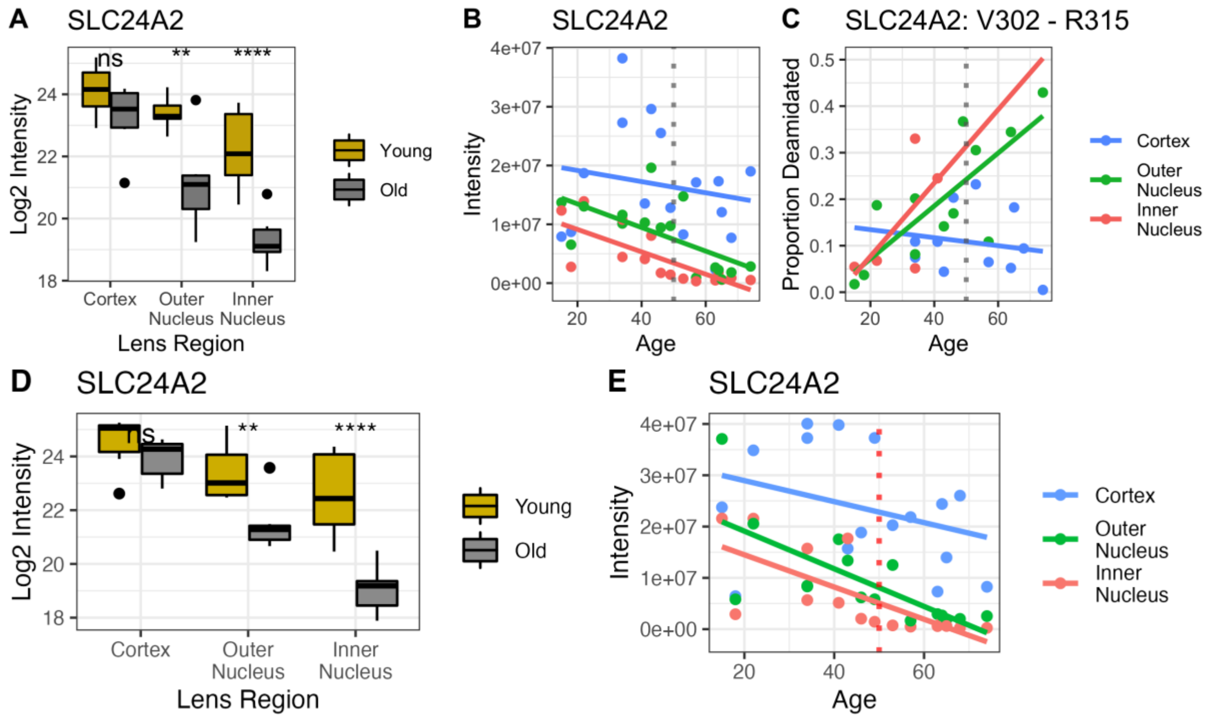


*Supplemental Figure S6 - Demonstration of age-related abundance change of SLC24A2 (Sodium, Potassium, Calcium Exchange Protein 2). A) A proteome-remodeling related decrease in SLC24A2 abundance was appreciable in each nuclear region, especially the inner nucleus, and B) There is a steady linear decline of SCL24A2, however, there is a significant decrease in the abundance of measured SLC24A2 in the inner nucleus young region relative to old region, which is not consistent with linearity. C) Proportional deamidation of V302-R315 on SLC24A2 is demonstrated, showing some accumulation of deamidation with age, but that nuclear samples are less likely to be singly deamidated, suggesting further modification or decrease in abundance below the limit of detection and that PTMs alone are unlikely to explain the decrease in measured abundance. D, E) When t-test significance and linearity of protein abundance change was assessed with deamidation enabled, identical trends to those measured in the unmodified dataset were demonstrated. T-test significances shown as * = <0.05, ** = <0.01, *** = <0.001, **** =<0.0001*

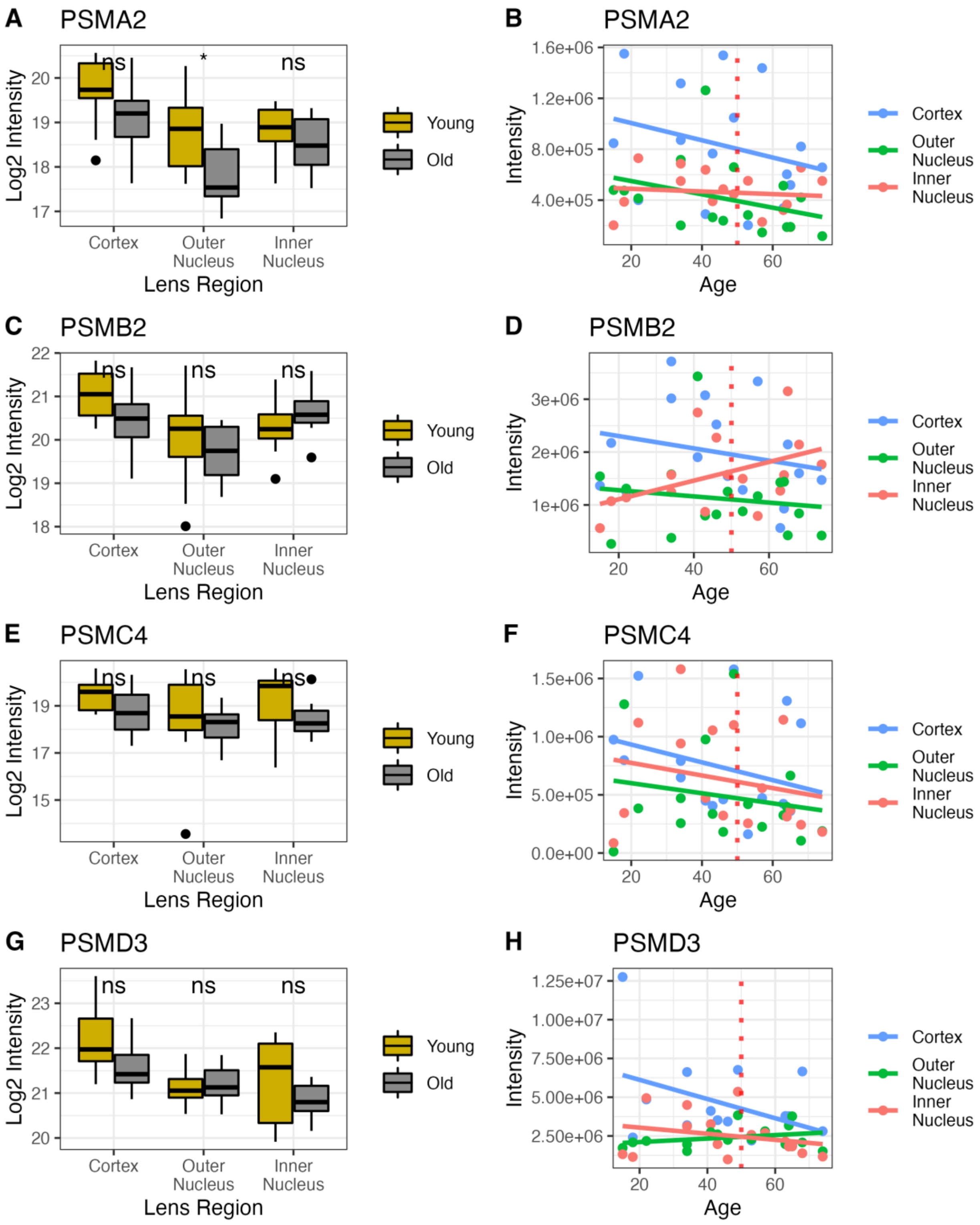
A
PSMA2
Log2 Intensity
ns
*
ns
17
18
19
20
Cortex
Outer Nucleus
Inner Nucleus
Lens Region
Young
Old
B
PSMA2
Intensity
4.0e+05
8.0e+05
1.2e+06
1.6e+06
20
40
60
Age
Cortex
Outer Nucleus
Inner Nucleus
C
PSMB2
Log2 Intensity
ns
ns
ns
18
19
20
21
22
Cortex
Outer Nucleus
Inner Nucleus
Lens Region
Young
Old
D
PSMB2
Intensity
1e+06
2e+06
3e+06
20
40
60
Age
Cortex
Outer Nucleus
Inner Nucleus
E
PSMC4
Log2 Intensity
ns
ns
ns
15
17
19
Cortex
Outer Nucleus
Inner Nucleus
Lens Region
Young
Old
F
PSMC4
Intensity
0.0e+00
5.0e+05
1.0e+06
1.5e+06
20
40
60
Age
Cortex
Outer Nucleus
Inner Nucleus
G
PSMD3
Log2 Intensity
ns
ns
ns
20
21
22
23
Cortex
Outer Nucleus
Inner Nucleus
Lens Region
Young
Old
H
PSMD3
Intensity
2.50e+06
5.00e+06
7.50e+06
1.00e+07
1.25e+07
20
40
60
Age
Cortex
Outer Nucleus
Inner Nucleus

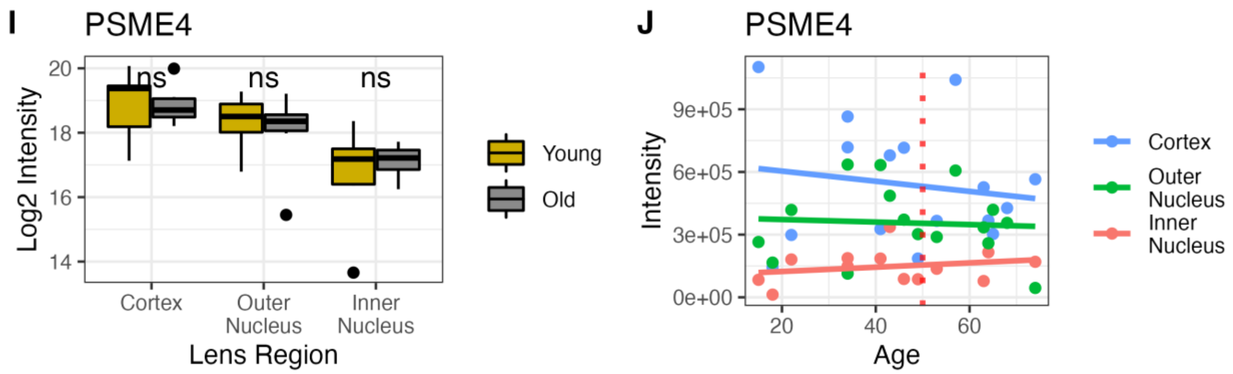


*Supplemental Figure S7 – Evaluation of several proteasome components and their statistical change relative to proteome remodeling event. All samples demonstrate some fiber cell maturation stage related degradation, but there is no significant change in the abundance of each proteasome component with age. T-test significance cutoffs were set at * = <0.05, ** = <0.01, *** = <0.001, **** =<0.0001 for boxplot comparison.*

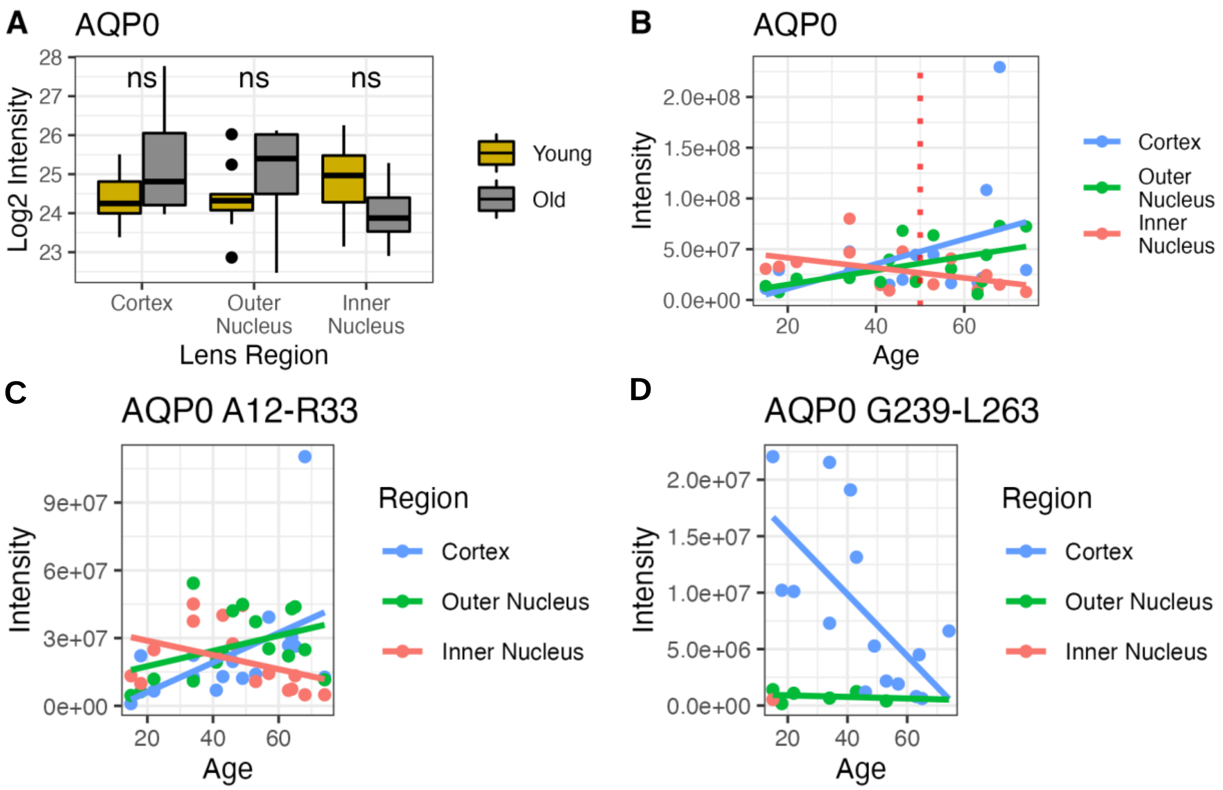


*Supplemental Figure S8 - Age-related abundance of AQP0 and two peptides. A) T-test of AQP0 abundance at either side of the proteome remodeling event. T-test significance cutoffs were set at * = <0.05, ** = <0.01, *** = <0.001, **** =<0.0001. B) Linear decline of AQP0 abundance is not established in any sample and closely resembles the abundance of C) AQP0 A12-R33, a ubiquitously measured peptide at the N-terminus of AQP0. D) The abundance of AQP0 G239-263 represents the abundance of a C-terminal peptide that is believed to be truncated in fiber cells with age. Coupled to evidence from B and C, it is suggested that measurement of G239-L263 supports identification of trucated residues with DIA analysis. The C-terminal peptide is detected in trace amounts in the aging outer nucleus and 15 y.o. inner nucleus, supporting the assignment of age-related truncation of AQP0.*

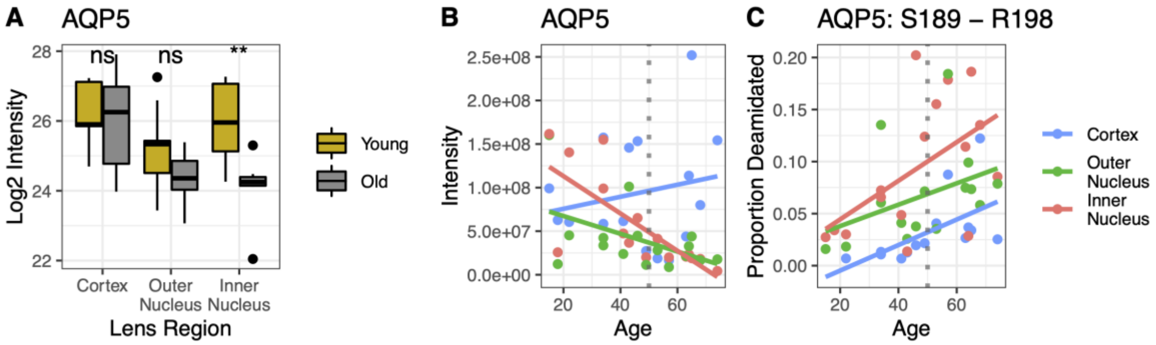


*Supplemental Figure S9 – A) T-test comparisons (* = <0.05, ** = <0.01, *** = <0.001, **** =<0.0001) and distribution of AQP5 abundance relative to the proteome remodeling event and B) AQP5 distribution with age relation. Increase of cortical AQP5 is not necessarily indicative of increased cortical expression, but instead that AQP5 contributes proportionally to the lens proteome more significantly in several biological replicates. Distribution and t-testing demonstrates that this is statistically insignificant . C) The proportion of deamidation on C-terminal peptide G241-R253 demonstrates that the accumulation of deamidation alone is not responsible for the age-related decrease in AQP5 representation in the lens proteome.*
