## Supplemental Table S1 for "Proteome Remodeling of the Eye Lens at 50 Years Identified with Data-Independent Acquisition"

Supplemental Table S1 – Protein groups filtered as differentially expressed in at least one lens region between young lenses and old lenses. Young lenses were defined as from human subject donors younger than 50 years, and older lenses from human subject donors older than 50 years. Imputed values were not used in this dataset. Negative Log2 Fold Change (Log2FC) indicates the protein is differentially present in the young lenses, and positive Log2FC indicates preferentially preserved in old lenses. P-values were calculated with a parametric t-test on log2 treated protein abundances. Protein groups determined as significant in Figures 2-5 are highlighted with their corresponding sample group (e.g., cortex significant proteins are highlighted red). Significance cutoffs were set at +/- 1.5 Log2FC and P-Values < 0.01.

| Protein Group | Gene Group | Protein Description | Cortex p-value | Cortex Log2FC | Outer Nucleus p-value | Outer Nucleus Log2FC | Inner Nucleus p-value | Inner Nucleus Log2FC |
|---|---|---|---|---|---|---|---|---|
| P04350;P07437;P68371;Q13885;Q9BVA1 | TUBB;TUBB2A;TUBB2B;TUBB4A;TUBB4B | Tubulin beta-4A chain | 1.12E-03 | -4.30 | 5.43E-01 | 0.24 | NA | NA |
| P07099 | EPHX1 | Epoxide hydrolase 1 | 2.19E-03 | -3.83 | NA | NA | NA | NA |
| Q16778;Q8N257 | H2BC21;H2BU1 | Histone H2B type 2-E | 1.62E-03 | -3.72 | NA | NA | NA | NA |
| Q6UW68 | TMEM205 | Transmembrane protein 205 | 1.34E-03 | -3.70 | NA | NA | NA | NA |
| Q9UEW8 | STK39 | STE20/SPS1-related proline-alanine-rich protein kinase | 1.16E-04 | -3.66 | NA | NA | NA | NA |
| P36578 | RPL4 | 60S ribosomal protein L4 | 2.94E-04 | -3.60 | NA | NA | NA | NA |
| P04350;P68371 | TUBB4A;TUBB4B | Tubulin beta-4A chain | 5.68E-04 | -3.59 | 7.69E-01 | -0.26 | NA | NA |
| Q9Y3D6 | FIS1 | Mitochondrial fission 1 protein | 2.72E-04 | -3.58 | NA | NA | NA | NA |
| P45880 | VDAC2 | Voltage-dependent anion-selective channel protein 2 | 9.11E-04 | -3.54 | 4.02E-02 | -1.87 | 1.19E-02 | -1.92 |
| P07437;P68371;Q13885;Q9BVA1 | TUBB;TUBB2A;TUBB2B;TUBB4B | Tubulin beta chain | 1.01E-04 | -3.54 | NA | NA | NA | NA |
| P35613 | BSG | Basigin | 1.07E-03 | -3.52 | NA | NA | NA | NA |
| P20339 | RAB5A | Ras-related protein Rab-5A | 2.28E-04 | -3.50 | NA | NA | NA | NA |
| P00403 | MT-CO2 | Cytochrome c oxidase subunit 2 | 8.22E-04 | -3.49 | 2.19E-02 | -1.59 | NA | NA |
| P14625 | HSP90B1 | Endoplasmin | 8.00E-04 | -3.46 | NA | NA | NA | NA |
| P06576 | ATP5F1B | ATP synthase subunit beta, mitochondrial | 1.12E-03 | -3.43 | NA | NA | NA | NA |
| Q9Y3B3 | TMED7 | Transmembrane emp24 domain-containing protein 7 | 8.64E-04 | -3.43 | NA | NA | NA | NA |
| P27824 | CANX | Calnexin | 1.15E-05 | -3.42 | NA | NA | NA | NA |
| P40429 | RPL13A | 60S ribosomal protein L13a | 9.18E-05 | -3.41 | NA | NA | 6.65E-02 | -1.27 |
| P35579;P35749;Q7Z406 | MYH11;MYH14;MYH9 | Myosin-9 | 3.48E-04 | -3.39 | NA | NA | NA | NA |
| P31930 | UQCRC1 | Cytochrome b-c1 complex subunit 1, mitochondrial | 1.81E-03 | -3.36 | NA | NA | NA | NA |
| P55072 | VCP | Transitional endoplasmic reticulum ATPase | 5.96E-06 | -3.35 | 2.26E-01 | -0.67 | 3.38E-01 | -0.47 |
| P21796 | VDAC1 | Voltage-dependent anion-selective channel protein 1 | 3.96E-03 | -3.33 | NA | NA | NA | NA |
| O00148;Q13838 | DDX39A;DDX39B | ATP-dependent RNA helicase DDX39A | 1.93E-05 | -3.32 | NA | NA | NA | NA |
| Q99733 | NAP1L4 | Nucleosome assembly protein 1-like 4 | 3.04E-05 | -3.32 | NA | NA | NA | NA |
| Q00325 | SLC25A3 | Phosphate carrier protein, mitochondrial | 5.48E-03 | -3.26 | 5.96E-01 | -0.26 | 1.12E-01 | -0.79 |
| P0DMV8;P0DMV9 | HSPA1A;HSPA1B | Heat shock 70 kDa protein 1A | 1.88E-06 | -3.26 | NA | NA | NA | NA |

| | | | | | | | | |
|---|---|---|---|---|---|---|---|---|
| Q15365 | PCBP1 | Poly(rC)-binding protein 1 | 5.07E-04 | -3.24 | NA | NA | NA | NA |
| Q6PGP7 | TTC37 | Tetratricopeptide repeat protein 37 | 2.79E-04 | -3.21 | NA | NA | NA | NA |
| Q9NZ01 | TECR | Very-long-chain enoyl-CoA reductase | 2.57E-03 | -3.19 | NA | NA | NA | NA |
| P53618 | COPB1 | Coatomer subunit beta | 3.53E-04 | -3.19 | NA | NA | NA | NA |
| P13073 | COX4I1 | Cytochrome c oxidase subunit 4 isoform 1, mitochondrial | 5.15E-03 | -3.18 | NA | NA | NA | NA |
| P11021;P11142;P54652 | HSPA2;HSPA5;HSPA8 | Endoplasmic reticulum chaperone BiP | 3.48E-06 | -3.17 | NA | NA | NA | NA |
| P46781 | RPS9 | 40S ribosomal protein S9 | 1.14E-03 | -3.13 | 1.03E-01 | -0.93 | 1.11E-01 | -1.08 |
| P39023 | RPL3 | 60S ribosomal protein L3 | 2.69E-04 | -3.13 | 1.36E-01 | -0.72 | 1.29E-01 | -1.01 |
| P53007 | SLC25A1 | Tricarboxylate transport protein, mitochondrial | 6.92E-04 | -3.08 | NA | NA | 5.21E-01 | -0.36 |
| P07900;P08238;Q58FF8 | HSP90AA1;HSP90AB1; HSP90AB2P | Heat shock protein HSP 90-alpha | 2.48E-04 | -3.06 | 1.32E-03 | -2.23 | 3.52E-05 | -3.30 |
| P12235 | SLC25A4 | ADP/ATP translocase 1 | 6.16E-04 | -3.06 | NA | NA | NA | NA |
| Q9Y512 | SAMM50 | Sorting and assembly machinery component 50 homolog | 2.77E-03 | -3.04 | NA | NA | NA | NA |
| P07437 | TUBB | Tubulin beta chain | 7.09E-04 | -3.04 | 7.74E-03 | -2.05 | NA | NA |
| P84243 | H3-3A | Histone H3.3 | 3.18E-03 | -3.02 | 3.37E-03 | -2.45 | NA | NA |
| P16104;Q8IUE6 | H2AC21;H2AX | Histone H2AX | 8.03E-03 | -3.02 | NA | NA | NA | NA |
| P24539 | ATP5PB | ATP synthase F(0) complex subunit B1, mitochondrial | 1.64E-03 | -3.00 | 8.54E-02 | -0.75 | 1.44E-04 | -2.24 |
| P43007 | SLC1A4 | Neutral amino acid transporter A | 2.53E-04 | -3.00 | NA | NA | NA | NA |
| O15020 | SPTBN2 | Spectrin beta chain, non-erythrocytic 2 | 3.88E-04 | -2.95 | 2.77E-02 | -1.14 | 4.11E-03 | -1.43 |
| P04350;P07437;P68371; Q13509;Q13885;Q9BUF5; Q9BVA1 | TUBB;TUBB2A;TUBB2B; TUBB3;TUBB4A;TUBB4 B;TUBB6 | Tubulin beta-4A chain | 1.91E-04 | -2.95 | NA | NA | NA | NA |
| P62906 | RPL10A | 60S ribosomal protein L10a | 3.50E-03 | -2.94 | NA | NA | NA | NA |
| P07900;P08238 | HSP90AA1;HSP90AB1 | Heat shock protein HSP 90-alpha | 5.15E-04 | -2.94 | 2.92E-01 | -0.27 | 2.53E-01 | -0.31 |
| P14923 | JUP | Junction plakoglobin | 7.97E-04 | -2.94 | NA | NA | NA | NA |
| P84243;Q5TEC6 | H3-2;H3-3A | Histone H3.3 | 3.72E-03 | -2.94 | NA | NA | NA | NA |
| P57053;Q16778 | H2BC21;H2BS1 | Histone H2B type F-S | 7.70E-03 | -2.93 | NA | NA | NA | NA |
| O76024 | WFS1 | Wolframin | 2.41E-04 | -2.91 | 2.58E-02 | -1.33 | 3.43E-03 | -1.39 |
| O00571;O15523 | DDX3X;DDX3Y | ATP-dependent RNA helicase DDX3X | 7.80E-04 | -2.90 | NA | NA | NA | NA |
| Q14697 | GANAB | Neutral alpha-glucosidase AB | 9.75E-04 | -2.90 | 5.83E-01 | 0.17 | 8.37E-01 | 0.08 |
| Q15366 | PCBP2 | Poly(rC)-binding protein 2 | 8.47E-04 | -2.86 | NA | NA | NA | NA |

| | | | | | | | | |
|---|---|---|---|---|---|---|---|---|
| Q15363 | TMED2 | Transmembrane emp24 domain-containing protein 2 | 6.72E-04 | -2.86 | NA | NA | NA | NA |
| Q15365;Q15366 | PCBP1;PCBP2 | Poly(rC)-binding protein 1 | 2.56E-06 | -2.83 | 1.85E-02 | -1.00 | 7.26E-01 | -0.12 |
| P17931 | LGALS3 | Galectin-3 | 5.02E-03 | -2.83 | NA | NA | NA | NA |
| P31946 | YWHAB | 14-3-3 protein beta/alpha | 9.55E-04 | -2.83 | NA | NA | NA | NA |
| P04350 | TUBB4A | Tubulin beta-4A chain | 2.42E-04 | -2.83 | 1.57E-02 | -2.28 | NA | NA |
| P04899 | GNAI2 | Guanine nucleotide-binding protein G(i) subunit alpha-2 | 3.06E-04 | -2.82 | NA | NA | NA | NA |
| P05141;P12235;P12236 | SLC25A4;SLC25A5;SLC25A6 | ADP/ATP translocase 2 | 6.67E-03 | -2.80 | 3.57E-01 | -0.45 | 3.52E-02 | -1.35 |
| P31948 | STIP1 | Stress-induced-phosphoprotein 1 | 5.74E-03 | -2.76 | NA | NA | NA | NA |
| P19022 | CDH2 | Cadherin-2 | 1.11E-03 | -2.73 | 4.77E-03 | -2.55 | 1.90E-03 | -3.06 |
| Q9BXM0 | PRX | Periaxin | 4.26E-04 | -2.71 | 2.27E-03 | -2.79 | 8.08E-04 | -2.50 |
| P62805 | H4C1 | Histone H4 | 4.89E-03 | -2.71 | 5.73E-02 | -1.35 | 1.29E-02 | -1.84 |
| P35579;P35749 | MYH11;MYH9 | Myosin-9 | 2.65E-03 | -2.71 | NA | NA | NA | NA |
| P55209 | NAP1L1 | Nucleosome assembly protein 1-like 1 | 8.54E-04 | -2.71 | 4.68E-02 | -1.30 | NA | NA |
| O75367;Q9P0M6 | MACROH2A1;MACROH2A2 | Core histone macro-H2A.1 | 4.62E-03 | -2.70 | NA | NA | NA | NA |
| P11142;P54652 | HSPA2;HSPA8 | Heat shock cognate 71 kDa protein | 3.30E-04 | -2.68 | 6.96E-03 | -1.55 | 1.89E-01 | -0.57 |
| P13489 | RNH1 | Ribonuclease inhibitor | 5.71E-03 | -2.67 | 1.30E-01 | -0.92 | 9.55E-03 | -1.47 |
| P27348 | YWHAQ | 14-3-3 protein theta | 3.68E-04 | -2.67 | 6.29E-02 | -0.89 | 4.05E-01 | -0.57 |
| Q7L099 | RUFY3 | Protein RUFY3 | 5.31E-04 | -2.66 | NA | NA | NA | NA |
| P07900;P08238;Q58FF6;Q58FF7;Q58FF8 | HSP90AA1;HSP90AB1;HSP90AB2P;HSP90AB3P;HSP90AB4P | Heat shock protein HSP 90-alpha | 3.05E-04 | -2.66 | NA | NA | NA | NA |
| P26232 | CTNNA2 | Catenin alpha-2 | 2.17E-03 | -2.65 | 1.17E-03 | -2.56 | 2.60E-04 | -1.71 |
| O00203 | AP3B1 | AP-3 complex subunit beta-1 | 8.48E-04 | -2.65 | NA | NA | NA | NA |
| O94874 | UFL1 | E3 UFM1-protein ligase 1 | 1.15E-04 | -2.65 | NA | NA | NA | NA |
| Q96HC4 | PDLIM5 | PDZ and LIM domain protein 5 | 6.04E-04 | -2.64 | NA | NA | NA | NA |
| P00450 | CP | Ceruloplasmin | 7.36E-07 | -2.63 | NA | NA | NA | NA |
| P04920 | SLC4A2 | Anion exchange protein 2 | 2.58E-04 | -2.62 | NA | NA | NA | NA |
| Q92597 | NDRG1 | Protein NDRG1 | 5.25E-03 | -2.62 | 7.94E-02 | -1.04 | 1.02E-01 | -1.22 |
| P51991 | HNRNPA3 | Heterogeneous nuclear ribonucleoprotein A3 | 2.54E-04 | -2.62 | NA | NA | NA | NA |
| P12236 | SLC25A6 | ADP/ATP translocase 3 | 6.01E-03 | -2.59 | NA | NA | NA | NA |

| | | | | | | | | |
|---|---|---|---|---|---|---|---|---|
| Q07960 | ARHGAP1 | Rho GTPase-activating protein 1 | 2.06E-03 | -2.59 | 5.43E-01 | 0.24 | 2.38E-01 | -0.59 |
| Q8IXS6 | PALM2 | Paralemmin-2 | 1.31E-04 | -2.57 | 3.35E-04 | -3.94 | NA | NA |
| Q02318 | CYP27A1 | Sterol 26-hydroxylase, mitochondrial | 6.02E-03 | -2.56 | NA | NA | NA | NA |
| Q04637 | EIF4G1 | Eukaryotic translation initiation factor 4 gamma 1 | 7.20E-05 | -2.56 | 1.07E-02 | -0.83 | 2.67E-01 | -0.49 |
| Q14974 | KPNB1 | Importin subunit beta-1 | 8.29E-04 | -2.56 | 1.55E-02 | -0.95 | 7.42E-03 | -1.39 |
| O00410 | IPO5 | Importin-5 | 2.06E-03 | -2.55 | 6.13E-02 | -0.81 | 4.88E-03 | -1.18 |
| P63027;Q15836 | VAMP2;VAMP3 | Vesicle-associated membrane protein 2 | 4.06E-04 | -2.54 | NA | NA | NA | NA |
| P49755 | TMED10 | Transmembrane emp24 domain-containing protein 10 | 6.38E-03 | -2.53 | NA | NA | NA | NA |
| Q12955 | ANK3 | Ankyrin-3 | 9.37E-05 | -2.53 | 4.91E-05 | -1.49 | 5.52E-03 | -1.63 |
| P55290 | CDH13 | Cadherin-13 | 8.39E-03 | -2.52 | NA | NA | NA | NA |
| P19086;P29992;P50148;Q14344 | GNA11;GNA13;GNAQ;GNAZ | Guanine nucleotide-binding protein G(z) subunit alpha | 3.28E-04 | -2.51 | NA | NA | NA | NA |
| O95747 | OXSR1 | Serine/threonine-protein kinase OSR1 | 4.79E-04 | -2.50 | NA | NA | NA | NA |
| P0DMV8;P0DMV9;P48741 | HSPA1A;HSPA1B;HSPA7 | Heat shock 70 kDa protein 1A | 1.09E-03 | -2.50 | 8.76E-01 | -0.10 | 1.39E-01 | -0.90 |
| Q9H853 | TUBA4B | Putative tubulin-like protein alpha-4B | 1.68E-04 | -2.49 | 1.19E-02 | -1.53 | 6.21E-04 | -1.26 |
| Q9H0A8 | COMMD4 | COMM domain-containing protein 4 | 8.15E-05 | -2.48 | NA | NA | NA | NA |
| O14880 | MGST3 | Microsomal glutathione S-transferase 3 | 8.90E-04 | -2.48 | NA | NA | 2.69E-01 | -0.74 |
| P04632 | CAPNS1 | Calpain small subunit 1 | 5.78E-04 | -2.48 | NA | NA | NA | NA |
| O43390 | HNRNPR | Heterogeneous nuclear ribonucleoprotein R | 4.28E-03 | -2.47 | NA | NA | NA | NA |
| O00519 | FAAH | Fatty-acid amide hydrolase 1 | 1.25E-03 | -2.47 | NA | NA | NA | NA |
| Q9H3Z4 | DNAJC5 | DnaJ homolog subfamily C member 5 | 1.45E-03 | -2.47 | NA | NA | NA | NA |
| P06396 | GSN | Gelsolin | 3.42E-04 | -2.47 | 3.41E-02 | -0.91 | 7.79E-01 | -0.12 |
| P21964 | COMT | Catechol O-methyltransferase | 1.40E-03 | -2.46 | 1.08E-02 | -1.56 | NA | NA |
| P57088 | TMEM33 | Transmembrane protein 33 | 4.89E-03 | -2.44 | NA | NA | NA | NA |
| P68366;Q9BQE3;Q9H853 | TUBA1C;TUBA4A;TUBA4B | Tubulin alpha-4A chain | 4.48E-05 | -2.44 | 2.48E-02 | -1.17 | 1.08E-03 | -2.19 |
| Q13492 | PICALM | Phosphatidylinositol-binding clathrin assembly protein | 1.30E-03 | -2.43 | NA | NA | NA | NA |
| P08238;P14625 | HSP90AB1;HSP90B1 | Heat shock protein HSP 90-beta | 5.34E-03 | -2.42 | 2.23E-02 | -1.27 | NA | NA |
| P04083 | ANXA1 | Annexin A1 | 6.26E-03 | -2.42 | 6.51E-01 | -0.25 | 1.20E-02 | -1.58 |
| Q5T4S7 | UBR4 | E3 ubiquitin-protein ligase UBR4 | 5.44E-05 | -2.42 | 1.94E-01 | -0.40 | 3.50E-01 | 0.25 |
| P35221 | CTNNA1 | Catenin alpha-1 | 7.61E-04 | -2.41 | NA | NA | NA | NA |

| | | | | | | | | |
|---|---|---|---|---|---|---|---|---|
| P53992 | SEC24C | Protein transport protein Sec24C | 2.63E-03 | -2.41 | NA | NA | NA | NA |
| P55209;Q99733 | NAP1L1;NAP1L4 | Nucleosome assembly protein 1-like 1 | 8.09E-04 | -2.40 | 8.34E-01 | -0.10 | 2.34E-01 | -0.60 |
| Q5T5P2 | KIAA1217 | Sickle tail protein homolog | 1.07E-03 | -2.40 | 4.43E-01 | -0.40 | NA | NA |
| P08195 | SLC3A2 | 4F2 cell-surface antigen heavy chain | 2.01E-03 | -2.40 | NA | NA | NA | NA |
| Q8NHG7 | SVIP | Small VCP/p97-interacting protein | 3.11E-04 | -2.40 | NA | NA | NA | NA |
| Q9C0E8 | LNPK | Endoplasmic reticulum junction formation protein lunapark | 1.01E-03 | -2.39 | NA | NA | NA | NA |
| O14949 | UQCRQ | Cytochrome b-c1 complex subunit 8 | 5.85E-03 | -2.38 | NA | NA | 8.99E-02 | -0.94 |
| P25686 | DNAJB2 | DnaJ homolog subfamily B member 2 | 3.60E-03 | -2.37 | 6.45E-02 | -1.35 | NA | NA |
| O96000 | NDUFB10 | NADH dehydrogenase [ubiquinone] 1 beta subcomplex subunit 10 | 9.31E-03 | -2.36 | NA | NA | NA | NA |
| O60282;P33176;Q12840 | KIF5A;KIF5B;KIF5C | Kinesin heavy chain isoform 5C | 1.96E-03 | -2.36 | NA | NA | NA | NA |
| P07900 | HSP90AA1 | Heat shock protein HSP 90-alpha | 2.75E-04 | -2.36 | 3.05E-03 | -1.13 | 4.93E-04 | -1.70 |
| P61019 | RAB2A | Ras-related protein Rab-2A | 5.16E-04 | -2.36 | NA | NA | NA | NA |
| Q16643 | DBN1 | Drebrin | 3.33E-03 | -2.35 | NA | NA | NA | NA |
| Q8N126 | CADM3 | Cell adhesion molecule 3 | 1.41E-04 | -2.35 | 1.97E-03 | -2.95 | 1.14E-02 | -1.23 |
| Q7Z406 | MYH14 | Myosin-14 | 1.11E-04 | -2.34 | 1.03E-03 | -1.20 | 8.28E-04 | -1.55 |
| O15031 | PLXNB2 | Plexin-B2 | 6.00E-03 | -2.34 | NA | NA | NA | NA |
| P30153 | PPP2R1A | Serine/threonine-protein phosphatase 2A 65 kDa regulatory subunit A alpha isoform | 3.73E-03 | -2.32 | 2.28E-04 | -1.19 | 5.74E-03 | -1.57 |
| O43390;O60506 | HNRNPR;SYNCRIP | Heterogeneous nuclear ribonucleoprotein R | 4.90E-03 | -2.31 | NA | NA | 6.62E-02 | -1.05 |
| Q02978 | SLC25A11 | Mitochondrial 2-oxoglutarate/malate carrier protein | 4.64E-03 | -2.31 | 9.57E-01 | -0.02 | 3.61E-01 | -0.43 |
| O75964 | ATP5MG | ATP synthase subunit g, mitochondrial | 5.20E-03 | -2.29 | 3.18E-01 | 0.47 | 2.57E-01 | -0.54 |
| Q9NQT8 | KIF13B | Kinesin-like protein KIF13B | 2.53E-05 | -2.29 | NA | NA | NA | NA |
| P46821 | MAP1B | Microtubule-associated protein 1B | 1.58E-04 | -2.29 | NA | NA | NA | NA |
| P53794 | SLC5A3 | Sodium/myo-inositol cotransporter | 5.11E-03 | -2.28 | NA | NA | NA | NA |
| P78310 | CXADR | Coxsackievirus and adenovirus receptor | 1.17E-03 | -2.27 | NA | NA | NA | NA |
| Q9NZJ7 | MTCH1 | Mitochondrial carrier homolog 1 | 7.70E-03 | -2.27 | 5.00E-01 | 0.36 | 5.29E-01 | -0.43 |
| O60664 | PLIN3 | Perilipin-3 | 6.96E-04 | -2.26 | 8.25E-03 | -1.66 | 7.97E-03 | -1.45 |
| P53675;Q00610 | CLTC;CLTCL1 | Clathrin heavy chain 2 | 4.26E-03 | -2.25 | 1.20E-02 | -1.06 | 7.92E-05 | -2.12 |
| P35580 | MYH10 | Myosin-10 | 5.89E-04 | -2.25 | NA | NA | NA | NA |
| P13861 | PRKAR2A | cAMP-dependent protein kinase type II-alpha regulatory subunit | 1.23E-04 | -2.24 | NA | NA | NA | NA |

| | | | | | | | | |
|---|---|---|---|---|---|---|---|---|
| P32856 | STX2 | Syntaxin-2 | 1.78E-04 | -2.23 | NA | NA | NA | NA |
| P11117 | ACP2 | Lysosomal acid phosphatase | 9.84E-03 | -2.23 | NA | NA | NA | NA |
| P62820 | RAB1A | Ras-related protein Rab-1A | 3.08E-04 | -2.21 | 1.53E-03 | -2.34 | 3.74E-04 | -3.02 |
| P63218 | GNG5 | Guanine nucleotide-binding protein G(I)/G(S)/G(O) subunit gamma-5 | 1.50E-04 | -2.21 | 7.39E-04 | -4.38 | NA | NA |
| Q00610 | CLTC | Clathrin heavy chain 1 | 2.24E-03 | -2.21 | 2.52E-03 | -0.82 | 3.79E-01 | -0.27 |
| P07384 | CAPN1 | Calpain-1 catalytic subunit | 6.97E-03 | -2.21 | NA | NA | NA | NA |
| Q6XZF7 | DNMBP | Dynamin-binding protein | 3.82E-04 | -2.21 | 2.51E-01 | -0.63 | NA | NA |
| Q92734 | TFG | Protein TFG | 5.69E-06 | -2.21 | NA | NA | 6.07E-02 | -1.41 |
| O14787;Q92973 | TNPO1;TNPO2 | Transportin-2 | 2.65E-04 | -2.20 | 8.45E-03 | -0.72 | 1.02E-01 | 0.83 |
| P06213 | INSR | Insulin receptor | 2.65E-03 | -2.19 | NA | NA | NA | NA |
| O75369;P21333;Q14315 | FLNA;FLNB;FLNC | Filamin-B | 2.52E-04 | -2.19 | 1.01E-03 | -2.13 | NA | NA |
| P08574 | CYC1 | Cytochrome c1, heme protein, mitochondrial | 1.51E-03 | -2.19 | 1.14E-01 | -0.69 | 3.05E-03 | -2.43 |
| Q92973 | TNPO1 | Transportin-1 | 1.19E-03 | -2.19 | 8.81E-02 | -0.64 | 3.23E-02 | -1.07 |
| P35579 | MYH9 | Myosin-9 | 2.35E-03 | -2.19 | 3.59E-01 | -0.24 | 7.64E-02 | -0.68 |
| P54709 | ATP1B3 | Sodium/potassium-transporting ATPase subunit beta-3 | 7.98E-03 | -2.17 | 4.12E-02 | -1.90 | NA | NA |
| O75695 | RP2 | Protein XRP2 | 1.80E-04 | -2.17 | 3.20E-04 | -2.75 | 1.26E-02 | -2.00 |
| P12956 | XRCC6 | X-ray repair cross-complementing protein 6 | 2.72E-04 | -2.17 | 5.69E-01 | -0.16 | 3.70E-03 | -1.43 |
| P61764 | STXBP1 | Syntaxin-binding protein 1 | 3.13E-04 | -2.16 | 6.06E-01 | 0.24 | 5.04E-02 | -1.36 |
| Q96FZ7 | CHMP6 | Charged multivesicular body protein 6 | 4.40E-03 | -2.16 | NA | NA | NA | NA |
| Q96QR8 | PURB | Transcriptional activator protein Pur-beta | 2.77E-04 | -2.15 | 8.07E-01 | -0.10 | 1.69E-04 | -1.84 |
| Q14008 | CKAP5 | Cytoskeleton-associated protein 5 | 1.49E-03 | -2.15 | NA | NA | NA | NA |
| P01111;P01112 | HRAS;NRAS | GTPase NRas | 5.01E-03 | -2.15 | NA | NA | NA | NA |
| P63010 | AP2B1 | AP-2 complex subunit beta | 1.19E-03 | -2.14 | 6.33E-01 | -0.18 | 1.52E-02 | -1.31 |
| O43707 | ACTN4 | Alpha-actinin-4 | 1.82E-03 | -2.13 | 9.88E-01 | -0.01 | 2.19E-03 | -1.73 |
| P29992;P50148 | GNA11;GNAQ | Guanine nucleotide-binding protein subunit alpha-11 | 3.45E-04 | -2.12 | NA | NA | NA | NA |
| Q12846 | STX4 | Syntaxin-4 | 2.44E-03 | -2.12 | NA | NA | NA | NA |
| P61020 | RAB5B | Ras-related protein Rab-5B | 2.92E-03 | -2.12 | 1.18E-03 | -2.43 | NA | NA |
| P21333;Q14315 | FLNA;FLNC | Filamin-A | 1.11E-04 | -2.12 | 1.68E-02 | -1.66 | 7.64E-04 | -1.94 |
| Q12905 | ILF2 | Interleukin enhancer-binding factor 2 | 3.78E-03 | -2.11 | 2.06E-01 | -0.59 | 1.24E-01 | -1.17 |
| Q9NZW5 | PALS2 | Protein PALS2 | 2.16E-03 | -2.11 | NA | NA | NA | NA |

| | | | | | | | | |
|---|---|---|---|---|---|---|---|---|
| P35222 | CTNNB1 | Catenin beta-1 | 1.66E-03 | -2.11 | 2.74E-02 | -1.34 | 4.18E-03 | -1.45 |
| Q86VP6 | CAND1 | Cullin-associated NEDD8-dissociated protein 1 | 1.04E-03 | -2.10 | 7.50E-05 | -1.29 | 6.91E-04 | -1.79 |
| P09543 | CNP | 2',3'-cyclic-nucleotide 3'-phosphodiesterase | 3.77E-04 | -2.10 | NA | NA | NA | NA |
| Q14203 | DCTN1 | Dynactin subunit 1 | 4.74E-04 | -2.10 | 4.44E-02 | -0.79 | 7.45E-03 | -1.83 |
| Q14194;Q14195;Q16555 | CRMP1;DPYSL2;DPYSL3 | Dihydropyrimidinase-related protein 1 | 5.72E-04 | -2.09 | NA | NA | NA | NA |
| Q9NZU5 | LMCD1 | LIM and cysteine-rich domains protein 1 | 6.44E-03 | -2.08 | 5.90E-02 | -1.03 | 1.48E-01 | -0.86 |
| P11277 | SPTB | Spectrin beta chain, erythrocytic | 2.03E-04 | -2.06 | 2.45E-03 | -1.51 | 1.54E-04 | -2.32 |
| P61106 | RAB14 | Ras-related protein Rab-14 | 1.29E-04 | -2.06 | 2.63E-03 | -1.74 | 5.21E-03 | -1.65 |
| P00533 | EGFR | Epidermal growth factor receptor | 5.05E-03 | -2.05 | NA | NA | NA | NA |
| P26447 | S100A4 | Protein S100-A4 | 4.78E-03 | -2.05 | 5.58E-01 | 0.33 | 5.85E-01 | -0.24 |
| O00505;O00629 | KPNA3;KPNA4 | Importin subunit alpha-4 | 2.93E-03 | -2.05 | 7.76E-01 | 0.07 | 4.56E-04 | -1.70 |
| P11277;Q01082 | SPTB;SPTBN1 | Spectrin beta chain, erythrocytic | 1.34E-03 | -2.04 | 4.48E-02 | -1.13 | NA | NA |
| P55060 | CSE1L | Exportin-2 | 1.69E-03 | -2.04 | 1.01E-01 | -0.57 | 3.96E-03 | -1.49 |
| Q13449 | LSAMP | Limbic system-associated membrane protein | 7.73E-04 | -2.03 | 9.83E-04 | -2.81 | 2.50E-04 | -2.36 |
| P21333 | FLNA | Filamin-A | 6.55E-04 | -2.02 | 4.40E-02 | -0.98 | 4.00E-01 | -0.28 |
| Q93050 | ATP6V0A1 | V-type proton ATPase 116 kDa subunit a1 | 8.32E-04 | -2.02 | NA | NA | NA | NA |
| Q13557 | CAMK2D | Calcium/calmodulin-dependent protein kinase type II subunit delta | 3.59E-03 | -2.02 | NA | NA | NA | NA |
| P05023 | ATP1A1 | Sodium/potassium-transporting ATPase subunit alpha-1 | 7.79E-04 | -2.01 | 9.83E-04 | -3.04 | NA | NA |
| P50570;Q9UQ16 | DNM2;DNM3 | Dynamin-2 | 3.62E-04 | -2.01 | NA | NA | NA | NA |
| P46783 | RPS10 | 40S ribosomal protein S10 | 9.27E-03 | -2.01 | NA | NA | NA | NA |
| Q9HBH5 | RDH14 | Retinol dehydrogenase 14 | 8.86E-03 | -2.01 | NA | NA | NA | NA |
| P11142 | HSPA8 | Heat shock cognate 71 kDa protein | 5.20E-04 | -2.00 | 1.68E-04 | -1.95 | 8.38E-04 | -1.88 |
| Q13162 | PRDX4 | Peroxiredoxin-4 | 1.79E-03 | -1.99 | 2.76E-01 | -0.75 | 7.09E-01 | 0.20 |
| Q6ZUX7 | LHFPL2 | LHFPL tetraspan subfamily member 2 protein | 1.50E-03 | -1.98 | NA | NA | NA | NA |
| O76003 | GLRX3 | Glutaredoxin-3 | 4.43E-03 | -1.98 | NA | NA | NA | NA |
| O60437 | PPL | Periplakin | 2.53E-04 | -1.97 | 8.38E-02 | -1.05 | 4.57E-02 | -1.56 |
| Q00577 | PURA | Transcriptional activator protein Pur-alpha | 1.20E-03 | -1.97 | 8.66E-02 | -0.69 | 4.12E-02 | -1.13 |
| Q01484;Q12955 | ANK2;ANK3 | Ankyrin-2 | 1.72E-03 | -1.97 | 5.97E-02 | -0.71 | 1.32E-02 | -1.02 |
| P17980 | PSMC3 | 26S proteasome regulatory subunit 6A | 1.33E-03 | -1.97 | NA | NA | NA | NA |
| Q9UNF1 | MAGED2 | Melanoma-associated antigen D2 | 9.54E-04 | -1.96 | 6.26E-03 | -1.85 | 8.66E-02 | -1.15 |

| | | | | | | | | |
|---|---|---|---|---|---|---|---|---|
| Q13443 | ADAM9 | Disintegrin and metalloproteinase domain-containing protein 9 | 7.12E-05 | -1.96 | NA | NA | NA | NA |
| Q93008 | USP9X | Probable ubiquitin carboxyl-terminal hydrolase FAF-X | 4.27E-03 | -1.96 | 7.95E-01 | 0.09 | 4.61E-01 | -0.38 |
| P26232;P35221 | CTNNA1;CTNNA2 | Catenin alpha-2 | 7.11E-03 | -1.95 | 7.04E-03 | -1.64 | 1.21E-01 | -0.67 |
| Q9BTU6 | PI4K2A | Phosphatidylinositol 4-kinase type 2-alpha | 7.15E-03 | -1.95 | NA | NA | NA | NA |
| P10301 | RRAS | Ras-related protein R-Ras | 3.44E-04 | -1.94 | NA | NA | NA | NA |
| P53985 | SLC16A1 | Monocarboxylate transporter 1 | 7.46E-03 | -1.94 | NA | NA | NA | NA |
| P62330 | ARF6 | ADP-ribosylation factor 6 | 2.65E-03 | -1.94 | NA | NA | NA | NA |
| P62826 | RAN | GTP-binding nuclear protein Ran | 6.24E-06 | -1.94 | 1.16E-01 | -0.74 | 5.01E-01 | -0.21 |
| Q5VYK3 | ECPAS | Proteasome adapter and scaffold protein ECM29 | 3.74E-03 | -1.94 | NA | NA | NA | NA |
| Q96LR9 | APOLD1 | Apolipoprotein L domain-containing protein 1 | 2.83E-04 | -1.94 | NA | NA | NA | NA |
| P20042 | EIF2S2 | Eukaryotic translation initiation factor 2 subunit 2 | 5.43E-03 | -1.93 | NA | NA | NA | NA |
| Q01484 | ANK2 | Ankyrin-2 | 2.56E-03 | -1.92 | 1.52E-02 | -1.15 | 1.46E-04 | -1.91 |
| Q14964 | RAB39A | Ras-related protein Rab-39A | 2.90E-04 | -1.92 | NA | NA | NA | NA |
| Q6IAA8 | LAMTOR1 | Ragulator complex protein LAMTOR1 | 3.77E-05 | -1.92 | 2.29E-04 | -1.59 | 1.23E-02 | -1.35 |
| P54289 | CACNA2D1 | Voltage-dependent calcium channel subunit alpha-2/delta-1 | 1.84E-03 | -1.91 | NA | NA | NA | NA |
| O15020;Q01082 | SPTBN1;SPTBN2 | Spectrin beta chain, non-erythrocytic 2 | 3.51E-03 | -1.90 | 2.89E-01 | -0.44 | 1.76E-03 | -1.23 |
| Q9NZH0 | GPRC5B | G-protein coupled receptor family C group 5 member B | 2.72E-03 | -1.90 | NA | NA | NA | NA |
| Q99436 | PSMB7 | Proteasome subunit beta type-7 | 1.68E-03 | -1.90 | NA | NA | NA | NA |
| Q96CS3 | FAF2 | FAS-associated factor 2 | 6.38E-04 | -1.90 | 8.69E-01 | -0.08 | NA | NA |
| Q14204 | DYNC1H1 | Cytoplasmic dynein 1 heavy chain 1 | 6.35E-04 | -1.89 | 8.63E-02 | -0.53 | 1.88E-02 | -1.06 |
| P18084 | ITGB5 | Integrin beta-5 | 5.62E-03 | -1.88 | NA | NA | NA | NA |
| Q9H0E2 | TOLLIP | Toll-interacting protein | 8.71E-04 | -1.88 | 4.82E-04 | -1.99 | NA | NA |
| P84085 | ARF5 | ADP-ribosylation factor 5 | 3.08E-04 | -1.88 | 1.81E-02 | -0.99 | 1.48E-03 | -1.78 |
| P62495 | ETF1 | Eukaryotic peptide chain release factor subunit 1 | 1.36E-04 | -1.88 | 8.23E-01 | -0.09 | 5.25E-01 | -0.26 |
| P20339;P51148;P61020 | RAB5A;RAB5B;RAB5C | Ras-related protein Rab-5A | 1.33E-03 | -1.88 | 8.83E-02 | -0.92 | 3.13E-02 | -1.05 |
| P01111;P01112;P01116 | HRAS;KRAS;NRAS | GTPase NRas | 1.45E-03 | -1.87 | 6.06E-02 | -0.99 | NA | NA |
| P07355 | ANXA2 | Annexin A2 | 2.58E-03 | -1.86 | 2.58E-02 | -1.11 | 4.79E-03 | -1.73 |
| Q15149 | PLEC | Plectin | 9.10E-04 | -1.86 | 4.30E-03 | -1.16 | 1.04E-02 | -1.33 |
| Q9NZJ9 | NUDT4 | Diphosphoinositol polyphosphate phosphohydrolase 2 | 1.50E-05 | -1.86 | 6.08E-01 | -0.22 | 1.50E-03 | -2.17 |

| | | | | | | | | |
|---|---|---|---|---|---|---|---|---|
| P11717 | IGF2R | Cation-independent mannose-6-phosphate receptor | 6.63E-03 | -1.85 | NA | NA | NA | NA |
| P23396 | RPS3 | 40S ribosomal protein S3 | 1.13E-03 | -1.85 | 3.01E-01 | 0.38 | 3.82E-01 | 0.44 |
| P01116 | KRAS | GTPase KRas | 1.12E-03 | -1.85 | 1.92E-03 | -2.13 | 7.45E-04 | -1.70 |
| P55011 | SLC12A2 | Solute carrier family 12 member 2 | 7.05E-03 | -1.83 | 6.65E-03 | -2.52 | NA | NA |
| P61019;Q8WUD1 | RAB2A;RAB2B | Ras-related protein Rab-2A | 1.14E-04 | -1.83 | 6.46E-03 | -1.42 | 1.71E-02 | -1.36 |
| Q58EX2 | SDK2 | Protein sidekick-2 | 1.07E-03 | -1.83 | NA | NA | NA | NA |
| Q9NP72 | RAB18 | Ras-related protein Rab-18 | 2.41E-03 | -1.81 | 1.86E-02 | -1.21 | 9.18E-02 | -0.74 |
| Q8TD20 | SLC2A12 | Solute carrier family 2, facilitated glucose transporter member 12 | 1.66E-03 | -1.81 | 2.57E-02 | -1.83 | NA | NA |
| P33527 | ABCC1 | Multidrug resistance-associated protein 1 | 3.82E-03 | -1.81 | NA | NA | NA | NA |
| P53621 | COPA | Coatomer subunit alpha | 3.54E-03 | -1.80 | 9.72E-01 | -0.01 | NA | NA |
| P35579;P35580;P35749;Q7Z406 | MYH10;MYH11;MYH14;MYH9 | Myosin-9 | 4.12E-03 | -1.79 | NA | NA | 2.77E-02 | -1.48 |
| P02786 | TFRC | Transferrin receptor protein 1 | 3.17E-03 | -1.78 | NA | NA | NA | NA |
| O75781 | PALM | Paralemmin-1 | 3.72E-03 | -1.78 | 1.22E-02 | -1.61 | 4.29E-04 | -1.52 |
| P05023;P13637 | ATP1A1;ATP1A3 | Sodium/potassium-transporting ATPase subunit alpha-1 | 1.43E-04 | -1.77 | 9.53E-03 | -2.49 | NA | NA |
| Q9H0U4 | RAB1B | Ras-related protein Rab-1B | 7.51E-03 | -1.77 | 1.08E-03 | -2.75 | NA | NA |
| Q9UL25 | RAB21 | Ras-related protein Rab-21 | 6.56E-03 | -1.77 | 3.78E-03 | -1.72 | 1.33E-02 | -1.24 |
| Q9Y6G9 | DYNC1LI1 | Cytoplasmic dynein 1 light intermediate chain 1 | 4.20E-03 | -1.76 | 9.71E-01 | -0.02 | NA | NA |
| P05023;P13637;P50993 | ATP1A1;ATP1A2;ATP1A3 | Sodium/potassium-transporting ATPase subunit alpha-1 | 1.83E-03 | -1.76 | 4.67E-03 | -2.45 | NA | NA |
| Q13619;Q13620 | CUL4A;CUL4B | Cullin-4A | 9.36E-04 | -1.76 | 4.34E-01 | -0.22 | 2.03E-01 | -0.63 |
| P04899;P08754;P11488;P63096 | GNAI1;GNAI2;GNAI3;GNAT1 | Guanine nucleotide-binding protein G(i) subunit alpha-2 | 4.93E-04 | -1.76 | NA | NA | NA | NA |
| P08238;Q58FF7 | HSP90AB1;HSP90AB3P | Heat shock protein HSP 90-beta | 2.85E-03 | -1.75 | NA | NA | NA | NA |
| Q7L9L4;Q9H8S9 | MOB1A;MOB1B | MOB kinase activator 1B | 5.22E-04 | -1.73 | 1.29E-01 | -0.64 | 1.10E-02 | -1.91 |
| O43491 | EPB41L2 | Band 4.1-like protein 2 | 1.15E-07 | -1.73 | 2.80E-03 | -1.75 | 1.33E-03 | -2.24 |
| Q8TAT6 | NPLOC4 | Nuclear protein localization protein 4 homolog | 1.63E-04 | -1.71 | NA | NA | 3.66E-02 | -0.98 |
| P08237 | PFKM | ATP-dependent 6-phosphofructokinase, muscle type | 1.84E-03 | -1.71 | 5.75E-02 | -0.82 | 4.91E-01 | -0.44 |
| P12814 | ACTN1 | Alpha-actinin-1 | 3.95E-03 | -1.71 | 1.67E-01 | -0.58 | 6.81E-02 | -0.92 |
| P26641 | EEF1G | Elongation factor 1-gamma | 5.50E-03 | -1.71 | 1.25E-01 | -0.55 | 3.69E-01 | -0.39 |
| O43681 | GET3 | ATPase GET3 | 3.45E-03 | -1.70 | NA | NA | 4.42E-02 | -0.96 |

| | | | | | | | | |
|---|---|---|---|---|---|---|---|---|
| O00192 | ARVCF | Armadillo repeat protein deleted in velo-cardio-facial syndrome | 8.97E-04 | -1.70 | 8.42E-03 | -1.75 | 1.52E-05 | -2.06 |
| Q68EM7 | ARHGAP17 | Rho GTPase-activating protein 17 | 5.42E-04 | -1.70 | 2.55E-01 | -0.55 | NA | NA |
| O43795 | MYO1B | Unconventional myosin-Ib | 1.81E-03 | -1.69 | 3.34E-02 | -0.89 | 7.87E-03 | -1.34 |
| P20339;P61020 | RAB5A;RAB5B | Ras-related protein Rab-5A | 1.82E-03 | -1.69 | NA | NA | NA | NA |
| Q7L576 | CYFIP1 | Cytoplasmic FMR1-interacting protein 1 | 3.08E-03 | -1.69 | NA | NA | NA | NA |
| O60763 | USO1 | General vesicular transport factor p115 | 6.92E-03 | -1.68 | 2.80E-01 | -0.43 | 2.69E-02 | -1.09 |
| P14923;P35222 | CTNNB1;JUP | Junction plakoglobin | 9.72E-03 | -1.68 | 1.04E-01 | -0.83 | 4.61E-02 | -0.71 |
| P43353 | ALDH3B1 | Aldehyde dehydrogenase family 3 member B1 | 4.06E-03 | -1.68 | NA | NA | NA | NA |
| P13591 | NCAM1 | Neural cell adhesion molecule 1 | 4.83E-03 | -1.67 | 1.95E-04 | -2.13 | 1.08E-02 | -1.89 |
| O60749;Q13596 | SNX1;SNX2 | Sorting nexin-2 | 4.30E-03 | -1.67 | NA | NA | NA | NA |
| Q8WVC6 | DCAKD | Dephospho-CoA kinase domain-containing protein | 4.68E-03 | -1.67 | 4.96E-03 | 0.92 | 6.28E-01 | -0.18 |
| P13473 | LAMP2 | Lysosome-associated membrane glycoprotein 2 | 1.41E-03 | -1.67 | NA | NA | NA | NA |
| P61026 | RAB10 | Ras-related protein Rab-10 | 1.99E-04 | -1.67 | 1.18E-03 | -1.84 | 2.54E-03 | -1.66 |
| Q13617 | CUL2 | Cullin-2 | 7.46E-05 | -1.66 | 1.19E-01 | -0.49 | 1.72E-01 | -0.53 |
| O43707;P12814 | ACTN1;ACTN4 | Alpha-actinin-4 | 3.76E-03 | -1.65 | 3.56E-01 | 0.39 | 9.79E-02 | -0.85 |
| Q9Y2Z0 | SUGT1 | Protein SGT1 homolog | 1.24E-03 | -1.64 | NA | NA | NA | NA |
| P51149 | RAB7A | Ras-related protein Rab-7a | 1.84E-03 | -1.63 | 1.77E-03 | -2.30 | 4.57E-04 | -2.42 |
| P42224 | STAT1 | Signal transducer and activator of transcription 1-alpha/beta | 3.57E-03 | -1.63 | 2.31E-01 | 0.55 | 3.05E-01 | -0.62 |
| Q96CW1 | AP2M1 | AP-2 complex subunit mu | 1.91E-03 | -1.62 | 3.85E-02 | -0.75 | 3.37E-02 | -0.73 |
| P05023;P13637;P50993;Q13733 | ATP1A1;ATP1A2;ATP1A3;ATP1A4 | Sodium/potassium-transporting ATPase subunit alpha-1 | 2.65E-03 | -1.62 | 5.72E-03 | -2.67 | NA | NA |
| P07900;P08238;Q58FF7 | HSP90AA1;HSP90AB1;HSP90AB3P | Heat shock protein HSP 90-alpha | 1.34E-03 | -1.61 | NA | NA | NA | NA |
| P61204;P84077 | ARF1;ARF3 | ADP-ribosylation factor 3 | 9.00E-04 | -1.61 | 6.98E-02 | -0.60 | 2.10E-02 | -1.07 |
| Q92729 | PTPRU | Receptor-type tyrosine-protein phosphatase U | 3.82E-03 | -1.61 | 1.05E-01 | -0.83 | NA | NA |
| P21926 | CD9 | CD9 antigen | 6.99E-03 | -1.61 | NA | NA | NA | NA |
| P11234 | RALB | Ras-related protein Ral-B | 2.85E-04 | -1.60 | 4.91E-03 | -1.68 | 3.47E-02 | -0.95 |
| O60331 | PIP5K1C | Phosphatidylinositol 4-phosphate 5-kinase type-1 gamma | 4.78E-03 | -1.59 | 1.32E-02 | -1.44 | NA | NA |
| P01112 | HRAS | GTPase HRas | 9.95E-03 | -1.59 | NA | NA | NA | NA |
| Q8N3E9 | PLCD3 | 1-phosphatidylinositol 4,5-bisphosphate phosphodiesterase delta-3 | 1.86E-04 | -1.59 | 4.35E-03 | -1.34 | NA | NA |
| P48165 | GJA8 | Gap junction alpha-8 protein | 1.33E-04 | -1.57 | 6.19E-04 | -1.53 | 8.35E-04 | -2.48 |

| | | | | | | | | |
|---|---|---|---|---|---|---|---|---|
| Q13555 | CAMK2G | Calcium/calmodulin-dependent protein kinase type II subunit gamma | 8.23E-03 | -1.57 | 1.55E-02 | -1.06 | NA | NA |
| P43490 | NAMPT | Nicotinamide phosphoribosyltransferase | 1.10E-04 | -1.57 | 1.90E-01 | -0.58 | NA | NA |
| O43865;Q96HN2 | AHCYL1;AHCYL2 | S-adenosylhomocysteine hydrolase-like protein 1 | 8.75E-03 | -1.56 | NA | NA | NA | NA |
| P61225 | RAP2B | Ras-related protein Rap-2b | 3.41E-03 | -1.56 | 1.96E-02 | -1.96 | NA | NA |
| P61224;P62834 | RAP1A;RAP1B | Ras-related protein Rap-1b | 7.36E-04 | -1.56 | 5.82E-02 | -0.88 | 2.05E-02 | -1.16 |
| Q8NCW6 | GALNT11 | Polypeptide N-acetylgalactosaminyltransferase 11 | 1.23E-03 | -1.56 | 4.08E-02 | -1.07 | 5.05E-03 | -2.13 |
| O00159 | MYO1C | Unconventional myosin-Ic | 1.42E-03 | -1.56 | 8.41E-02 | -0.69 | 5.74E-03 | -1.51 |
| P63096 | GNAI1 | Guanine nucleotide-binding protein G(i) subunit alpha-1 | 1.16E-03 | -1.56 | NA | NA | NA | NA |
| Q14152 | EIF3A | Eukaryotic translation initiation factor 3 subunit A | 6.79E-03 | -1.55 | 3.70E-02 | -0.83 | 1.11E-02 | -1.16 |
| Q9Y3S1 | WNK2 | Serine/threonine-protein kinase WNK2 | 2.37E-04 | -1.54 | 8.73E-01 | -0.11 | 3.01E-02 | -0.98 |
| P26196 | DDX6 | Probable ATP-dependent RNA helicase DDX6 | 6.24E-03 | -1.54 | NA | NA | NA | NA |
| O00231 | PSMD11 | 26S proteasome non-ATPase regulatory subunit 11 | 5.90E-03 | -1.54 | 3.17E-02 | -0.76 | 2.46E-04 | -1.84 |
| Q6IQ22 | RAB12 | Ras-related protein Rab-12 | 1.89E-03 | -1.54 | 1.56E-03 | -2.04 | 3.49E-02 | -1.66 |
| P63010;Q10567 | AP1B1;AP2B1 | AP-2 complex subunit beta | 9.65E-03 | -1.54 | 4.65E-01 | -0.19 | 2.65E-04 | -1.98 |
| Q13555;Q13557;Q9UQM7 | CAMK2A;CAMK2D;CAMK2G | Calcium/calmodulin-dependent protein kinase type II subunit gamma | 9.76E-04 | -1.54 | 9.09E-07 | -2.51 | 7.79E-04 | -3.63 |
| Q14964;Q96DA2 | RAB39A;RAB39B | Ras-related protein Rab-39A | 5.25E-05 | -1.53 | NA | NA | NA | NA |
| O94973 | AP2A2 | AP-2 complex subunit alpha-2 | 1.06E-03 | -1.52 | 9.18E-02 | -0.56 | 7.69E-02 | -0.76 |
| Q9HA65 | TBC1D17 | TBC1 domain family member 17 | 3.74E-03 | -1.52 | 2.17E-01 | -0.40 | 1.28E-01 | -0.55 |
| P15260 | IFNGR1 | Interferon gamma receptor 1 | 6.63E-04 | -1.51 | 4.61E-03 | -1.19 | NA | NA |
| Q13045 | FLII | Protein flightless-1 homolog | 7.25E-03 | -1.51 | 8.61E-01 | 0.06 | NA | NA |
| P35579;P35580 | MYH10;MYH9 | Myosin-9 | 6.79E-03 | -1.51 | 2.12E-02 | -0.95 | 2.00E-04 | -1.71 |
| P54136 | RARS1 | Arginine--tRNA ligase, cytoplasmic | 1.61E-03 | -1.51 | 3.48E-01 | -0.33 | 1.98E-02 | -0.94 |
| Q01968 | OCRL | Inositol polyphosphate 5-phosphatase OCRL | 9.34E-03 | -1.50 | NA | NA | NA | NA |
| Q9Y2J2 | EPB41L3 | Band 4.1-like protein 3 | 5.48E-06 | -1.49 | 1.62E-02 | -1.50 | 1.57E-03 | -1.99 |
| P63092;Q5JWF2 | GNAS | Guanine nucleotide-binding protein G(s) subunit alpha isoforms short | 1.02E-03 | -1.49 | 3.62E-03 | -2.03 | 2.08E-03 | -1.67 |
| Q8N0X7 | SPART | Spartin | 3.03E-03 | -1.48 | 1.51E-03 | -1.67 | NA | NA |
| O14576 | DYNC1I1 | Cytoplasmic dynein 1 intermediate chain 1 | 8.57E-04 | -1.48 | 8.02E-01 | -0.11 | 7.65E-03 | -1.50 |
| Q8TCT0 | CERK | Ceramide kinase | 6.09E-03 | -1.45 | 7.63E-05 | -2.07 | 5.67E-04 | -2.35 |
| Q9NV96 | TMEM30A | Cell cycle control protein 50A | 6.51E-03 | -1.45 | 2.39E-03 | -1.50 | 9.04E-03 | -1.91 |

| | | | | | | | | |
|---|---|---|---|---|---|---|---|---|
| O15498 | YKT6 | Synaptobrevin homolog YKT6 | 2.45E-03 | -1.44 | 6.57E-02 | -1.16 | 2.05E-03 | -1.97 |
| P62820;Q9H0U4 | RAB1A;RAB1B | Ras-related protein Rab-1A | 5.82E-03 | -1.43 | 1.98E-03 | -1.92 | 2.09E-03 | -1.15 |
| O14818 | PSMA7 | Proteasome subunit alpha type-7 | 3.83E-05 | -1.43 | 3.40E-02 | -0.72 | 4.74E-03 | -1.91 |
| Q99426 | TBCB | Tubulin-folding cofactor B | 2.53E-03 | -1.43 | 1.02E-04 | -1.59 | 1.15E-03 | -1.92 |
| P11171;Q9Y2J2 | EPB41;EPB41L3 | Protein 4.1 | 3.60E-03 | -1.42 | 9.76E-03 | -1.98 | 6.30E-02 | -1.00 |
| Q9NX46 | ADPRS | ADP-ribose glycohydrolase ARH3 | 1.68E-03 | -1.42 | 6.25E-04 | -1.27 | 1.26E-03 | -1.54 |
| Q8NFZ8 | CADM4 | Cell adhesion molecule 4 | 3.19E-04 | -1.42 | 7.88E-03 | -1.85 | NA | NA |
| P13987 | CD59 | CD59 glycoprotein | 9.78E-03 | -1.38 | 2.02E-02 | -2.07 | 9.94E-05 | -3.71 |
| Q5T9C9 | PIP5KL1 | Phosphatidylinositol 4-phosphate 5-kinase-like protein 1 | 8.74E-05 | -1.37 | 4.47E-03 | -1.62 | 1.58E-04 | -2.62 |
| P47756 | CAPZB | F-actin-capping protein subunit beta | 1.48E-04 | -1.36 | 1.71E-01 | -0.55 | 9.11E-04 | -2.09 |
| Q9NXG0 | CNTLN | Centlein | 5.49E-04 | -1.35 | 5.44E-03 | -2.05 | NA | NA |
| P09382 | LGALS1 | Galectin-1 | 5.53E-03 | -1.35 | 6.48E-03 | -2.34 | 7.18E-04 | -2.87 |
| P04899;P08754;P63096 | GNAI1;GNAI2;GNAI3 | Guanine nucleotide-binding protein G(i) subunit alpha-2 | 6.61E-03 | -1.33 | 3.34E-02 | -1.24 | 9.40E-05 | -2.75 |
| P40227;Q92526 | CCT6A;CCT6B | T-complex protein 1 subunit zeta | 3.89E-03 | -1.32 | 1.07E-01 | -0.70 | 6.72E-04 | -1.87 |
| Q01082 | SPTBN1 | Spectrin beta chain, non-erythrocytic 1 | 2.38E-03 | -1.30 | 4.80E-02 | -0.88 | 5.23E-04 | -1.57 |
| Q08722 | CD47 | Leukocyte surface antigen CD47 | 4.50E-03 | -1.29 | 7.32E-03 | -2.26 | 1.56E-03 | -3.20 |
| P22748 | CA4 | Carbonic anhydrase 4 | 9.30E-04 | -1.27 | 3.77E-04 | -1.84 | 6.66E-03 | -2.19 |
| Q01484;Q9Y4W6 | AFG3L2;ANK2 | Ankyrin-2 | 5.58E-03 | -1.25 | 2.27E-03 | -1.15 | 1.11E-03 | -1.70 |
| Q15126 | PMVK | Phosphomevalonate kinase | 9.07E-03 | -1.24 | 8.70E-01 | 0.07 | 5.13E-03 | -1.61 |
| Q9Y5L0 | TNPO3 | Transportin-3 | 3.51E-03 | -1.23 | 2.87E-01 | -0.47 | 4.77E-03 | -2.20 |
| Q93034 | CUL5 | Cullin-5 | 1.49E-04 | -1.18 | 1.47E-01 | -0.58 | 5.27E-05 | -1.79 |
| P11171;Q9H4G0 | EPB41;EPB41L1 | Protein 4.1 | 3.65E-03 | -1.17 | 3.39E-03 | -1.74 | 2.99E-03 | -2.39 |
| Q96DA2 | RAB39B | Ras-related protein Rab-39B | 7.16E-03 | -1.13 | 2.06E-04 | -1.66 | 5.25E-03 | -1.80 |
| P05026 | ATP1B1 | Sodium/potassium-transporting ATPase subunit beta-1 | 7.63E-03 | -1.09 | 1.50E-03 | -2.37 | 1.34E-03 | -4.41 |
| P53816 | PLAAT3 | Phospholipase A and acyltransferase 3 | 5.76E-03 | -1.05 | 2.56E-03 | -1.06 | 3.10E-03 | -1.66 |
| P20340;Q9NRW1 | RAB6A;RAB6B | Ras-related protein Rab-6A | 8.46E-03 | -1.04 | 1.57E-04 | -1.56 | 1.40E-04 | -2.44 |
| P08134;P61586;P62745 | RHOA;RHOB;RHOC | Rho-related GTP-binding protein RhoC | 7.27E-03 | -1.00 | 1.93E-03 | -1.75 | 6.82E-01 | -0.20 |
| Q99829 | CPNE1 | Copine-1 | 4.34E-03 | -0.93 | 5.69E-03 | -1.66 | NA | NA |
| P13693 | TPT1 | Translationally-controlled tumor protein | 5.80E-03 | -0.89 | 7.50E-02 | -0.97 | 4.66E-03 | -2.06 |
| Q9H977 | WDR54 | WD repeat-containing protein 54 | 4.75E-03 | 1.70 | 3.17E-02 | 0.66 | 7.60E-02 | -0.92 |

| | | | | | | | | |
|---|---|---|---|---|---|---|---|---|
| P07315 | CRYGC | Gamma-crystallin C | 1.78E-03 | 1.79 | 8.80E-02 | 0.57 | 9.50E-02 | -0.67 |
| Q9BXT8 | RNF17 | RING finger protein 17 | 8.55E-04 | 1.90 | 2.30E-01 | 0.50 | 3.91E-01 | -0.36 |
| P07205 | PGK2 | Phosphoglycerate kinase 2 | 5.99E-03 | 1.91 | 3.29E-01 | 0.41 | 3.28E-01 | -0.57 |
| Q64LD2 | WDR25 | WD repeat-containing protein 25 | 9.02E-04 | 2.00 | 8.35E-03 | 1.28 | 3.82E-01 | 0.45 |
| P32754 | HPD | 4-hydroxyphenylpyruvate dioxygenase | 6.61E-04 | 2.10 | 4.16E-02 | 1.16 | 1.52E-01 | 0.91 |
| P07900;P08238;Q58FF6; Q58FF8 | HSP90AA1;HSP90AB1; HSP90AB2P;HSP90AB4 P | Heat shock protein HSP 90-alpha | NA | NA | 2.05E-04 | -2.86 | NA | NA |
| P80723 | BASP1 | Brain acid soluble protein 1 | 5.77E-01 | -0.39 | 4.76E-03 | -3.21 | 2.82E-04 | -4.77 |
| Q16270 | IGFBP7 | Insulin-like growth factor-binding protein 7 | 1.04E-01 | -0.72 | 3.96E-03 | -3.11 | NA | NA |
| O94856 | NFASC | Neurofascin | 2.34E-02 | -1.13 | 4.48E-04 | -2.84 | 2.31E-04 | -3.05 |
| O43491;Q9Y2J2 | EPB41L2;EPB41L3 | Band 4.1-like protein 2 | 6.82E-02 | -0.65 | 2.06E-03 | -2.67 | NA | NA |
| Q8N3J6 | CADM2 | Cell adhesion molecule 2 | 4.24E-02 | -1.16 | 4.80E-03 | -2.63 | 2.91E-04 | -2.86 |
| Q6N022;Q9P273 | TENM3;TENM4 | Teneurin-4 | 6.08E-01 | -0.23 | 3.67E-03 | -2.53 | NA | NA |
| Q8IXI1 | RHOT2 | Mitochondrial Rho GTPase 2 | 6.91E-02 | -0.80 | 5.03E-03 | -2.48 | NA | NA |
| O14960 | LECT2 | Leukocyte cell-derived chemotaxin-2 | 9.27E-01 | 0.08 | 2.01E-03 | -2.47 | NA | NA |
| Q9BY67 | CADM1 | Cell adhesion molecule 1 | 1.12E-02 | -1.73 | 2.71E-03 | -2.47 | NA | NA |
| P11233 | RALA | Ras-related protein Ral-A | 1.28E-01 | -0.61 | 2.33E-04 | -2.39 | 1.92E-05 | -3.41 |
| P06899;P23527;P33778; Q16778;Q8N257;Q99880 | H2BC11;H2BC13;H2BC 17;H2BC21;H2BC3;H2B U1 | Histone H2B type 1-J | NA | NA | 7.18E-03 | -2.39 | NA | NA |
| P39060 | COL18A1 | Collagen alpha-1(XVIII) chain | 2.67E-02 | -2.13 | 3.41E-03 | -2.38 | NA | NA |
| P63165 | SUMO1 | Small ubiquitin-related modifier 1 | 1.97E-02 | -1.30 | 8.96E-03 | -2.36 | NA | NA |
| O60884 | DNAJA2 | DnaJ homolog subfamily A member 2 | 4.49E-02 | -0.97 | 4.70E-04 | -2.35 | NA | NA |
| P35241 | RDX | Radixin | 3.66E-02 | -1.24 | 3.84E-03 | -2.34 | 5.16E-03 | -2.52 |
| Q9ULX7 | CA14 | Carbonic anhydrase 14 | 1.34E-01 | -0.73 | 4.44E-04 | -2.33 | 8.97E-04 | -3.28 |
| Q9BX67 | JAM3 | Junctional adhesion molecule C | 2.47E-02 | -1.24 | 2.43E-03 | -2.23 | 1.13E-03 | -2.63 |
| Q9NTI2 | ATP8A2 | Phospholipid-transporting ATPase IB | 9.54E-02 | -0.86 | 1.46E-03 | -2.20 | 2.83E-03 | -1.84 |
| Q9UGT4 | SUSD2 | Sushi domain-containing protein 2 | 1.04E-02 | -1.94 | 2.58E-03 | -2.20 | NA | NA |
| P08754 | GNAI3 | Guanine nucleotide-binding protein G(i) subunit alpha-3 | 1.31E-02 | -1.45 | 3.88E-03 | -2.18 | NA | NA |
| O95715 | CXCL14 | C-X-C motif chemokine 14 | 3.78E-02 | -1.87 | 5.97E-03 | -2.17 | NA | NA |
| Q9UHN6 | CEMIP2 | Cell surface hyaluronidase | 1.55E-02 | -1.45 | 4.14E-03 | -2.12 | 1.14E-03 | -2.81 |

| | | | | | | | | |
|---|---|---|---|---|---|---|---|---|
| Q6N022 | TENM4 | Teneurin-4 | 1.51E-01 | -0.63 | 7.95E-03 | -2.11 | NA | NA |
| P20340 | RAB6A | Ras-related protein Rab-6A | 4.33E-02 | -0.57 | 2.77E-04 | -2.11 | 1.47E-03 | -2.11 |
| Q8IVW8 | SPNS2 | Protein spinster homolog 2 | 8.65E-02 | -0.86 | 1.35E-03 | -2.10 | NA | NA |
| Q9UI40 | SLC24A2 | Sodium/potassium/calcium exchanger 2 | 3.55E-02 | -0.82 | 1.09E-03 | -2.07 | 2.02E-04 | -2.96 |
| P08134;P61586 | RHOA;RHOC | Rho-related GTP-binding protein RhoC | 8.45E-01 | -0.14 | 1.37E-03 | -2.06 | 3.95E-04 | -2.23 |
| P34096 | RNASE4 | Ribonuclease 4 | 5.00E-01 | -0.36 | 8.18E-03 | -2.06 | NA | NA |
| P04156 | PRNP | Major prion protein | 3.81E-02 | -0.79 | 5.19E-03 | -2.01 | 1.43E-03 | -2.49 |
| Q9NQ79 | CRTAC1 | Cartilage acidic protein 1 | 6.29E-02 | -0.85 | 4.60E-03 | -1.97 | 6.09E-05 | -3.24 |
| Q6UWM7 | LCTL | Lactase-like protein | 2.07E-02 | -1.28 | 5.96E-04 | -1.95 | 1.66E-03 | -1.88 |
| P48509 | CD151 | CD151 antigen | 7.48E-02 | -1.06 | 6.43E-03 | -1.95 | 4.19E-03 | -2.45 |
| O94812 | BAIAP3 | BAI1-associated protein 3 | NA | NA | 2.74E-03 | -1.94 | NA | NA |
| P26038 | MSN | Moesin | 4.98E-02 | -0.99 | 4.77E-04 | -1.93 | 7.93E-04 | -1.61 |
| Q92823 | NRCAM | Neuronal cell adhesion molecule | 1.39E-02 | -1.31 | 5.65E-03 | -1.93 | 8.66E-05 | -3.02 |
| O75955 | FLOT1 | Flotillin-1 | 3.90E-02 | -0.79 | 3.84E-04 | -1.92 | 2.16E-04 | -2.58 |
| P29317 | EPHA2 | Ephrin type-A receptor 2 | 1.80E-02 | -1.08 | 1.65E-03 | -1.89 | NA | NA |
| P0DOY2;P0DOY3 | IGLC2;IGLC3 | Immunoglobulin lambda constant 2 | 8.44E-01 | 0.09 | 4.00E-03 | -1.88 | 1.42E-03 | -2.35 |
| P01034 | CST3 | Cystatin-C | 8.97E-01 | -0.10 | 6.13E-03 | -1.87 | 4.56E-03 | -2.73 |
| P36955 | SERPINF1 | Pigment epithelium-derived factor | 6.88E-01 | 0.24 | 5.31E-03 | -1.84 | NA | NA |
| O95490 | ADGRL2 | Adhesion G protein-coupled receptor L2 | 1.10E-01 | -0.87 | 6.96E-03 | -1.83 | 6.09E-04 | -2.51 |
| P15153;P17081;P60953;<br>P63000;P84095 | CDC42;RAC1;RAC2;RHO<br>G;RHOQ | Ras-related C3 botulinum toxin substrate 2 | 3.02E-01 | 0.69 | 4.40E-03 | -1.82 | NA | NA |
| Q9Y624 | F11R | Junctional adhesion molecule A | 1.74E-02 | -0.98 | 7.06E-03 | -1.75 | NA | NA |
| Q96ST8 | CEP89 | Centrosomal protein of 89 kDa | 1.79E-01 | -0.76 | 6.85E-03 | -1.75 | 9.31E-04 | -3.54 |
| P61586 | RHOA | Transforming protein RhoA | 7.12E-01 | -0.15 | 3.61E-04 | -1.71 | 1.05E-02 | -2.17 |
| P04216 | THY1 | Thy-1 membrane glycoprotein | 4.05E-01 | -0.46 | 3.65E-03 | -1.70 | 1.32E-03 | -3.51 |
| P11233;P11234 | RALA;RALB | Ras-related protein Ral-A | 2.16E-01 | -0.52 | 2.87E-04 | -1.70 | 6.82E-03 | -1.40 |
| Q15223 | NECTIN1 | Nectin-1 | 4.61E-01 | -0.36 | 8.62E-03 | -1.68 | NA | NA |
| P31946;P61981 | YWHAB;YWHAG | 14-3-3 protein beta/alpha | 1.33E-02 | -1.98 | 8.14E-04 | -1.68 | NA | NA |
| P31946;P61981.1 | YWHAB;YWHAG | 14-3-3 protein beta/alpha | NA | NA | 8.14E-04 | -1.68 | NA | NA |
| P10909 | CLU | Clusterin | 8.82E-02 | -1.05 | 6.87E-03 | -1.65 | 3.13E-03 | -1.76 |
| Q13515 | BFSP2 | Phakinin | 1.05E-01 | -0.62 | 6.58E-04 | -1.64 | 1.13E-03 | -1.99 |

| | | | | | | | | |
|---|---|---|---|---|---|---|---|---|
| Q96KP4 | CNDP2 | Cytosolic non-specific dipeptidase | 8.94E-02 | -0.57 | 9.59E-04 | -1.64 | 3.65E-04 | -2.20 |
| P11171 | EPB41 | Protein 4.1 | 2.68E-02 | -1.36 | 6.52E-03 | -1.61 | 4.98E-05 | -2.18 |
| P12268;P20839 | IMPDH1;IMPDH2 | Inosine-5'-monophosphate dehydrogenase 2 | 9.31E-02 | -0.73 | 2.48E-03 | -1.60 | NA | NA |
| P01857;P01860 | IGHG1;IGHG3 | Immunoglobulin heavy constant gamma 1 | 9.31E-01 | -0.06 | 4.91E-03 | -1.57 | 2.48E-03 | -2.00 |
| P62714;P67775 | PPP2CA;PPP2CB | Serine/threonine-protein phosphatase 2A catalytic subunit beta isoform | NA | NA | 8.53E-03 | -1.53 | NA | NA |
| Q1MSJ5 | CSPP1 | Centrosome and spindle pole-associated protein 1 | 3.86E-01 | 0.44 | 1.10E-03 | -1.53 | 1.20E-03 | -2.33 |
| O75436;Q4G0F5 | VPS26A;VPS26B | Vacuolar protein sorting-associated protein 26A | 1.23E-01 | 0.83 | 2.18E-03 | 1.98 | 9.44E-02 | 0.91 |
| Q13555;Q13557;Q9UQM7.1 | CAMK2A;CAMK2D;CAMK2G | Calcium/calmodulin-dependent protein kinase type II subunit gamma | NA | NA | NA | NA | 7.79E-04 | -3.63 |
| O43491;Q9H4G0;Q9Y2J2 | EPB41L1;EPB41L2;EPB41L3 | Band 4.1-like protein 2 | 1.81E-02 | -1.46 | 1.80E-02 | -1.43 | 8.12E-05 | -3.32 |
| Q14254 | FLOT2 | Flotillin-2 | 1.51E-02 | -1.06 | 1.06E-02 | -2.10 | 1.39E-04 | -3.03 |
| P04899;P08754;P63096.1 | GNAI1;GNAI2;GNAI3 | Guanine nucleotide-binding protein G(i) subunit alpha-2 | NA | NA | NA | NA | 9.40E-05 | -2.75 |
| P60953 | CDC42 | Cell division control protein 42 homolog | 2.60E-01 | -0.44 | 2.43E-02 | -1.14 | 2.70E-05 | -2.70 |
| P06733;P09104;P13929 | ENO1;ENO2;ENO3 | Alpha-enolase | 8.27E-02 | -0.93 | 1.14E-01 | -1.10 | 9.51E-03 | -2.68 |
| Q9H0C3 | TMEM117 | Transmembrane protein 117 | 4.14E-01 | -0.33 | 4.05E-02 | -0.91 | 3.28E-04 | -2.61 |
| Q9BQJ4 | TMEM47 | Transmembrane protein 47 | 7.41E-01 | -0.19 | 9.29E-02 | -0.95 | 1.77E-03 | -2.60 |
| O94760 | DDAH1 | N(G),N(G)-dimethylarginine dimethylaminohydrolase 1 | 3.43E-01 | -0.47 | 3.46E-02 | -1.48 | 4.43E-04 | -2.59 |
| P0C7U9 | FAM87A | Protein FAM87A | NA | NA | NA | NA | 3.57E-05 | -2.58 |
| Q03135 | CAV1 | Caveolin-1 | 7.86E-01 | -0.16 | 4.52E-02 | -1.22 | 1.28E-03 | -2.57 |
| P36969 | GPX4 | Phospholipid hydroperoxide glutathione peroxidase | 3.11E-01 | 0.40 | 2.08E-02 | -1.29 | 5.88E-04 | -2.56 |
| Q14244 | MAP7 | Ensconsin | 7.36E-01 | -0.19 | 2.46E-02 | -1.30 | 1.09E-04 | -2.53 |
| Q92743 | HTRA1 | Serine protease HTRA1 | 9.23E-01 | -0.06 | 4.44E-02 | -1.25 | 7.65E-03 | -2.46 |
| Q92542 | NCSTN | Nicastrin | 1.90E-01 | -0.42 | 4.25E-02 | -1.25 | 1.94E-04 | -2.43 |
| P09972 | ALDOC | Fructose-bisphosphate aldolase C | 1.11E-01 | -0.53 | 6.33E-03 | -1.18 | 1.27E-04 | -2.42 |
| P08237;P17858 | PFKL;PFKM | ATP-dependent 6-phosphofructokinase, muscle type | 7.58E-01 | 0.12 | 2.67E-01 | -0.38 | 9.36E-05 | -2.39 |
| P27361;P28482 | MAPK1;MAPK3 | Mitogen-activated protein kinase 3 | 3.31E-02 | -0.60 | 2.13E-02 | -1.04 | 3.39E-04 | -2.37 |
| P04792 | HSPB1 | Heat shock protein beta-1 | 4.04E-02 | -1.20 | 1.16E-02 | -1.43 | 7.74E-04 | -2.36 |
| Q9BWS9 | CHID1 | Chitinase domain-containing protein 1 | 3.28E-01 | -0.40 | 6.98E-02 | -1.30 | 3.52E-04 | -2.33 |
| P63000 | RAC1 | Ras-related C3 botulinum toxin substrate 1 | 8.31E-01 | 0.10 | 1.50E-02 | -1.66 | 8.96E-05 | -2.32 |
| P15151 | PVR | Poliovirus receptor | 1.61E-02 | -1.43 | 1.55E-02 | -1.34 | 4.02E-04 | -2.29 |

| | | | | | | | | |
|---|---|---|---|---|---|---|---|---|
| P61626 | LYZ | Lysozyme C | 8.51E-01 | -0.11 | 1.22E-02 | -1.87 | 7.01E-03 | -2.23 |
| Q08357 | SLC20A2 | Sodium-dependent phosphate transporter 2 | 5.37E-02 | -1.23 | 1.68E-01 | -0.81 | 2.84E-04 | -2.21 |
| P21860 | ERBB3 | Receptor tyrosine-protein kinase erbB-3 | 6.28E-01 | -0.27 | 3.86E-02 | -1.54 | 9.71E-03 | -2.19 |
| P09960 | LTA4H | Leukotriene A-4 hydrolase | 9.58E-01 | -0.02 | 2.72E-02 | -0.95 | 1.71E-04 | -2.17 |
| Q15286 | RAB35 | Ras-related protein Rab-35 | 1.04E-02 | -1.30 | 2.18E-02 | -1.63 | 2.84E-03 | -2.15 |
| P10301;P62070 | RRAS;RRAS2 | Ras-related protein R-Ras | 2.78E-02 | -1.61 | 8.21E-02 | -0.77 | 1.11E-03 | -2.11 |
| P10301;P62070.1 | RRAS;RRAS2 | Ras-related protein R-Ras | NA | NA | 8.21E-02 | -0.77 | 1.11E-03 | -2.11 |
| P22059 | OSBP | Oxysterol-binding protein 1 | 3.51E-02 | -0.66 | 1.85E-01 | -0.56 | 5.61E-03 | -2.01 |
| P06733 | ENO1 | Alpha-enolase | 5.00E-01 | -0.31 | 5.48E-02 | -0.99 | 6.04E-04 | -1.99 |
| P09488;P28161 | GSTM1;GSTM2 | Glutathione S-transferase Mu 1 | NA | NA | NA | NA | 4.77E-03 | -1.99 |
| Q9Y6D5 | ARFGEF2 | Brefeldin A-inhibited guanine nucleotide-exchange protein 2 | 4.67E-01 | 0.20 | 8.70E-03 | -1.38 | 2.40E-05 | -1.95 |
| P09211 | GSTP1 | Glutathione S-transferase P | 4.46E-02 | -1.14 | 6.72E-03 | -1.05 | 2.34E-03 | -1.95 |
| Q99832 | CCT7 | T-complex protein 1 subunit eta | 4.03E-02 | -0.92 | 8.16E-03 | -0.94 | 5.74E-05 | -1.94 |
| P17081;P60953;P63000;P84095 | CDC42;RAC1;RHOG;RHOQ | Rho-related GTP-binding protein RhoQ | NA | NA | NA | NA | 8.96E-04 | -1.93 |
| P55344 | LIM2 | Lens fiber membrane intrinsic protein | 7.91E-01 | 0.19 | 1.85E-01 | -0.69 | 2.95E-03 | -1.93 |
| P52888 | THOP1 | Thimet oligopeptidase | NA | NA | NA | NA | 7.18E-04 | -1.92 |
| P55064 | AQP5 | Aquaporin-5 | 3.64E-01 | -0.53 | 2.44E-01 | -0.77 | 9.48E-03 | -1.91 |
| Q96F07 | CYFIP2 | Cytoplasmic FMR1-interacting protein 2 | 2.97E-02 | -1.21 | 9.41E-03 | -1.13 | 8.82E-04 | -1.91 |
| O75663 | TIPRL | TIP41-like protein | 2.17E-01 | -0.45 | 2.86E-03 | -1.10 | 3.99E-04 | -1.91 |
| P23528 | CFL1 | Cofilin-1 | 8.90E-02 | -0.61 | 2.32E-02 | -1.04 | 7.07E-03 | -1.91 |
| P60900 | PSMA6 | Proteasome subunit alpha type-6 | 3.49E-01 | -0.38 | 2.12E-01 | -0.24 | 1.89E-04 | -1.89 |
| P13639 | EEF2 | Elongation factor 2 | 6.89E-02 | -0.57 | 9.34E-03 | -0.82 | 8.53E-04 | -1.87 |
| P38919 | EIF4A3 | Eukaryotic initiation factor 4A-III | 5.64E-02 | -0.77 | 4.85E-01 | -0.31 | 1.43E-03 | -1.85 |
| P56192 | MARS1 | Methionine--tRNA ligase, cytoplasmic | 2.10E-02 | -1.57 | 8.18E-01 | -0.11 | 8.03E-03 | -1.85 |
| P26639 | TARS1 | Threonine--tRNA ligase 1, cytoplasmic | 2.56E-01 | -0.48 | 6.43E-02 | -0.74 | 6.39E-04 | -1.85 |
| O75083 | WDR1 | WD repeat-containing protein 1 | 8.79E-01 | -0.05 | 5.74E-01 | -0.24 | 3.77E-03 | -1.83 |
| Q9Y281 | CFL2 | Cofilin-2 | 3.80E-02 | -0.93 | 4.40E-01 | -0.28 | 1.37E-03 | -1.83 |
| Q9UJW0 | DCTN4 | Dynactin subunit 4 | 3.23E-02 | -1.21 | 3.64E-01 | -0.24 | 2.77E-04 | -1.83 |
| P04406 | GAPDH | Glyceraldehyde-3-phosphate dehydrogenase | 1.38E-01 | 0.52 | 1.77E-01 | -0.72 | 2.67E-04 | -1.78 |
| Q9BTW9 | TBCD | Tubulin-specific chaperone D | 1.48E-02 | -1.10 | 1.13E-01 | -0.71 | 3.44E-03 | -1.78 |

| | | | | | | | | |
|---|---|---|---|---|---|---|---|---|
| P28482 | MAPK1 | Mitogen-activated protein kinase 1 | 4.04E-02 | -0.96 | 8.26E-02 | -0.79 | 1.23E-03 | -1.77 |
| Q12934 | BFSP1 | Filensin | 1.09E-01 | -0.54 | 4.91E-02 | -0.70 | 1.20E-03 | -1.77 |
| O75935 | DCTN3 | Dynactin subunit 3 | 9.61E-01 | -0.03 | 1.98E-01 | 0.59 | 1.53E-03 | -1.76 |
| Q13619 | CUL4A | Cullin-4A | 2.29E-02 | -0.85 | 1.19E-01 | -0.62 | 1.52E-03 | -1.76 |
| P62258 | YWHAE | 14-3-3 protein epsilon | 9.98E-02 | -0.94 | 2.54E-01 | -0.37 | 3.29E-03 | -1.76 |
| Q9NY33 | DPP3 | Dipeptidyl peptidase 3 | 1.70E-01 | 0.63 | 2.15E-01 | -0.54 | 8.91E-04 | -1.75 |
| Q9UJ70 | NAGK | N-acetyl-D-glucosamine kinase | 1.43E-01 | 0.41 | 1.76E-01 | -0.42 | 4.24E-04 | -1.74 |
| P50747 | HLCS | Biotin--protein ligase | 1.22E-01 | 0.65 | 6.06E-01 | -0.19 | 6.66E-04 | -1.74 |
| P61201 | COPS2 | COP9 signalosome complex subunit 2 | 2.61E-02 | -0.78 | 4.28E-01 | -0.35 | 9.44E-04 | -1.74 |
| P48449 | LSS | Lanosterol synthase | 3.38E-01 | -0.41 | 5.10E-02 | -0.83 | 3.07E-04 | -1.72 |
| P0C870 | JMJD7 | Bifunctional peptidase and (3S)-lysyl hydroxylase JMJD7 | 6.31E-02 | 0.79 | 2.48E-01 | -0.45 | 2.72E-03 | -1.72 |
| P48739 | PITPNB | Phosphatidylinositol transfer protein beta isoform | 1.12E-01 | -0.81 | 3.99E-01 | -0.35 | 9.19E-03 | -1.72 |
| B3SHH9 | TMEM114 | Transmembrane protein 114 | 1.38E-01 | -0.88 | 2.61E-02 | -1.40 | 1.14E-04 | -1.71 |
| A2RU48 | SMCO3 | Single-pass membrane and coiled-coil domain-containing protein 3 | 9.35E-01 | 0.03 | 6.98E-04 | -1.30 | 3.22E-04 | -1.70 |
| Q7L5N1 | COPS6 | COP9 signalosome complex subunit 6 | 3.59E-01 | -0.37 | 5.06E-01 | -0.22 | 1.29E-03 | -1.68 |
| P61163 | ACTR1A | Alpha-centractin | 6.75E-01 | 0.13 | 1.11E-02 | -1.12 | 9.66E-03 | -1.67 |
| O95352 | ATG7 | Ubiquitin-like modifier-activating enzyme ATG7 | 1.85E-01 | -0.43 | 4.73E-01 | -0.31 | 6.01E-03 | -1.66 |
| P17252 | PRKCA | Protein kinase C alpha type | NA | NA | NA | NA | 4.16E-03 | -1.66 |
| E9PAV3;Q13765 | NACA | Nascent polypeptide-associated complex subunit alpha, muscle-specific form | 1.20E-02 | -0.84 | 3.20E-03 | -1.45 | 6.82E-03 | -1.65 |
| O95834 | EML2 | Echinoderm microtubule-associated protein-like 2 | 2.88E-01 | 0.40 | 2.21E-01 | -0.41 | 2.17E-03 | -1.65 |
| P08238 | HSP90AB1 | Heat shock protein HSP 90-beta | 3.94E-02 | -1.24 | 1.68E-03 | -1.05 | 3.88E-03 | -1.65 |
| P28070 | PSMB4 | Proteasome subunit beta type-4 | 6.96E-01 | 0.14 | 1.39E-01 | -0.46 | 1.61E-04 | -1.64 |
| O95861 | BPNT1 | 3'(2'),5'-bisphosphate nucleotidase 1 | 8.37E-01 | -0.08 | 6.62E-01 | -0.16 | 6.45E-04 | -1.63 |
| P60510 | PPP4C | Serine/threonine-protein phosphatase 4 catalytic subunit | NA | NA | 7.21E-01 | 0.17 | 3.61E-03 | -1.63 |
| Q9P2R3 | ANKFY1 | Rabankyrin-5 | NA | NA | 6.66E-02 | -0.73 | 1.57E-03 | -1.61 |
| P49327 | FASN | Fatty acid synthase | 4.20E-02 | -0.70 | 5.75E-02 | -0.71 | 4.40E-04 | -1.61 |
| Q9UBQ5 | EIF3K | Eukaryotic translation initiation factor 3 subunit K | NA | NA | NA | NA | 3.47E-03 | -1.61 |
| P53602 | MVD | Diphosphomevalonate decarboxylase | 5.90E-01 | 0.24 | 1.57E-01 | -0.63 | 8.25E-03 | -1.61 |
| Q70IA6 | MOB2 | MOB kinase activator 2 | 8.18E-02 | -0.61 | 1.68E-02 | -0.68 | 2.82E-03 | -1.61 |

| | | | | | | | | |
|---|---|---|---|---|---|---|---|---|
| P29218 | IMPA1 | Inositol monophosphatase 1 | 8.21E-01 | 0.07 | 2.41E-01 | -0.48 | 1.47E-03 | -1.60 |
| O43813 | LANCL1 | Glutathione S-transferase LANCL1 | 1.93E-01 | 0.52 | 4.25E-02 | -0.61 | 7.03E-04 | -1.60 |
| P13010 | XRCC5 | X-ray repair cross-complementing protein 5 | 7.71E-02 | -0.93 | 1.28E-01 | -0.56 | 1.06E-03 | -1.60 |
| Q16401 | PSMD5 | 26S proteasome non-ATPase regulatory subunit 5 | 2.12E-01 | -0.68 | 3.46E-01 | -0.46 | 1.49E-03 | -1.60 |
| Q8IW45 | NAXD | ATP-dependent (S)-NAD(P)H-hydrate dehydratase | 5.67E-01 | 0.22 | 9.62E-02 | -0.55 | 4.07E-03 | -1.58 |
| O95163 | ELP1 | Elongator complex protein 1 | 6.53E-02 | -0.76 | 3.90E-01 | -0.36 | 2.02E-04 | -1.58 |
| P30566 | ADSL | Adenylosuccinate lyase | 2.56E-01 | 0.36 | 3.25E-01 | -0.35 | 9.58E-03 | -1.57 |
| Q9Y3F4 | STRAP | Serine-threonine kinase receptor-associated protein | 1.44E-01 | -0.57 | 3.40E-02 | -0.92 | 5.34E-03 | -1.57 |
| Q7L523 | RRAGA | Ras-related GTP-binding protein A | 3.25E-01 | -0.36 | 7.44E-01 | -0.13 | 2.26E-03 | -1.57 |
| P23528;Q9Y281 | CFL1;CFL2 | Cofilin-1 | 7.76E-01 | 0.11 | 3.16E-01 | -0.34 | 1.78E-03 | -1.57 |
| P54920 | NAPA | Alpha-soluble NSF attachment protein | 2.03E-01 | -0.81 | 7.09E-02 | -0.76 | 3.38E-04 | -1.56 |
| P08758 | ANXA5 | Annexin A5 | 2.30E-02 | -1.28 | 1.49E-01 | -0.55 | 1.22E-03 | -1.56 |
| P14324 | FDPS | Farnesyl pyrophosphate synthase | 5.93E-01 | -0.17 | 7.68E-04 | -1.14 | 2.67E-04 | -1.55 |
| Q9H0R4 | HDHD2 | Haloacid dehalogenase-like hydrolase domain-containing protein 2 | 4.50E-01 | -0.24 | 8.05E-03 | -0.67 | 1.60E-03 | -1.55 |
| P21980 | TGM2 | Protein-glutamine gamma-glutamyltransferase 2 | 3.41E-02 | -1.19 | 1.78E-01 | -0.51 | 3.29E-03 | -1.55 |
| Q06830 | PRDX1 | Peroxiredoxin-1 | 1.25E-01 | -0.66 | 5.75E-01 | -0.18 | 2.84E-03 | -1.54 |
| P07737 | PFN1 | Profilin-1 | 4.63E-02 | -1.26 | 3.07E-02 | -1.56 | 1.25E-03 | -1.52 |
| P27361 | MAPK3 | Mitogen-activated protein kinase 3 | 5.73E-02 | -0.81 | 1.74E-01 | -0.59 | 2.55E-03 | -1.51 |
| Q9UQN3 | CHMP2B | Charged multivesicular body protein 2b | 6.96E-01 | -0.17 | 6.54E-03 | -1.07 | 5.47E-03 | -1.50 |
| P17987 | TCP1 | T-complex protein 1 subunit alpha | 5.71E-01 | -0.22 | 1.01E-01 | -0.48 | 3.78E-03 | -1.50 |
