## Supplemental Table S2-S7 for "Proteome Remodeling of the Eye Lens at 50 Years Identified with Data-Independent Acquisition"

Supplemental Tables S2-S7 – Gene Ontologies calculated as enriched by PSEA-Quant in a single region of the Young/Old Lens relative to the Old/Young Lens. Significance was determined at a p-value < 0.01 and FDR < 0.1. PSEA-Quant was operated in labeled mode with 1,000,000 iterations used to empirically evaluate p-values. P-values below 1/1,000,000 ($1x10^{-6}$) are marked as such and correspond to distributions of protein enrichment not similar to prior distributions evaluated in 1,000,000 iterations. Header is colored by corresponding region of lens and annotation enrichment refers to specific table annotation header (red = cortex, green = outer nucleus, blue = inner nucleus). Number of proteins refers to proteins detected in dataset (no missing values from 16 samples) with the corresponding annotation.

| **Supplemental Table S2**<br>**Annotation: Ontology Enriched in Young Lens Cortex relative to Old Lens Cortex** | **P-Value** | **FDR** | **Number of Proteins with Annotation in Dataset** |
|---|---|---|---|
| GO:0090150 Establishment of protein localization to membrane | <1E-06 | 1.75E-02 | 30 |
| GO:0006334 Nucleosome assembly | <1E-06 | 1.75E-02 | 11 |
| GO:0044391 Ribosomal subunit | <1E-06 | 1.75E-02 | 14 |
| GO:0031224 Intrinsic component of membrane | <1E-06 | 1.75E-02 | 206 |
| GO:0005886 Plasma membrane | <1E-06 | 1.75E-02 | 295 |
| GO:0006605 Protein targeting | <1E-06 | 1.75E-02 | 40 |
| GO:0006612 Protein targeting to membrane | <1E-06 | 1.75E-02 | 22 |
| GO:0006810 Transport | <1E-06 | 1.75E-02 | 347 |
| GO:0006886 Intracellular protein transport | <1E-06 | 1.75E-02 | 104 |
| GO:0015031 Protein transport | <1E-06 | 1.75E-02 | 178 |
| GO:0016020 Membrane | <1E-06 | 1.75E-02 | 502 |
| GO:0016818 Hydrolase activity, acting on acid anhydrides, in phosphorus-containing anhydrides | <1E-06 | 1.75E-02 | 140 |
| GO:0017111 Nucleoside-triphosphatase activity | <1E-06 | 1.75E-02 | 133 |
| GO:0031090 Organelle membrane | <1E-06 | 1.75E-02 | 254 |
| GO:0032991 Protein-containing complex | <1E-06 | 1.75E-02 | 472 |
| GO:0042470 Melanosome | <1E-06 | 1.75E-02 | 50 |
| GO:0045184 Establishment of protein localization | <1E-06 | 1.75E-02 | 182 |
| GO:0046907 Intracellular transport | <1E-06 | 1.75E-02 | 165 |
| GO:0048770 Pigment granule | <1E-06 | 1.75E-02 | 50 |
| GO:0051234 Establishment of localization | <1E-06 | 1.75E-02 | 350 |
| GO:0051649 Establishment of localization in cell | <1E-06 | 1.75E-02 | 201 |
| GO:0065004 Protein-DNA complex assembly | <1E-06 | 1.75E-02 | 11 |
| GO:0065008 Regulation of biological quality | <1E-06 | 1.75E-02 | 283 |
| GO:0071702 Organic substance transport | <1E-06 | 1.75E-02 | 230 |
| GO:0034655 Nucleobase-containing compound catabolic process | 1.00E-06 | 1.75E-02 | 93 |
| GO:0019058 Viral life cycle | 2.00E-06 | 1.75E-02 | 16 |
| GO:0016462 Pyrophosphatase activity | 2.00E-06 | 1.75E-02 | 139 |
| GO:0006415 Translational termination | 2.00E-06 | 1.75E-02 | 12 |
| GO:0071824 Protein-DNA complex subunit organization | 2.00E-06 | 1.75E-02 | 12 |
| GO:0016482 Cytosolic transport | 2.00E-06 | 1.75E-02 | 74 |
| GO:0034728 Nucleosome organization | 2.00E-06 | 1.75E-02 | 12 |
| GO:0016817 Hydrolase activity, acting on acid anhydrides | 2.00E-06 | 1.75E-02 | 141 |
| GO:0005789 Endoplasmic reticulum membrane | 3.00E-06 | 1.75E-02 | 45 |
| GO:0072599 Establishment of protein localization to endoplasmic reticulum | 3.00E-06 | 1.75E-02 | 12 |
| GO:0015075 Ion transmembrane transporter activity | 4.00E-06 | 1.75E-02 | 50 |
| GO:0000184 Nuclear-transcribed mRNA catabolic process, nonsense-mediated decay | 4.00E-06 | 1.75E-02 | 18 |
| GO:0006613 Cotranslational protein targeting to membrane | 5.00E-06 | 1.75E-02 | 12 |
| GO:0006614 SRP-dependent cotranslational protein targeting to membrane | 7.00E-06 | 1.75E-02 | 12 |

| Supplemental Table S2<br>**Annotation: Ontology Enriched in Young Lens Cortex relative to Old Lens Cortex** | P-Value | FDR | Number of Proteins with Annotation in Dataset |
|---|---|---|---|
| GO:0005575 Cellular component | 8.00E-06 | 1.75E-02 | 1201 |
| GO:0019083 Viral transcription | 9.00E-06 | 1.75E-02 | 11 |
| GO:0045047 Protein targeting to ER | 9.00E-06 | 1.75E-02 | 12 |
| GO:0031410 Cytoplasmic vesicle | 1.10E-05 | 1.75E-02 | 124 |
| GO:0016021 Integral component of membrane | 1.50E-05 | 1.75E-02 | 198 |
| GO:0031982 Vesicle | 2.30E-05 | 1.75E-02 | 154 |
| GO:0032993 Protein-DNA complex | 2.40E-05 | 1.75E-02 | 13 |
| GO:0030433 Ubiquitin-dependent ERAD pathway | 3.30E-05 | 1.75E-02 | 7 |
| GO:0044270 Cellular nitrogen compound catabolic process | 5.50E-05 | 1.75E-02 | 99 |
| GO:0043624 Cellular protein complex disassembly | 6.00E-05 | 2.36E-02 | 16 |
| GO:0072594 Establishment of protein localization to organelle | 6.00E-05 | 2.36E-02 | 43 |
| GO:0006414 Translational elongation | 6.80E-05 | 2.36E-02 | 17 |
| GO:0005515 Protein binding | 6.90E-05 | 2.36E-02 | 810 |
| GO:0000956 Nuclear-transcribed mRNA catabolic process | 7.00E-05 | 2.36E-02 | 25 |
| GO:0005743 Mitochondrial inner membrane | 7.20E-05 | 2.36E-02 | 45 |
| GO:0022890 Inorganic cation transmembrane transporter activity | 7.30E-05 | 2.36E-02 | 27 |
| GO:0019866 Organelle inner membrane | 8.00E-05 | 2.36E-02 | 46 |
| GO:0006402 mRNA catabolic process | 8.60E-05 | 2.36E-02 | 28 |
| GO:0046700 Heterocycle catabolic process | 9.60E-05 | 2.36E-02 | 100 |
| GO:0022857 Transmembrane transporter activity | 1.01E-04 | 2.36E-02 | 65 |
| GO:0015934 Large ribosomal subunit | 1.10E-04 | 2.36E-02 | 5 |
| GO:0022625 Cytosolic large ribosomal subunit | 1.12E-04 | 2.36E-02 | 5 |
| GO:0006996 Organelle organization | 1.29E-04 | 2.36E-02 | 182 |
| GO:0019439 Aromatic compound catabolic process | 1.40E-04 | 2.36E-02 | 104 |
| GO:0003735 Structural constituent of ribosome | 1.49E-04 | 2.36E-02 | 12 |
| GO:0005793 Endoplasmic reticulum-golgi intermediate compartment | 1.58E-04 | 2.36E-02 | 10 |
| GO:0006401 RNA catabolic process | 1.60E-04 | 2.36E-02 | 29 |
| GO:0071840 Cellular component organization or biogenesis | 2.24E-04 | 2.36E-02 | 376 |
| GO:0016043 Cellular component organization | 2.39E-04 | 2.36E-02 | 376 |
| GO:0030176 Integral component of endoplasmic reticulum membrane | 2.48E-04 | 2.36E-02 | 6 |
| GO:0032984 Protein-containing complex disassembly | 2.54E-04 | 2.36E-02 | 17 |
| GO:0031966 Mitochondrial membrane | 2.65E-04 | 2.36E-02 | 58 |
| GO:0016192 Vesicle-mediated transport | 2.84E-04 | 2.36E-02 | 119 |
| GO:0031227 Intrinsic component of endoplasmic reticulum membrane | 2.93E-04 | 2.36E-02 | 6 |
| GO:0009125 Nucleoside monophosphate catabolic process | 3.57E-04 | 2.36E-02 | 20 |
| GO:0016887 ATP hydrolysis activity | 3.65E-04 | 2.36E-02 | 52 |
| GO:0009169 Purine ribonucleoside monophosphate catabolic process | 3.74E-04 | 2.36E-02 | 20 |
| GO:0009158 Ribonucleoside monophosphate catabolic process | 3.93E-04 | 2.36E-02 | 20 |
| GO:0009128 Purine nucleoside monophosphate catabolic process | 4.02E-04 | 2.36E-02 | 20 |

| Supplemental Table S2<br>Annotation: Ontology Enriched in Young Lens Cortex relative to Old Lens Cortex | P-Value | FDR | Number of Proteins with Annotation in Dataset |
|---|---|---|---|
| GO:0008509 Anion transmembrane transporter activity | 4.33E-04 | 2.36E-02 | 22 |
| GO:0009203 Ribonucleoside triphosphate catabolic process | 4.39E-04 | 2.36E-02 | 54 |
| GO:0009207 Purine ribonucleoside triphosphate catabolic process | 4.52E-04 | 2.36E-02 | 54 |
| GO:0016071 mRNA metabolic process | 4.56E-04 | 2.36E-02 | 80 |
| GO:0008324 Cation transmembrane transporter activity | 5.11E-04 | 2.36E-02 | 33 |
| GO:0005215 Transporter activity | 5.78E-04 | 2.36E-02 | 106 |
| GO:0005783 Endoplasmic reticulum | 6.02E-04 | 2.36E-02 | 60 |
| GO:0009146 Purine nucleoside triphosphate catabolic process | 6.29E-04 | 2.36E-02 | 55 |
| GO:0009261 Ribonucleotide catabolic process | 6.42E-04 | 2.36E-02 | 55 |
| GO:0009154 Purine ribonucleotide catabolic process | 6.63E-04 | 2.36E-02 | 55 |
| GO:0009143 Nucleoside triphosphate catabolic process | 6.64E-04 | 2.36E-02 | 55 |
| GO:0046130 Purine ribonucleoside catabolic process | 8.06E-04 | 2.36E-02 | 55 |
| GO:0006152 Purine nucleoside catabolic process | 8.52E-04 | 2.36E-02 | 55 |
| GO:0044815 DNA packaging complex | 8.61E-04 | 2.36E-02 | 7 |
| GO:1990104 DNA bending complex | 8.72E-04 | 2.36E-02 | 7 |
| GO:0000398 mRNA splicing, via spliceosome | 9.02E-04 | 2.36E-02 | 9 |
| GO:0008380 RNA splicing | 9.18E-04 | 2.36E-02 | 16 |
| GO:0000375 RNA splicing, via transesterification reactions | 9.26E-04 | 2.36E-02 | 9 |
| GO:0000786 Nucleosome | 9.32E-04 | 2.36E-02 | 7 |
| GO:0000377 RNA splicing, via transesterification reactions with bulged adenosine as nucleophile | 9.63E-04 | 2.36E-02 | 9 |
| GO:0003774 Cytoskeletal motor activity | 9.78E-04 | 2.36E-02 | 15 |
| GO:0009166 Nucleotide catabolic process | 1.10E-03 | 2.36E-02 | 58 |
| GO:0030662 Coated vesicle membrane | 1.15E-03 | 2.36E-02 | 15 |
| GO:0044265 Cellular macromolecule catabolic process | 1.16E-03 | 2.36E-02 | 103 |
| GO:0006195 Purine nucleotide catabolic process | 1.25E-03 | 2.36E-02 | 56 |
| GO:0022411 Cellular component disassembly | 1.27E-03 | 2.36E-02 | 39 |
| GO:0072523 Purine-containing compound catabolic process | 1.29E-03 | 2.36E-02 | 56 |
| GO:0050839 Cell adhesion molecule binding | 1.31E-03 | 2.36E-02 | 15 |
| GO:0014704 Intercalated disc | 1.33E-03 | 2.36E-02 | 11 |
| GO:1901361 Organic cyclic compound catabolic process | 1.39E-03 | 2.36E-02 | 111 |
| GO:0043933 Protein-containing complex subunit organization | 1.50E-03 | 3.08E-02 | 161 |
| GO:0003924 GTPase activity | 1.59E-03 | 3.08E-02 | 71 |
| GO:0097480 Establishment of synaptic vesicle localization | 1.63E-03 | 3.55E-02 | 7 |
| GO:0044291 Cell-cell contact zone | 1.72E-03 | 3.55E-02 | 12 |
| GO:0048489 Synaptic vesicle transport | 1.73E-03 | 3.55E-02 | 7 |
| GO:0006986 Response to unfolded protein | 1.76E-03 | 3.55E-02 | 20 |
| GO:0009295 Nucleoid | 1.85E-03 | 3.55E-02 | 4 |
| GO:0042645 Mitochondrial nucleoid | 1.92E-03 | 3.55E-02 | 4 |
| GO:1901292 Nucleoside phosphate catabolic process | 1.94E-03 | 3.55E-02 | 59 |

| Supplemental Table S2<br>Annotation: Ontology Enriched in Young Lens Cortex relative to Old Lens Cortex | P-Value | FDR | Number of Proteins with Annotation in Dataset |
|---|---|---|---|
| GO:0009164 Nucleoside catabolic process | 1.97E-03 | 3.55E-02 | 56 |
| GO:1901658 Glycosyl compound catabolic process | 2.01E-03 | 3.55E-02 | 56 |
| GO:0006397 mRNA processing | 2.02E-03 | 3.55E-02 | 20 |
| GO:0030117 Membrane coat | 2.07E-03 | 3.95E-02 | 11 |
| GO:0042454 Ribonucleoside catabolic process | 2.08E-03 | 3.95E-02 | 56 |
| GO:0006836 Neurotransmitter transport | 2.23E-03 | 3.95E-02 | 7 |
| GO:0015078 Proton transmembrane transporter activity | 2.25E-03 | 3.95E-02 | 15 |
| GO:0032469 Endoplasmic reticulum calcium ion homeostasis | 2.59E-03 | 3.95E-02 | 3 |
| GO:0045296 Cadherin binding | 2.65E-03 | 3.95E-02 | 8 |
| GO:0033116 Endoplasmic reticulum-golgi intermediate compartment membrane | 2.67E-03 | 3.95E-02 | 4 |
| GO:0006839 Mitochondrial transport | 2.67E-03 | 3.95E-02 | 19 |
| GO:0009205 Purine ribonucleoside triphosphate metabolic process | 2.73E-03 | 3.95E-02 | 61 |
| GO:0006901 Vesicle coating | 2.74E-03 | 3.95E-02 | 12 |
| GO:0006413 Translational initiation | 2.85E-03 | 4.58E-02 | 31 |
| GO:0022627 Cytosolic small ribosomal subunit | 3.10E-03 | 5.00E-02 | 7 |
| GO:0009199 Ribonucleoside triphosphate metabolic process | 3.10E-03 | 5.00E-02 | 62 |
| GO:0016323 Basolateral plasma membrane | 3.34E-03 | 5.00E-02 | 34 |
| GO:0046943 Carboxylic acid transmembrane transporter activity | 3.35E-03 | 5.00E-02 | 9 |
| GO:0005342 Organic acid transmembrane transporter activity | 3.42E-03 | 5.00E-02 | 9 |
| GO:0042383 Sarcolemma | 3.46E-03 | 5.00E-02 | 11 |
| GO:0051235 Maintenance of location | 3.58E-03 | 5.00E-02 | 16 |
| GO:0045185 Maintenance of protein location | 3.71E-03 | 6.10E-02 | 12 |
| GO:0050817 Coagulation | 4.04E-03 | 6.10E-02 | 69 |
| GO:0046034 ATP metabolic process | 4.11E-03 | 6.10E-02 | 24 |
| GO:0007596 Blood coagulation | 4.14E-03 | 6.10E-02 | 69 |
| GO:0046434 Organophosphate catabolic process | 4.60E-03 | 8.38E-02 | 64 |
| GO:0050878 Regulation of body fluid levels | 4.68E-03 | 8.38E-02 | 78 |
| GO:0005840 Ribosome | 4.72E-03 | 8.38E-02 | 8 |
| GO:0032507 Maintenance of protein location in cell | 4.81E-03 | 8.88E-02 | 10 |
| GO:0065007 Biological regulation | 4.82E-03 | 8.88E-02 | 709 |
| GO:0048205 Cop I coating of golgi vesicle | 4.97E-03 | 8.94E-02 | 6 |
| GO:0048200 Golgi transport vesicle coating | 5.01E-03 | 8.94E-02 | 6 |
| GO:0032940 Secretion by cell | 5.14E-03 | 8.94E-02 | 43 |
| GO:0043229 Intracellular organelle | 5.16E-03 | 8.94E-02 | 720 |
| GO:0023061 Signal release | 5.18E-03 | 8.94E-02 | 10 |
| GO:0001505 Regulation of neurotransmitter levels | 5.31E-03 | 8.94E-02 | 8 |
| GO:0007599 Hemostasis | 5.32E-03 | 8.94E-02 | 71 |
| GO:0051650 Establishment of vesicle localization | 5.40E-03 | 8.94E-02 | 9 |
| GO:0034622 Cellular protein-containing complex assembly | 5.45E-03 | 8.94E-02 | 64 |

| **Supplemental Table S2**<br>**Annotation: Ontology Enriched in Young Lens Cortex relative to Old Lens Cortex** | **P-Value** | **FDR** | **Number of Proteins with Annotation in Dataset** |
|---|---|---|---|
| GO:0019001 Guanyl nucleotide binding | 5.67E-03 | 9.50E-02 | 96 |
| GO:0005525 GTP binding | 5.68E-03 | 9.50E-02 | 96 |
| GO:0034976 Response to endoplasmic reticulum stress | 5.72E-03 | 9.50E-02 | 15 |
| GO:0032561 Guanyl ribonucleotide binding | 5.76E-03 | 9.50E-02 | 96 |
| GO:0005253 Anion channel activity | 5.79E-03 | 9.50E-02 | 5 |
| GO:0008022 Protein c-terminus binding | 5.92E-03 | 9.50E-02 | 18 |
| GO:0046873 Metal ion transmembrane transporter activity | 6.19E-03 | 9.50E-02 | 12 |
| GO:0030120 Vesicle coat | 6.21E-03 | 9.50E-02 | 10 |
| GO:0005200 Structural constituent of cytoskeleton | 6.21E-03 | 9.50E-02 | 27 |
| GO:0030968 Endoplasmic reticulum unfolded protein response | 6.22E-03 | 9.50E-02 | 11 |
| GO:0034620 Cellular response to unfolded protein | 6.30E-03 | 9.50E-02 | 11 |
| GO:0019903 Protein phosphatase binding | 6.35E-03 | 9.50E-02 | 16 |
| GO:0016050 Vesicle organization | 6.37E-03 | 9.50E-02 | 25 |
| GO:0009144 Purine nucleoside triphosphate metabolic process | 6.38E-03 | 9.50E-02 | 64 |
| GO:0044087 Regulation of cellular component biogenesis | 6.94E-03 | 9.50E-02 | 62 |
| GO:0031226 Intrinsic component of plasma membrane | 6.97E-03 | 9.50E-02 | 58 |
| GO:0048193 Golgi vesicle transport | 6.98E-03 | 9.50E-02 | 39 |
| GO:0005911 Cell-cell junction | 7.07E-03 | 9.50E-02 | 46 |

| Supplemental Table S3<br>Annotation: Ontology Enriched in Old Lens Cortex relative to Young Lens Cortex | P-Value | FDR | Number of Proteins with Annotation in Dataset |
|---|---|---|---|
| GO:0003824 Catalytic activity | <1E-06 | 1.25E-02 | 614 |
| GO:0005737 Cytoplasm | <1E-06 | 1.25E-02 | 508 |
| GO:0005829 Cytosol | <1E-06 | 1.25E-02 | 504 |
| GO:0005975 Carbohydrate metabolic process | <1E-06 | 1.25E-02 | 106 |
| GO:0005996 Monosaccharide metabolic process | <1E-06 | 1.25E-02 | 61 |
| GO:0006006 Glucose metabolic process | <1E-06 | 1.25E-02 | 50 |
| GO:0006082 Organic acid metabolic process | <1E-06 | 1.25E-02 | 128 |
| GO:0006520 Cellular amino acid metabolic process | <1E-06 | 1.25E-02 | 75 |
| GO:0006558 L-phenylalanine metabolic process | <1E-06 | 1.25E-02 | 6 |
| GO:0006559 L-phenylalanine catabolic process | <1E-06 | 1.25E-02 | 6 |
| GO:0006807 Nitrogen compound metabolic process | <1E-06 | 1.25E-02 | 362 |
| GO:0008152 Metabolic process | <1E-06 | 1.25E-02 | 736 |
| GO:0009058 Biosynthetic process | <1E-06 | 1.25E-02 | 261 |
| GO:0009063 Cellular amino acid catabolic process | <1E-06 | 1.25E-02 | 16 |
| GO:0009072 Aromatic amino acid family metabolic process | <1E-06 | 1.25E-02 | 6 |
| GO:0009074 Aromatic amino acid family catabolic process | <1E-06 | 1.25E-02 | 6 |
| GO:0016052 Carbohydrate catabolic process | <1E-06 | 1.25E-02 | 38 |
| GO:0016054 Organic acid catabolic process | <1E-06 | 1.25E-02 | 24 |
| GO:0016740 Transferase activity | <1E-06 | 1.25E-02 | 142 |
| GO:0019318 Hexose metabolic process | <1E-06 | 1.25E-02 | 56 |
| GO:0019320 Hexose catabolic process | <1E-06 | 1.25E-02 | 28 |
| GO:0019752 Carboxylic acid metabolic process | <1E-06 | 1.25E-02 | 122 |
| GO:0043167 Ion binding | <1E-06 | 1.25E-02 | 530 |
| GO:0043436 Oxoacid metabolic process | <1E-06 | 1.25E-02 | 126 |
| GO:0044237 Cellular metabolic process | <1E-06 | 1.25E-02 | 678 |
| GO:0044238 Primary metabolic process | <1E-06 | 1.25E-02 | 655 |
| GO:0044249 Cellular biosynthetic process | <1E-06 | 1.25E-02 | 226 |
| GO:0044281 Small molecule metabolic process | <1E-06 | 1.25E-02 | 368 |
| GO:0044282 Small molecule catabolic process | <1E-06 | 1.25E-02 | 38 |
| GO:0046365 Monosaccharide catabolic process | <1E-06 | 1.25E-02 | 31 |
| GO:0046395 Carboxylic acid catabolic process | <1E-06 | 1.25E-02 | 24 |
| GO:0046872 Metal ion binding | <1E-06 | 1.25E-02 | 247 |
| GO:0071704 Organic substance metabolic process | <1E-06 | 1.25E-02 | 682 |
| GO:1901564 Organonitrogen compound metabolic process | <1E-06 | 1.25E-02 | 202 |
| GO:1901566 Organonitrogen compound biosynthetic process | <1E-06 | 1.25E-02 | 66 |
| GO:1901576 Organic substance biosynthetic process | <1E-06 | 1.25E-02 | 258 |
| GO:1901605 Alpha-amino acid metabolic process | <1E-06 | 1.25E-02 | 33 |
| GO:1901606 Alpha-amino acid catabolic process | <1E-06 | 1.25E-02 | 14 |
| GO:1902221 Erythrose 4-phosphate/phosphoenolpyruvate family amino acid metabolic process | <1E-06 | 1.25E-02 | 6 |

| Supplemental Table S3<br>Annotation: Ontology Enriched in Old Lens Cortex relative to Young Lens Cortex | P-Value | FDR | Number of Proteins with Annotation in Dataset |
|---|---|---|---|
| GO:1902222 Erythrose 4-phosphate/phosphoenolpyruvate family amino acid catabolic process | <1E-06 | 1.25E-02 | 6 |
| GO:0005212 Structural constituent of eye lens | 1.00E-06 | 1.25E-02 | 17 |
| GO:0016614 Oxidoreductase activity, acting on CH-OH group of donors | 1.00E-06 | 1.25E-02 | 29 |
| GO:0034641 Cellular nitrogen compound metabolic process | 1.00E-06 | 1.25E-02 | 335 |
| GO:0043169 Cation binding | 2.00E-06 | 1.25E-02 | 253 |
| GO:0016616 Oxidoreductase activity, acting on the CH-OH group of donors, NAD or NADP as acceptor | 2.00E-06 | 1.25E-02 | 29 |
| GO:0006007 Glucose catabolic process | 2.00E-06 | 1.25E-02 | 25 |
| GO:0042803 Protein homodimerization activity | 4.00E-06 | 1.25E-02 | 90 |
| GO:0042180 Cellular ketone metabolic process | 6.00E-06 | 1.25E-02 | 8 |
| GO:0044283 Small molecule biosynthetic process | 6.00E-06 | 1.25E-02 | 57 |
| GO:0042802 Identical protein binding | 1.20E-05 | 1.25E-02 | 134 |
| GO:0050661 NADP binding | 1.60E-05 | 1.25E-02 | 14 |
| GO:0046394 Carboxylic acid biosynthetic process | 3.90E-05 | 1.25E-02 | 39 |
| GO:0016053 Organic acid biosynthetic process | 5.20E-05 | 1.25E-02 | 39 |
| GO:0006570 Tyrosine metabolic process | 6.20E-05 | 1.25E-02 | 3 |
| GO:0006572 Tyrosine catabolic process | 7.10E-05 | 1.25E-02 | 3 |
| GO:0016810 Hydrolase activity, acting on carbon-nitrogen (but not peptide) bonds | 7.20E-05 | 1.25E-02 | 16 |
| GO:0006739 NADP metabolic process | 7.70E-05 | 1.25E-02 | 11 |
| GO:0002088 Lens development in camera-type eye | 9.20E-05 | 1.25E-02 | 9 |
| GO:0043094 Cellular metabolic compound salvage | 9.40E-05 | 1.25E-02 | 9 |
| GO:0016829 Lyase activity | 9.60E-05 | 1.25E-02 | 30 |
| GO:0016853 Isomerase activity | 9.60E-05 | 1.25E-02 | 34 |
| GO:0000287 Magnesium ion binding | 9.90E-05 | 1.25E-02 | 34 |
| GO:0006096 Glycolytic process | 1.02E-04 | 1.25E-02 | 20 |
| GO:0008652 Cellular amino acid biosynthetic process | 1.04E-04 | 1.25E-02 | 23 |
| GO:0006790 Sulfur compound metabolic process | 1.20E-04 | 1.25E-02 | 33 |
| GO:1901575 Organic substance catabolic process | 1.32E-04 | 1.25E-02 | 253 |
| GO:0019321 Pentose metabolic process | 1.47E-04 | 1.25E-02 | 8 |
| GO:0006144 Purine nucleobase metabolic process | 1.64E-04 | 1.25E-02 | 11 |
| GO:0009056 Catabolic process | 1.66E-04 | 1.25E-02 | 278 |
| GO:0006081 Cellular aldehyde metabolic process | 1.72E-04 | 1.25E-02 | 17 |
| GO:0016772 Transferase activity, transferring phosphorus-containing groups | 1.95E-04 | 2.20E-02 | 80 |
| GO:0044271 Cellular nitrogen compound biosynthetic process | 2.17E-04 | 2.20E-02 | 121 |
| GO:0016701 Oxidoreductase activity, acting on single donors with incorporation of molecular oxygen | 2.33E-04 | 2.20E-02 | 2 |
| GO:1901137 Carbohydrate derivative biosynthetic process | 2.33E-04 | 2.20E-02 | 44 |
| GO:0009112 Nucleobase metabolic process | 2.37E-04 | 2.20E-02 | 17 |
| GO:0016702 Oxidoreductase activity, acting on single donors with incorporation of molecular oxygen, incorporation of two atoms of oxygen | 2.39E-04 | 2.20E-02 | 2 |
| GO:1901661 Quinone metabolic process | 2.42E-04 | 2.20E-02 | 7 |

| Supplemental Table S3<br>Annotation: Ontology Enriched in Old Lens Cortex relative to Young Lens Cortex | P-Value | FDR | Number of Proteins with Annotation in Dataset |
|---|---|---|---|
| GO:0016903 Oxidoreductase activity, acting on the aldehyde or oxo group of donors | 2.50E-04 | 2.20E-02 | 11 |
| GO:0006044 N-acetylglucosamine metabolic process | 2.53E-04 | 2.20E-02 | 6 |
| GO:0006575 Cellular modified amino acid metabolic process | 2.69E-04 | 2.20E-02 | 33 |
| GO:0044262 Cellular carbohydrate metabolic process | 2.78E-04 | 2.20E-02 | 31 |
| GO:0090407 Organophosphate biosynthetic process | 3.83E-04 | 2.61E-02 | 64 |
| GO:0036094 Small molecule binding | 3.99E-04 | 2.61E-02 | 348 |
| GO:0008237 Metallopeptidase activity | 4.00E-04 | 2.61E-02 | 19 |
| GO:0042558 Pteridine-containing compound metabolic process | 4.04E-04 | 2.61E-02 | 6 |
| GO:0016861 Intramolecular oxidoreductase activity, interconverting aldoses and ketoses | 4.06E-04 | 2.61E-02 | 8 |
| GO:1901360 Organic cyclic compound metabolic process | 4.87E-04 | 2.61E-02 | 327 |
| GO:0005977 Glycogen metabolic process | 4.94E-04 | 2.61E-02 | 13 |
| GO:0044264 Cellular polysaccharide metabolic process | 4.94E-04 | 2.61E-02 | 13 |
| GO:0044042 Glucan metabolic process | 5.06E-04 | 2.61E-02 | 13 |
| GO:0005976 Polysaccharide metabolic process | 5.08E-04 | 2.61E-02 | 13 |
| GO:0006073 Cellular glucan metabolic process | 5.12E-04 | 2.61E-02 | 13 |
| GO:1901071 Glucosamine-containing compound metabolic process | 5.16E-04 | 2.61E-02 | 8 |
| GO:0019362 Pyridine nucleotide metabolic process | 5.25E-04 | 2.61E-02 | 18 |
| GO:0046496 Nicotinamide nucleotide metabolic process | 5.36E-04 | 2.61E-02 | 18 |
| GO:0072524 Pyridine-containing compound metabolic process | 5.59E-04 | 2.61E-02 | 18 |
| GO:0006725 Cellular aromatic compound metabolic process | 6.08E-04 | 2.61E-02 | 305 |
| GO:1901293 Nucleoside phosphate biosynthetic process | 6.21E-04 | 2.61E-02 | 38 |
| GO:0009165 Nucleotide biosynthetic process | 6.31E-04 | 2.61E-02 | 38 |
| GO:0072522 Purine-containing compound biosynthetic process | 6.35E-04 | 2.61E-02 | 26 |
| GO:0016620 Oxidoreductase activity, acting on the aldehyde or oxo group of donors, NAD or NADP as acceptor | 7.24E-04 | 2.61E-02 | 10 |
| GO:0005524 ATP binding | 7.32E-04 | 2.61E-02 | 187 |
| GO:0016051 Carbohydrate biosynthetic process | 7.34E-04 | 2.61E-02 | 37 |
| GO:0016774 Phosphotransferase activity, carboxyl group as acceptor | 7.73E-04 | 2.61E-02 | 2 |
| GO:0016491 Oxidoreductase activity | 7.78E-04 | 3.28E-02 | 95 |
| GO:0043603 Cellular amide metabolic process | 8.14E-04 | 3.28E-02 | 29 |
| GO:0004618 Phosphoglycerate kinase activity | 8.43E-04 | 3.28E-02 | 2 |
| GO:0008235 Metalloexopeptidase activity | 8.71E-04 | 3.28E-02 | 5 |
| GO:0034660 ncRNA metabolic process | 9.00E-04 | 3.28E-02 | 31 |
| GO:0016301 Kinase activity | 9.16E-04 | 3.28E-02 | 72 |
| GO:0006740 NADPH regeneration | 9.58E-04 | 3.28E-02 | 7 |
| GO:0015949 Nucleobase-containing small molecule interconversion | 9.79E-04 | 3.82E-02 | 7 |
| GO:0008233 Peptidase activity | 9.85E-04 | 3.82E-02 | 66 |
| GO:0042559 Pteridine-containing compound biosynthetic process | 1.01E-03 | 3.82E-02 | 4 |
| GO:0044272 Sulfur compound biosynthetic process | 1.05E-03 | 3.82E-02 | 14 |

| **Supplemental Table S3**<br>**Annotation: Ontology Enriched in Old Lens Cortex relative to Young Lens Cortex** | **P-Value** | **FDR** | **Number of Proteins with Annotation in Dataset** |
|---|---|---|---|
| GO:0033559 Unsaturated fatty acid metabolic process | 1.08E-03 | 3.82E-02 | 11 |
| GO:0032559 Adenyl ribonucleotide binding | 1.08E-03 | 3.82E-02 | 192 |
| GO:1901568 Fatty acid derivative metabolic process | 1.08E-03 | 3.82E-02 | 11 |
| GO:0006690 Icosanoid metabolic process | 1.10E-03 | 3.82E-02 | 11 |
| GO:0030554 Adenyl nucleotide binding | 1.10E-03 | 3.82E-02 | 195 |
| GO:0019322 Pentose biosynthetic process | 1.17E-03 | 4.55E-02 | 4 |
| GO:0043412 Macromolecule modification | 1.34E-03 | 5.07E-02 | 199 |
| GO:0016788 Hydrolase activity, acting on ester bonds | 1.37E-03 | 5.07E-02 | 68 |
| GO:0008238 Exopeptidase activity | 1.42E-03 | 5.07E-02 | 19 |
| GO:0036211 Protein modification process | 1.52E-03 | 5.07E-02 | 197 |
| GO:0006464 Cellular protein modification process | 1.53E-03 | 5.07E-02 | 197 |
| GO:0061134 Peptidase regulator activity | 1.54E-03 | 5.07E-02 | 17 |
| GO:0019262 N-acetylneuraminate catabolic process | 1.73E-03 | 5.76E-02 | 3 |
| GO:0005997 Xylulose metabolic process | 1.81E-03 | 6.43E-02 | 3 |
| GO:0006040 Amino sugar metabolic process | 1.88E-03 | 6.85E-02 | 9 |
| GO:0004177 Aminopeptidase activity | 1.94E-03 | 6.85E-02 | 9 |
| GO:0051260 Protein homooligomerization | 2.01E-03 | 6.85E-02 | 25 |
| GO:0016570 Histone modification | 2.18E-03 | 6.85E-02 | 11 |
| GO:0009055 Electron transfer activity | 2.21E-03 | 7.48E-02 | 14 |
| GO:0019438 Aromatic compound biosynthetic process | 2.32E-03 | 7.60E-02 | 113 |
| GO:0004033 Aldo-keto reductase (NADP) activity | 2.34E-03 | 7.60E-02 | 5 |
| GO:0016765 Transferase activity, transferring alkyl or aryl (other than methyl) groups | 2.47E-03 | 7.60E-02 | 14 |
| GO:0018130 Heterocycle biosynthetic process | 2.47E-03 | 7.60E-02 | 112 |
| GO:0016846 Carbon-sulfur lyase activity | 2.58E-03 | 7.60E-02 | 2 |
| GO:0033293 Monocarboxylic acid binding | 2.66E-03 | 7.60E-02 | 8 |
| GO:0006720 Isoprenoid metabolic process | 2.76E-03 | 7.60E-02 | 10 |
| GO:0043101 Purine-containing compound salvage | 2.78E-03 | 7.60E-02 | 3 |
| GO:0005634 Nucleus | 2.82E-03 | 7.60E-02 | 322 |
| GO:0046348 Amino sugar catabolic process | 2.83E-03 | 7.60E-02 | 6 |
| GO:0042440 Pigment metabolic process | 2.87E-03 | 7.60E-02 | 9 |
| GO:0000096 Sulfur amino acid metabolic process | 2.93E-03 | 7.60E-02 | 14 |
| GO:1901362 Organic cyclic compound biosynthetic process | 2.98E-03 | 7.60E-02 | 129 |
| GO:0019205 Nucleobase-containing compound kinase activity | 3.08E-03 | 7.60E-02 | 5 |
| GO:0007286 Spermatid development | 3.17E-03 | 7.60E-02 | 3 |
| GO:0016857 Racemase and epimerase activity, acting on carbohydrates and derivatives | 3.23E-03 | 7.60E-02 | 3 |
| GO:0046483 Heterocycle metabolic process | 3.28E-03 | 7.60E-02 | 297 |
| GO:0016854 Racemase and epimerase activity | 3.28E-03 | 7.60E-02 | 3 |
| GO:0044106 Cellular amine metabolic process | 3.33E-03 | 7.60E-02 | 12 |
| GO:0006576 Cellular biogenic amine metabolic process | 3.34E-03 | 7.60E-02 | 12 |

| **Supplemental Table S3**<br>**Annotation: Ontology Enriched in Old Lens Cortex relative to Young Lens Cortex** | **P-Value** | **FDR** | **Number of Proteins with Annotation in Dataset** |
|---|---|---|---|
| GO:0004180 Carboxypeptidase activity | 3.39E-03 | 7.60E-02 | 4 |
| GO:0042182 Ketone catabolic process | 3.43E-03 | 7.60E-02 | 2 |
| GO:0009308 Amine metabolic process | 3.46E-03 | 7.60E-02 | 12 |
| GO:0008270 Zinc ion binding | 3.58E-03 | 7.60E-02 | 51 |
| GO:0016769 Transferase activity, transferring nitrogenous groups | 3.59E-03 | 7.91E-02 | 5 |
| GO:0016866 Intramolecular transferase activity | 3.68E-03 | 7.91E-02 | 10 |
| GO:1901135 Carbohydrate derivative metabolic process | 3.75E-03 | 7.91E-02 | 123 |
| GO:0051287 NAD binding | 3.79E-03 | 7.91E-02 | 15 |
| GO:0006098 Pentose-phosphate shunt | 3.86E-03 | 7.91E-02 | 6 |
| GO:0006012 Galactose metabolic process | 3.97E-03 | 8.15E-02 | 5 |
| GO:0004812 Aminoacyl-tRNA ligase activity | 4.00E-03 | 8.15E-02 | 20 |
| GO:0009584 Detection of visible light | 4.00E-03 | 8.15E-02 | 8 |
| GO:0007602 Phototransduction | 4.00E-03 | 8.15E-02 | 8 |
| GO:0009583 Detection of light stimulus | 4.01E-03 | 8.15E-02 | 8 |
| GO:0043039 tRNA aminoacylation | 4.04E-03 | 8.15E-02 | 20 |
| GO:0046148 Pigment biosynthetic process | 4.07E-03 | 8.51E-02 | 8 |
| GO:0006796 Phosphate-containing compound metabolic process | 4.08E-03 | 8.51E-02 | 216 |
| GO:0006418 tRNA aminoacylation for protein translation | 4.08E-03 | 8.51E-02 | 20 |
| GO:0016875 Ligase activity, forming carbon-oxygen bonds | 4.10E-03 | 8.51E-02 | 20 |
| GO:0007603 Phototransduction, visible light | 4.15E-03 | 8.90E-02 | 8 |
| GO:0043038 Amino acid activation | 4.17E-03 | 8.90E-02 | 20 |
| GO:0004181 Metallocarboxypeptidase activity | 4.19E-03 | 8.90E-02 | 2 |

| Supplemental Table S4<br>Annotation: Ontology Enriched in Young Lens Outer Nucleus relative to Old Lens Outer Nucleus | P-Value | FDR | Number of Proteins with Annotation in Dataset |
|---|---|---|---|
| GO:0019001 Guanyl nucleotide binding | <1E-06 | 1.35E-02 | 75 |
| GO:0034329 Cell junction assembly | <1E-06 | 1.35E-02 | 23 |
| GO:0034330 Cell junction organization | <1E-06 | 1.35E-02 | 26 |
| GO:0005525 GTP binding | <1E-06 | 1.35E-02 | 75 |
| GO:0005886 Plasma membrane | <1E-06 | 1.35E-02 | 204 |
| GO:0007155 Cell adhesion | <1E-06 | 1.35E-02 | 43 |
| GO:0007264 Small GTPase mediated signal transduction | <1E-06 | 1.35E-02 | 58 |
| GO:0016020 Membrane | <1E-06 | 1.35E-02 | 344 |
| GO:0016021 Integral component of membrane | <1E-06 | 1.35E-02 | 113 |
| GO:0019898 Extrinsic component of membrane | <1E-06 | 1.35E-02 | 19 |
| GO:0022610 Biological adhesion | <1E-06 | 1.35E-02 | 43 |
| GO:0031224 Intrinsic component of membrane | <1E-06 | 1.35E-02 | 119 |
| GO:0032561 Guanyl ribonucleotide binding | <1E-06 | 1.35E-02 | 75 |
| GO:0045216 Cell-cell junction organization | 2.00E-06 | 1.35E-02 | 23 |
| GO:0031410 Cytoplasmic vesicle | 3.00E-06 | 1.35E-02 | 71 |
| GO:0031012 Extracellular matrix | 6.00E-06 | 1.35E-02 | 14 |
| GO:0031982 Vesicle | 9.00E-06 | 1.35E-02 | 99 |
| GO:0003924 GTPase activity | 1.00E-05 | 1.35E-02 | 55 |
| GO:0017111 Nucleoside-triphosphatase activity | 1.70E-05 | 1.35E-02 | 102 |
| GO:0050817 Coagulation | 4.70E-05 | 1.35E-02 | 44 |
| GO:0016462 Pyrophosphatase activity | 4.80E-05 | 1.35E-02 | 108 |
| GO:0007596 Blood coagulation | 5.00E-05 | 1.35E-02 | 44 |
| GO:0016818 Hydrolase activity, acting on acid anhydrides, in phosphorus-containing anhydrides | 5.10E-05 | 1.35E-02 | 109 |
| GO:0007599 Hemostasis | 5.90E-05 | 1.35E-02 | 46 |
| GO:0016817 Hydrolase activity, acting on acid anhydrides | 6.80E-05 | 1.35E-02 | 109 |
| GO:0006928 Movement of cell or subcellular component | 6.80E-05 | 1.35E-02 | 103 |
| GO:0032502 Developmental process | 7.00E-05 | 1.35E-02 | 232 |
| GO:0007156 Homophilic cell adhesion via plasma membrane adhesion molecules | 1.45E-04 | 1.35E-02 | 5 |
| GO:0007411 Axon guidance | 1.48E-04 | 1.35E-02 | 42 |
| GO:0097485 Neuron projection guidance | 1.48E-04 | 1.35E-02 | 42 |
| GO:0005834 Heterotrimeric G-protein complex | 1.69E-04 | 1.35E-02 | 5 |
| GO:0050794 Regulation of cellular process | 1.69E-04 | 1.35E-02 | 512 |
| GO:0009986 Cell surface | 1.73E-04 | 1.35E-02 | 21 |
| GO:0031344 Regulation of cell projection organization | 2.05E-04 | 1.35E-02 | 23 |
| GO:0019897 Extrinsic component of plasma membrane | 2.28E-04 | 1.35E-02 | 12 |
| GO:0008366 Axon ensheathment | 2.52E-04 | 1.35E-02 | 4 |
| GO:0007272 Ensheathment of neurons | 2.79E-04 | 1.35E-02 | 4 |
| GO:0031225 Anchored component of membrane | 3.05E-04 | 1.35E-02 | 6 |
| GO:0006813 Potassium ion transport | 3.27E-04 | 1.35E-02 | 4 |

| Supplemental Table S4<br>Annotation: Ontology Enriched in Young Lens Outer Nucleus relative to Old Lens Outer Nucleus | P-Value | FDR | Number of Proteins with Annotation in Dataset |
|---|---|---|---|
| GO:0007157 Heterophilic cell-cell adhesion via plasma membrane cell adhesion molecules | 3.34E-04 | 1.35E-02 | 4 |
| GO:0007267 Cell-cell signaling | 3.35E-04 | 1.35E-02 | 34 |
| GO:0045121 Membrane raft | 3.36E-04 | 1.35E-02 | 19 |
| GO:0008324 Cation transmembrane transporter activity | 3.83E-04 | 1.35E-02 | 17 |
| GO:0046039 GTP metabolic process | 3.86E-04 | 1.35E-02 | 29 |
| GO:0032989 Cellular component morphogenesis | 3.92E-04 | 1.35E-02 | 32 |
| GO:0065007 Biological regulation | 4.25E-04 | 1.35E-02 | 564 |
| GO:0048770 Pigment granule | 5.05E-04 | 1.35E-02 | 31 |
| GO:1901068 Guanosine-containing compound metabolic process | 5.23E-04 | 1.35E-02 | 31 |
| GO:0042470 Melanosome | 5.25E-04 | 1.35E-02 | 31 |
| GO:0050767 Regulation of neurogenesis | 5.28E-04 | 1.35E-02 | 26 |
| GO:0046873 Metal ion transmembrane transporter activity | 5.31E-04 | 1.35E-02 | 7 |
| GO:0051960 Regulation of nervous system development | 5.40E-04 | 1.35E-02 | 26 |
| GO:0048731 System development | 5.42E-04 | 1.35E-02 | 31 |
| GO:0050900 Leukocyte migration | 5.46E-04 | 1.35E-02 | 17 |
| GO:0034332 Adherens junction organization | 5.79E-04 | 1.35E-02 | 16 |
| GO:0050896 Response to stimulus | 6.31E-04 | 1.35E-02 | 444 |
| GO:0007165 Signal transduction | 6.39E-04 | 1.35E-02 | 286 |
| GO:0050789 Regulation of biological process | 6.55E-04 | 1.35E-02 | 534 |
| GO:0016323 Basolateral plasma membrane | 6.90E-04 | 1.35E-02 | 22 |
| GO:0019829 ATPase-coupled cation transmembrane transporter activity | 7.31E-04 | 1.35E-02 | 9 |
| GO:0031234 Extrinsic component of cytoplasmic side of plasma membrane | 7.31E-04 | 1.35E-02 | 6 |
| GO:0042625 ATPase-coupled ion transmembrane transporter activity | 7.44E-04 | 1.35E-02 | 9 |
| GO:0030054 Cell junction | 7.53E-04 | 1.35E-02 | 67 |
| GO:0050808 Synapse organization | 7.93E-04 | 1.35E-02 | 10 |
| GO:0019003 GDP binding | 8.21E-04 | 1.35E-02 | 18 |
| GO:1901069 Guanosine-containing compound catabolic process | 8.58E-04 | 2.17E-02 | 26 |
| GO:0060089 Molecular transducer activity | 9.12E-04 | 2.17E-02 | 38 |
| GO:0030198 Extracellular matrix organization | 9.16E-04 | 2.17E-02 | 13 |
| GO:0009205 Purine ribonucleoside triphosphate metabolic process | 9.55E-04 | 2.17E-02 | 44 |
| GO:0009653 Anatomical structure morphogenesis | 9.59E-04 | 2.17E-02 | 65 |
| GO:0043062 Extracellular structure organization | 9.65E-04 | 2.17E-02 | 13 |
| GO:0003382 Epithelial cell morphogenesis | 9.86E-04 | 2.17E-02 | 4 |
| GO:0009199 Ribonucleoside triphosphate metabolic process | 1.01E-03 | 2.17E-02 | 44 |
| GO:0048856 Anatomical structure development | 1.06E-03 | 2.17E-02 | 151 |
| GO:1900542 Regulation of purine nucleotide metabolic process | 1.09E-03 | 2.17E-02 | 19 |
| GO:0043194 Axon initial segment | 1.10E-03 | 2.17E-02 | 3 |
| GO:0000902 Cell morphogenesis | 1.15E-03 | 2.17E-02 | 14 |
| GO:0040011 Locomotion | 1.20E-03 | 2.17E-02 | 49 |

| **Supplemental Table S4**<br>**Annotation: Ontology Enriched in Young Lens Outer Nucleus relative to Old Lens Outer Nucleus** | **P-Value** | **FDR** | **Number of Proteins with Annotation in Dataset** |
|---|---|---|---|
| GO:0023052 Signaling | 1.27E-03 | 2.17E-02 | 38 |
| GO:0009144 Purine nucleoside triphosphate metabolic process | 1.31E-03 | 2.17E-02 | 47 |
| GO:0051056 Regulation of small GTPase mediated signal transduction | 1.41E-03 | 3.19E-02 | 24 |
| GO:0048869 Cellular developmental process | 1.44E-03 | 3.19E-02 | 98 |
| GO:0008104 Protein localization | 1.51E-03 | 4.21E-02 | 40 |
| GO:0033010 Paranodal junction | 1.56E-03 | 5.05E-02 | 2 |
| GO:0031226 Intrinsic component of plasma membrane | 1.57E-03 | 5.05E-02 | 36 |
| GO:0030139 Endocytic vesicle | 1.58E-03 | 5.05E-02 | 11 |
| GO:0065008 Regulation of biological quality | 1.58E-03 | 5.05E-02 | 220 |
| GO:0033270 Paranode region of axon | 1.59E-03 | 5.83E-02 | 3 |
| GO:0009141 Nucleoside triphosphate metabolic process | 1.62E-03 | 5.83E-02 | 48 |
| GO:0050878 Regulation of body fluid levels | 1.66E-03 | 5.83E-02 | 55 |
| GO:0042626 ATPase-coupled transmembrane transporter activity | 1.80E-03 | 7.62E-02 | 10 |
| GO:0051179 Localization | 1.90E-03 | 7.89E-02 | 48 |
| GO:0045664 Regulation of neuron differentiation | 1.90E-03 | 7.89E-02 | 20 |
| GO:0030154 Cell differentiation | 1.95E-03 | 7.89E-02 | 57 |
| GO:0033268 Node of ranvier | 2.01E-03 | 7.89E-02 | 2 |
| GO:0010927 Cellular component assembly involved in morphogenesis | 2.02E-03 | 7.89E-02 | 7 |
| GO:0015081 Sodium ion transmembrane transporter activity | 2.10E-03 | 7.89E-02 | 5 |
| GO:0033036 Macromolecule localization | 2.25E-03 | 7.89E-02 | 41 |
| GO:0060560 Developmental growth involved in morphogenesis | 2.27E-03 | 7.89E-02 | 3 |
| GO:0045761 Regulation of adenylate cyclase activity | 2.30E-03 | 8.20E-02 | 7 |
| GO:0031279 Regulation of cyclase activity | 2.31E-03 | 8.20E-02 | 7 |
| GO:0005391 P-type sodium:potassium-exchanging transporter activity | 2.36E-03 | 8.20E-02 | 3 |
| GO:0015662 P-type ion transporter activity | 2.39E-03 | 8.20E-02 | 3 |
| GO:0098533 ATPase dependent transmembrane transport complex | 2.40E-03 | 8.20E-02 | 3 |
| GO:0005890 Sodium:potassium-exchanging ATPase complex | 2.40E-03 | 8.20E-02 | 3 |
| GO:0008556 P-type potassium transmembrane transporter activity | 2.41E-03 | 8.20E-02 | 3 |
| GO:0090533 Cation-transporting ATPase complex | 2.44E-03 | 8.20E-02 | 3 |
| GO:0015079 Potassium ion transmembrane transporter activity | 2.51E-03 | 8.59E-02 | 3 |
| GO:0002685 Regulation of leukocyte migration | 2.59E-03 | 8.59E-02 | 5 |
| GO:0001525 Angiogenesis | 2.67E-03 | 8.59E-02 | 16 |
| GO:0051339 Regulation of lyase activity | 2.68E-03 | 8.59E-02 | 8 |
| GO:0045202 Synapse | 2.76E-03 | 8.59E-02 | 17 |
| GO:0016324 Apical plasma membrane | 2.88E-03 | 8.59E-02 | 16 |
| GO:0005768 Endosome | 2.91E-03 | 9.02E-02 | 34 |
| GO:0050870 Positive regulation of t cell activation | 2.94E-03 | 9.02E-02 | 13 |
| GO:0060491 Regulation of cell projection assembly | 2.97E-03 | 9.02E-02 | 8 |
| GO:0042552 Myelination | 3.08E-03 | 9.02E-02 | 3 |

| **Supplemental Table S4**<br>**Annotation: Ontology Enriched in Young Lens Outer Nucleus relative to Old Lens Outer Nucleus** | **P-Value** | **FDR** | **Number of Proteins with Annotation in Dataset** |
|---|---|---|---|
| GO:0086080 Protein binding involved in heterotypic cell-cell adhesion | 3.22E-03 | 9.49E-02 | 2 |
| GO:0045162 Clustering of voltage-gated sodium channels | 3.26E-03 | 9.49E-02 | 2 |
| GO:0050839 Cell adhesion molecule binding | 3.28E-03 | 9.49E-02 | 10 |
| GO:0010975 Regulation of neuron projection development | 3.30E-03 | 9.49E-02 | 17 |

| Supplemental Table S5<br>**Annotation: Ontology Enriched in Old Lens Outer Nucleus relative to Young Lens Outer Nucleus** | P-Value | FDR | Number of Proteins with Annotation in Dataset |
|---|---|---|---|
| GO:0003824 Catalytic activity | <1E-06 | 2.94E-02 | 517 |
| GO:0008152 Metabolic process | <1E-06 | 2.94E-02 | 604 |
| GO:0016491 Oxidoreductase activity | <1E-06 | 2.94E-02 | 81 |
| GO:0043170 Macromolecule metabolic process | <1E-06 | 2.94E-02 | 357 |
| GO:0044237 Cellular metabolic process | <1E-06 | 2.94E-02 | 559 |
| GO:0044238 Primary metabolic process | <1E-06 | 2.94E-02 | 543 |
| GO:0044281 Small molecule metabolic process | <1E-06 | 2.94E-02 | 312 |
| GO:0071704 Organic substance metabolic process | <1E-06 | 2.94E-02 | 565 |
| GO:0019538 Protein metabolic process | 2.00E-06 | 2.94E-02 | 263 |
| GO:0005829 Cytosol | 2.00E-06 | 2.94E-02 | 436 |
| GO:0044260 Cellular macromolecule metabolic process | 3.00E-06 | 2.94E-02 | 330 |
| GO:1902494 Catalytic complex | 6.00E-06 | 2.94E-02 | 72 |
| GO:0005739 Mitochondrion | 1.20E-05 | 2.94E-02 | 142 |
| GO:0006464 Cellular protein modification process | 1.50E-05 | 2.94E-02 | 165 |
| GO:0036211 Protein modification process | 1.60E-05 | 2.94E-02 | 165 |
| GO:0043412 Macromolecule modification | 2.50E-05 | 2.94E-02 | 167 |
| GO:0044267 Cellular protein metabolic process | 3.40E-05 | 2.94E-02 | 234 |
| GO:1990204 Oxidoreductase complex | 4.10E-05 | 2.94E-02 | 11 |
| GO:0009058 Biosynthetic process | 4.10E-05 | 2.94E-02 | 228 |
| GO:1901576 Organic substance biosynthetic process | 4.80E-05 | 2.94E-02 | 226 |
| GO:0044249 Cellular biosynthetic process | 7.70E-05 | 2.94E-02 | 198 |
| GO:0005654 Nucleoplasm | 1.48E-04 | 2.94E-02 | 82 |
| GO:0006807 Nitrogen compound metabolic process | 1.73E-04 | 2.94E-02 | 312 |
| GO:0034641 Cellular nitrogen compound metabolic process | 1.91E-04 | 2.94E-02 | 289 |
| GO:0006082 Organic acid metabolic process | 1.95E-04 | 2.94E-02 | 114 |
| GO:0019752 Carboxylic acid metabolic process | 2.32E-04 | 2.94E-02 | 110 |
| GO:0043436 Oxoacid metabolic process | 2.50E-04 | 2.94E-02 | 112 |
| GO:0019318 Hexose metabolic process | 3.03E-04 | 2.94E-02 | 53 |
| GO:0031966 Mitochondrial membrane | 3.25E-04 | 2.94E-02 | 39 |
| GO:0005743 Mitochondrial inner membrane | 4.40E-04 | 4.76E-02 | 28 |
| GO:0019866 Organelle inner membrane | 4.42E-04 | 4.76E-02 | 28 |
| GO:0005747 Mitochondrial respiratory chain complex I | 4.56E-04 | 4.76E-02 | 7 |
| GO:0045271 Respiratory chain complex I | 4.63E-04 | 4.76E-02 | 7 |
| GO:0030964 NADH dehydrogenase complex | 4.82E-04 | 4.76E-02 | 7 |
| GO:0016655 Oxidoreductase activity, acting on NAD(P)H, quinone or similar compound as acceptor | 5.80E-04 | 4.76E-02 | 11 |
| GO:1901575 Organic substance catabolic process | 6.19E-04 | 4.76E-02 | 217 |
| GO:0006091 Generation of precursor metabolites and energy | 6.25E-04 | 4.76E-02 | 63 |
| GO:0016651 Oxidoreductase activity, acting on NAD(P)H | 8.12E-04 | 7.69E-02 | 18 |
| GO:0005996 Monosaccharide metabolic process | 8.54E-04 | 7.69E-02 | 58 |

| Supplemental Table S5<br>Annotation: Ontology Enriched in Old Lens Outer Nucleus relative to Young Lens Outer Nucleus | P-Value | FDR | Number of Proteins with Annotation in Dataset |
|---|---|---|---|
| GO:0048145 Regulation of fibroblast proliferation | 8.97E-04 | 7.69E-02 | 9 |
| GO:0071482 Cellular response to light stimulus | 9.23E-04 | 7.69E-02 | 6 |
| GO:0005737 Cytoplasm | 9.84E-04 | 7.69E-02 | 432 |
| GO:0006006 Glucose metabolic process | 1.00E-03 | 7.69E-02 | 48 |
| GO:0004497 Monooxygenase activity | 1.18E-03 | 7.69E-02 | 5 |
| GO:0022900 Electron transport chain | 1.19E-03 | 7.69E-02 | 12 |
| GO:0016702 Oxidoreductase activity, acting on single donors with incorporation of molecular oxygen, incorporation of two atoms of oxygen | 1.19E-03 | 7.69E-02 | 4 |
| GO:0005506 Iron ion binding | 1.21E-03 | 7.69E-02 | 9 |
| GO:0022904 Respiratory electron transport chain | 1.21E-03 | 7.69E-02 | 12 |
| GO:0051213 Dioxygenase activity | 1.24E-03 | 7.69E-02 | 4 |
| GO:0048146 Positive regulation of fibroblast proliferation | 1.24E-03 | 7.69E-02 | 5 |
| GO:0005975 Carbohydrate metabolic process | 1.26E-03 | 7.69E-02 | 88 |
| GO:0016701 Oxidoreductase activity, acting on single donors with incorporation of molecular oxygen | 1.30E-03 | 7.69E-02 | 4 |
| GO:0046906 Tetrapyrrole binding | 1.32E-03 | 7.69E-02 | 5 |
| GO:0020037 Heme binding | 1.32E-03 | 7.69E-02 | 5 |
| GO:0051726 Regulation of cell cycle | 1.48E-03 | 7.69E-02 | 82 |
| GO:0032787 Monocarboxylic acid metabolic process | 1.49E-03 | 7.69E-02 | 36 |
| GO:0032446 Protein modification by small protein conjugation | 1.51E-03 | 7.69E-02 | 60 |
| GO:0015718 Monocarboxylic acid transport | 1.54E-03 | 9.09E-02 | 6 |
| GO:0071478 Cellular response to radiation | 1.55E-03 | 1.00E-01 | 7 |
| GO:1901360 Organic cyclic compound metabolic process | 1.56E-03 | 1.00E-01 | 282 |
| GO:0046483 Heterocycle metabolic process | 1.57E-03 | 1.00E-01 | 255 |
| GO:0006839 Mitochondrial transport | 1.58E-03 | 1.00E-01 | 13 |
| GO:0070647 Protein modification by small protein conjugation or removal | 1.58E-03 | 1.00E-01 | 76 |
| GO:0022402 Cell cycle process | 1.70E-03 | 1.00E-01 | 106 |
| GO:0050136 NADH dehydrogenase (quinone) activity | 1.71E-03 | 1.00E-01 | 6 |
| GO:0008137 NADH dehydrogenase (ubiquinone) activity | 1.76E-03 | 1.00E-01 | 6 |
| GO:0003954 NADH dehydrogenase activity | 1.77E-03 | 1.00E-01 | 6 |
| GO:0009056 Catabolic process | 1.78E-03 | 1.00E-01 | 234 |
| GO:0006725 Cellular aromatic compound metabolic process | 1.79E-03 | 1.00E-01 | 261 |
| GO:0090304 Nucleic acid metabolic process | 1.82E-03 | 1.00E-01 | 150 |
| GO:0016567 Protein ubiquitination | 1.83E-03 | 1.00E-01 | 57 |
| GO:0010803 Regulation of tumor necrosis factor-mediated signaling pathway | 1.91E-03 | 1.00E-01 | 3 |
| GO:1901566 Organonitrogen compound biosynthetic process | 1.91E-03 | 1.00E-01 | 63 |
| GO:0016070 RNA metabolic process | 1.95E-03 | 1.00E-01 | 131 |
| GO:0019371 Cyclooxygenase pathway | 2.36E-03 | 1.00E-01 | 4 |
| GO:0042168 Heme metabolic process | 2.86E-03 | 1.00E-01 | 4 |
| GO:0010467 Gene expression | 2.95E-03 | 1.00E-01 | 93 |

| Supplemental Table S5<br>Annotation: Ontology Enriched in Old Lens Outer Nucleus relative to Young Lens Outer Nucleus | P-Value | FDR | Number of Proteins with Annotation in Dataset |
|---|---|---|---|
| GO:0000082 G1/S transition of mitotic cell cycle | 3.02E-03 | 1.00E-01 | 43 |
| GO:0033554 Cellular response to stress | 3.14E-03 | 1.00E-01 | 101 |
| GO:0071013 Catalytic step 2 spliceosome | 3.19E-03 | 1.00E-01 | 2 |

| Supplemental Table S6<br>Annotation: Ontology Enriched in Young Lens Inner Nucleus relative to Old Lens Inner Nucleus | P-Value | FDR | Number of Proteins with Annotation in Dataset |
|---|---|---|---|
| GO:0005886 Plasma membrane | <1E-06 | 2.63E-02 | 174 |
| GO:0007155 Cell adhesion | <1E-06 | 2.63E-02 | 35 |
| GO:0016020 Membrane | <1E-06 | 2.63E-02 | 302 |
| GO:0022610 Biological adhesion | <1E-06 | 2.63E-02 | 35 |
| GO:0031224 Intrinsic component of membrane | <1E-06 | 2.63E-02 | 97 |
| GO:0034330 Cell junction organization | <1E-06 | 2.63E-02 | 23 |
| GO:0045121 Membrane raft | <1E-06 | 2.63E-02 | 16 |
| GO:0045216 Cell-cell junction organization | <1E-06 | 2.63E-02 | 20 |
| GO:0050870 Positive regulation of T-cell activation | <1E-06 | 2.63E-02 | 10 |
| GO:0002696 Positive regulation of leukocyte activation | 2.00E-06 | 2.63E-02 | 11 |
| GO:0050867 Positive regulation of cell activation | 2.00E-06 | 2.63E-02 | 11 |
| GO:0051251 Positive regulation of lymphocyte activation | 3.00E-06 | 2.63E-02 | 11 |
| GO:0031226 Intrinsic component of plasma membrane | 3.00E-06 | 2.63E-02 | 26 |
| GO:0034329 Cell junction assembly | 5.00E-06 | 2.63E-02 | 20 |
| GO:0050863 Regulation of T-cell activation | 9.00E-06 | 2.63E-02 | 15 |
| GO:0009986 Cell surface | 4.60E-05 | 2.63E-02 | 18 |
| GO:0001525 Angiogenesis | 5.70E-05 | 2.63E-02 | 13 |
| GO:0007411 Axon guidance | 8.20E-05 | 2.63E-02 | 40 |
| GO:0051249 Regulation of lymphocyte activation | 8.20E-05 | 2.63E-02 | 16 |
| GO:0031225 Anchored component of membrane | 9.10E-05 | 2.63E-02 | 6 |
| GO:0097485 Neuron projection guidance | 9.50E-05 | 2.63E-02 | 40 |
| GO:0002694 Regulation of leukocyte activation | 1.00E-04 | 2.63E-02 | 16 |
| GO:0016021 Integral component of membrane | 1.08E-04 | 2.63E-02 | 91 |
| GO:0019898 Extrinsic component of membrane | 1.09E-04 | 2.63E-02 | 16 |
| GO:0005887 Integral component of plasma membrane | 1.20E-04 | 2.63E-02 | 23 |
| GO:0032502 Developmental process | 1.21E-04 | 2.63E-02 | 193 |
| GO:0005901 Caveola | 1.23E-04 | 2.63E-02 | 8 |
| GO:0048646 Anatomical structure formation involved in morphogenesis | 1.42E-04 | 2.63E-02 | 27 |
| GO:0030054 Cell junction | 1.60E-04 | 2.63E-02 | 57 |
| GO:0050865 Regulation of cell activation | 1.74E-04 | 2.63E-02 | 18 |
| GO:0050817 Coagulation | 3.83E-04 | 2.63E-02 | 42 |
| GO:0007596 Blood coagulation | 4.29E-04 | 2.63E-02 | 42 |
| GO:0045785 Positive regulation of cell adhesion | 5.15E-04 | 2.63E-02 | 8 |

| **Supplemental Table S6**<br>**Annotation: Ontology Enriched in Young Lens Inner Nucleus relative to Old Lens Inner Nucleus** | **P-Value** | **FDR** | **Number of Proteins with Annotation in Dataset** |
|---|---|---|---|
| GO:0002684 Positive regulation of immune system process | 5.16E-04 | 2.63E-02 | 36 |
| GO:0034332 Adherens junction organization | 5.30E-04 | 7.69E-02 | 13 |

| **Supplemental Table S7**<br>**Annotation: Ontology Enriched in Old Lens Inner Nucleus relative to Young Lens Inner Nucleus** | **P-Value** | **FDR** | **Number of Proteins with Annotation in Dataset** |
|---|---|---|---|
| GO:0016491 Oxidoreductase activity | 9.60E-05 | 1.00E-01 | 78 |
| GO:0004867 Serine-type endopeptidase inhibitor activity | 1.34E-04 | 1.00E-01 | 4 |
| GO:0051346 Negative regulation of hydrolase activity | 1.44E-04 | 1.00E-01 | 14 |
| GO:0010951 Negative regulation of endopeptidase activity | 2.56E-04 | 1.00E-01 | 11 |
| GO:0010466 Negative regulation of peptidase activity | 2.78E-04 | 1.00E-01 | 11 |
| GO:0010948 Negative regulation of cell cycle process | 4.53E-04 | 1.00E-01 | 34 |
| GO:0046906 Tetrapyrrole binding | 5.20E-04 | 1.00E-01 | 4 |
| GO:0016705 Oxidoreductase activity, acting on paired donors, with incorporation or reduction of molecular oxygen | 5.34E-04 | 1.00E-01 | 10 |
| GO:0020037 Heme binding | 5.39E-04 | 1.00E-01 | 4 |
